# CellBouncer, A Unified Toolkit for Single-Cell Demultiplexing and Ambient RNA Analysis, Reveals Hominid Mitochondrial Incompatibilities

**DOI:** 10.1101/2025.03.23.644821

**Authors:** Nathan K. Schaefer, Bryan J. Pavlovic, Alex A. Pollen

**Author notes:** Correspondence (N.K.S), (A.A.P.).

## Abstract

Pooled processing, in which cells from multiple sources are cultured or captured together, is an increasingly popular strategy for droplet-based single cell sequencing studies. This design allows efficient scaling of experiments, isolation of cell-intrinsic differences, and mitigation of batch effects. We present CellBouncer, a computational toolkit for demultiplexing and analyzing single-cell sequencing data from pooled experiments. We demonstrate that CellBouncer can separate and quantify multi-species and multi-individual cell mixtures, identify unknown mitochondrial haplotypes in cells, assign treatments from lipid-conjugated barcodes or CRISPR sgRNAs, and infer pool composition, outperforming existing methods. We also introduce methods to quantify ambient RNA contamination per cell, infer individual donors’ contributions to the ambient RNA pool, and determine a consensus doublet rate harmonized across data types. Applying these tools to tetraploid composite cells, we identify a competitive advantage of human over chimpanzee mitochondria across 10 cell fusion lines and provide evidence for inter-mitochondrial incompatibility and mito-nuclear incompatibility between species.

## Background

Pooled processing of tissue or cells from many donors reduces batch effects from dissociation and cell capture, while increasing biological replication and experimental throughput^1–3^. Combining pooled processing with single cell genomics increases the scale of experiments and provides new resolution for large-scale studies of disease consequences^2,4^. Similarly, the “village-in-a-dish” approach, in which many cell lines are cultured together, can capture effects of natural genetic variation while avoiding difficulties inherent in scaling arrayed culture experiments^5–9^. Pooled culture experiments can also be used to isolate cell-extrinsic from cell-intrinsic phenotypic variation^10^, discover variation in stimulus-dependent responses^11^,and measure the consequences of epistatic interaction among groups of small-effect mutations^12^. While bringing together single cell genomics and population genetics^13^, pooled experiments pose new computational challenges. Bioinformatic tools are required to assign cells to individual-of-origin and perform quality control. Some demultiplexing tools exist, which can label cells by donor or cell line-of-origin using preexisting genotype data^14–16^ or without genotype data^15,17–19^ and assign cells to treatment groups using lipid-conjugated barcodes^20–26^. These methods can also identify droplets containing heterotypic cell doublets, upstream of methods that use gene expression signatures to identify doublets. Additionally, methods exist to quantify and remove the effects of ambient RNA contamination^27–29^ by inferring a gene expression signature in ambient RNA using information in empty droplets and/or heterogeneity across transcriptomic clusters.

There are several limitations of these existing tools, however. First, the common practice of aligning mixed-species data to composite reference genomes^30^ is memory-intensive, scales poorly with species number, and lacks statistical underpinning. Methods for inferring the individual-of-origin for each cell, especially without genotype data, lack a technique to independently verify results. Methods that infer individual-of-origin using genotype data can be slow to run, and assignments are sensitive to ambient RNA^30,31^. Many methods that assign custom labels to cells using lipid-conjugated barcode sequencing^24^ or assign CRISPR perturbations using sgRNA capture sequencing^32^ rely on fitting simple mixture models to barcode counts, and therefore require low background noise and high sequencing depth. Methods that identify cell doublets can only report specific doublet types rather than a consensus doublet rate. Finally, methods that detect ambient RNA contamination from gene expression data cannot identify the donors or cell lines driving contamination and can struggle to remove contamination in the dominant cell type, which can result in misinterpretation of cell types and differential expression^33^

We present CellBouncer, a computational toolkit that rectifies these shortcomings, improves speed and accuracy over existing tools, and provides novel functionality (Figure 1). For an improved user experience, CellBouncer programs have minimal dependencies and function as integrated programs rather than software pipelines. CellBouncer sets a new benchmark in accuracy, matching or exceeding existing tools where available, while delivering substantial performance gains—especially in handling large variant sets, genotype-free assignments, and high ambient RNA. Additionally, CellBouncer introduces new functionalities, including the ability to determine genotypic composition of ambient RNA, making it possible to identify and correct for its effects in ways that were previously not feasible. Using polymorphisms as a ground truth for ambient RNA contamination, we further explore the nature of ambient RNA sources at the transcript and cell compartment level. Finally, we apply a combination of CellBouncer tools to study mitochondrial competition in inter- and intra-species tetraploid cell fusions we generated from human, chimpanzee, and bonobo iPSCs, highlighting a possible mitochondrial-nuclear hybrid incompatibility separating the *Homo* and *Pan* lineages and clues to underlying mechanisms.

**Figure 1.**
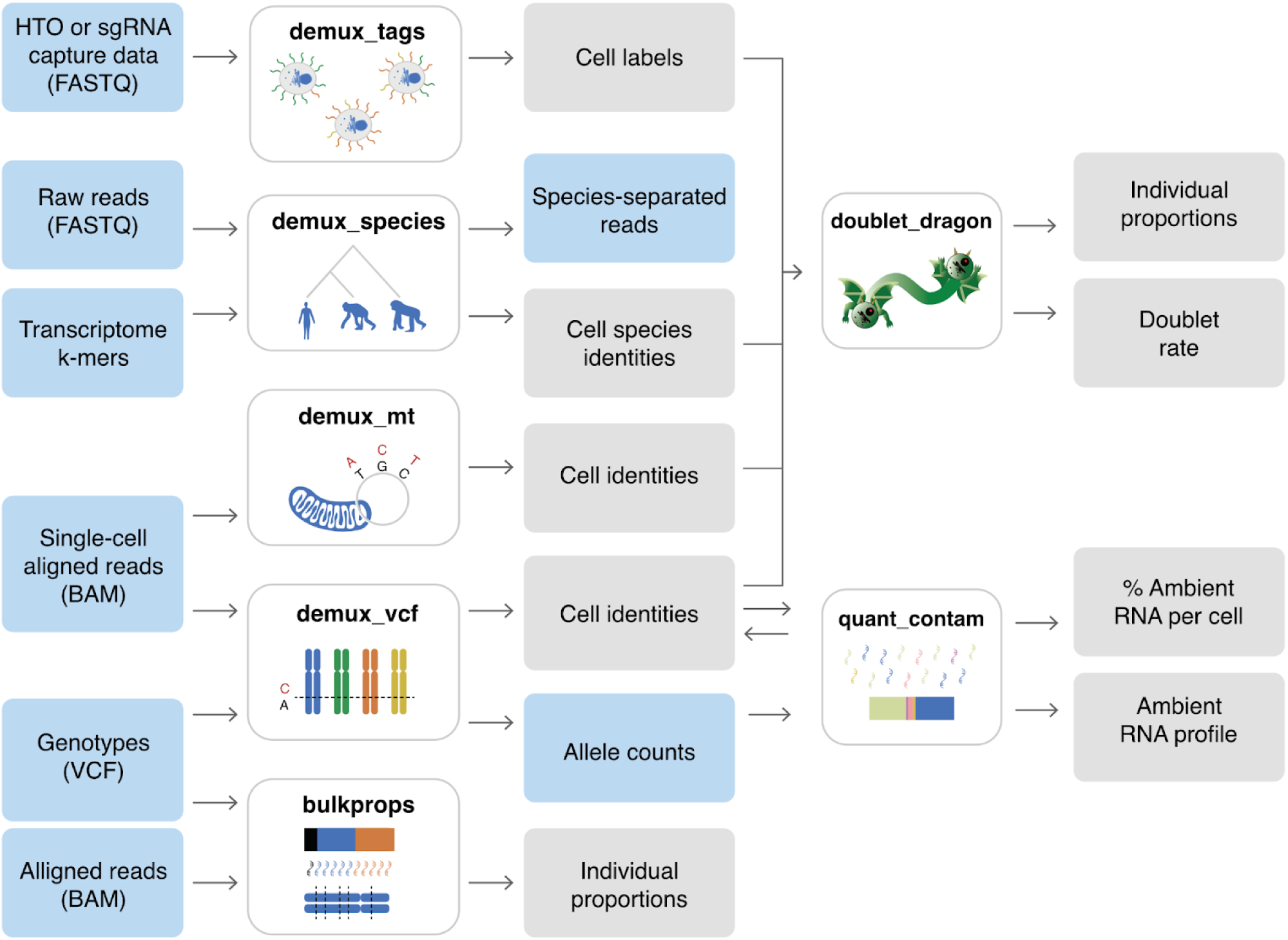
Overview of CellBouncer, a unified toolkit for demultiplexing and ambient RNA profiling of single cell sequencing data. Blue rectangles are input data, gray rectangles are output data and results. White rectangles with bold text are CellBouncer programs. The ambient RNA inference tool quant_contam and the doublet rate inference tool doublet_dragon use output from other CellBouncer tools as input.

### Species demultiplexing

Aside from “barnyard” experiments where cells from two species are mixed with the goal of calibrating measurements, single cell sequencing experiments that pool together cells from multiple species are rare, despite their utility for comparative studies^34–36^. Currently, no dedicated tools exist for separating cells by species of origin: standard practice is to use a composite reference genome and assign cells to the species to which the most reads align, after applying a cutoff^37^. This technique lacks statistical rigor, can become prohibitively memory intensive and cannot systematically identify inter-species doublets.

CellBouncer’s species demultiplexing algorithm, demux_species, counts UMI-collapsed short subsequences (k-mers) unique to each species’ transcriptome in reads from each cell, fitting a multinomial mixture model to assign each cell to its likeliest species (or doublet type)-of-origin and report the log likelihood ratio of the likeliest to second likeliest assignment (henceforth assignment LLR) (Figure 2A, Supplementary Methods). In doing so, it avoids the need for reference genome alignment or arbitrary cutoffs and identifies inter-species doublet barcodes. Demux_species also estimates which droplet barcodes are cell-positive, allowing an early estimate of the species proportions of true cells, and separates reads by species for downstream applications (Supplementary Methods).

**Figure 2.**
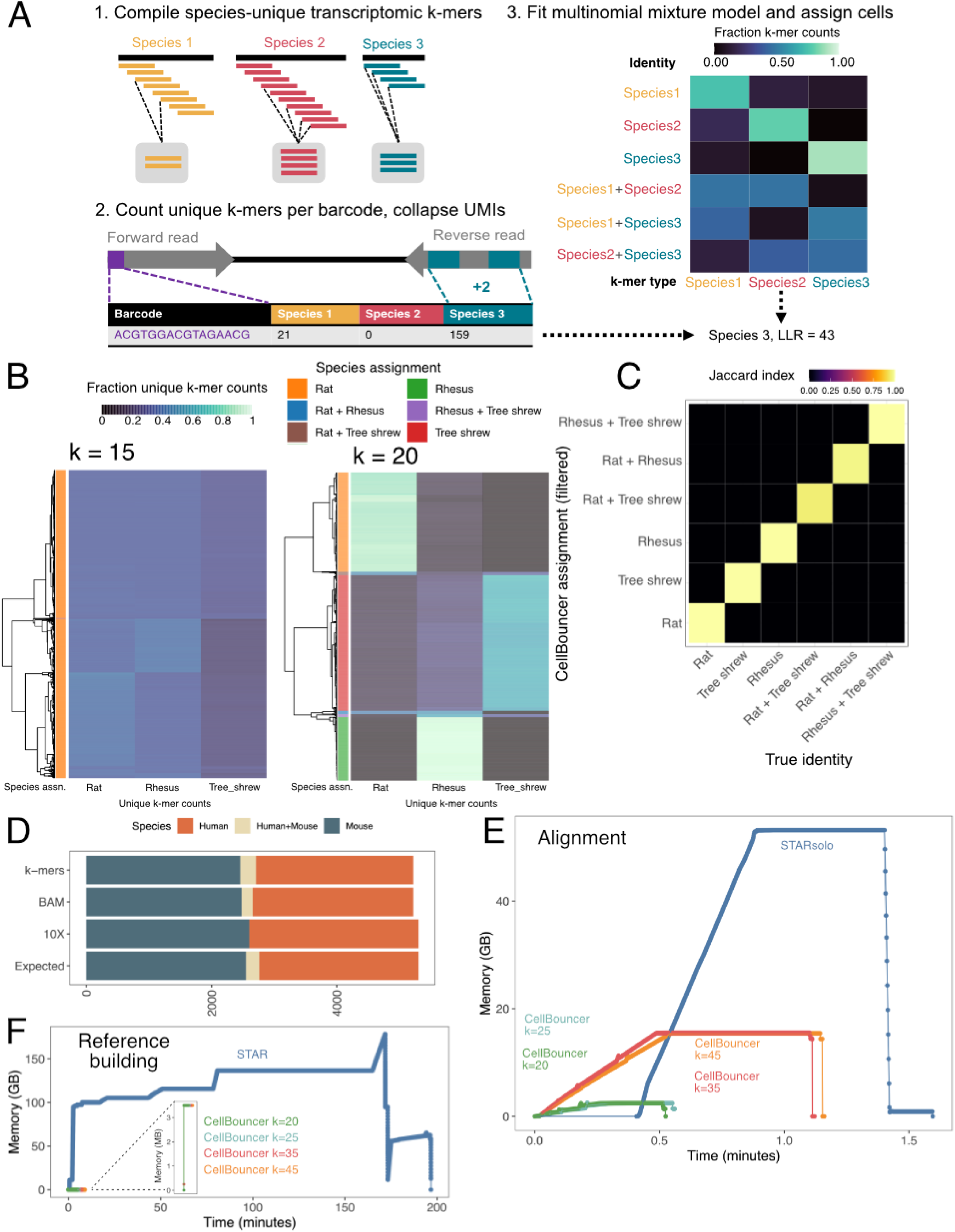
Rapid alignment-free species demultiplexing by species-unique k-mers. **A**: Schematic of demux_species algorithm. **B**: Species-specific k-mer counts in cell barcodes from labeled data set (rows), hierarchically clustered and labeled with species assignment (colored bars on left). Left plot: using 15-mers; right plot: using 20-mers. **C**: Comparison of CellBouncer-filtered species assignments (Y axis) to true species identity (X axis) in labeled data set with k=20 and 10 million sampled k-mers per species. Colors show Jaccard index (measure of mutual overlap, with 1.0 indicating complete agreement). **D**: Number of cells from each species in an unlabeled data set. Targeted: expected counts before collecting data, assuming an even proportion of human and mouse, 5,000 recovered cells, and a doublet rate of 0.04 (obtained from the 10X Genomics website). Heuristic: species assignments released by 10X Genomics, using read counts on a composite reference and a cutoff proportion. BAM: CellBouncer-filtered assignments from demux_species, using as input read counts on the composite reference genome. K-mers: CellBouncer-filtered assignments from demux_species, using species-specific transcriptomic 20-mer counts, with 20 million k-mers sampled per species. **E**: RAM usage vs time for a series of CellBouncer runs on a subset of the unlabeled data set, compared to mapping the same data to a composite reference genome using starSolo^7^. CellBouncer runs include time spent counting k-mers and assigning cells to species, but not demultiplexing reads. **F**: RAM usage vs time counting k-mers and producing unique transcriptomic k-mer lists for CellBouncer versus building a STAR index over a composite human/mouse genome.

We evaluated demux_species using two datasets: sequence data from three nonhuman species processed separately^38^, which provides a ground truth and for which we created simulated doublets (henceforth “labeled” dataset), and from an even mixture of 5,000 human and mouse cells^39^ for which the expected pool composition is known but the identities of individual cells were unknown (henceforth “unlabeled” dataset). On the labeled data, insufficiently long k-mers resulted in poor accuracy (Figure 2B, Table S1), but 20-mers were a sufficient length that achieved 99.9% accuracy labeling cell barcodes in the publication’s filtered barcode list and 99.8% accuracy labeling droplets that CellBouncer inferred to contain real cells (Table S1, Figure 2C). Moreover, clustering and visualizing k-mer counts can help evaluate success in the absence of ground truth data (Figure 2B). Sampling as few as 10 million k-mers per species improved speed and still gave accurate results and identified all true cell-containing droplets (Table S1).

In the unlabeled data set, CellBouncer’s inferred species proportions agreed with expectations (Figure 2D; Two-tailed chi-squared test p = 0.22) as did the doublet rate inferred from species assignments (inferred: 4.5%, expected 4.0%).

Tracking memory usage, we found running demux_species on 1 million reads from the unlabeled data set, with k=20 and sampling 20 million k-mers per species, to require only 44% of the execution time and 4.6% of the memory used by STARsolo^40^ to align the same reads to a composite reference genome (Figure 2E). Longer k-mers may be required for more closely related species, but our data structure’s resource requirements are mostly consistent within 32-base increments, meaning that any choice of k up to 32 comes with little additional cost (Online Methods). Additionally, identifying species-specific transcriptomic k-mers required much less time and memory (peak memory usage 3.5 MB, 5.33 minutes) than building a two-species STAR reference (peak memory usage 178 GB, 3.28 hours) (Figure 2F). This highlights that CellBouncer’s species demultiplexing program presents improvements in both speed and functionality over the current standard.

### Demultiplexing using SNPs

When cells from multiple individuals are pooled together, methods exist to assign cells to individual-of-origin using naturally occurring sequence polymorphisms, either with^14–16^ or without^15,17–19^ prior knowledge of genotypes. These methods eliminate the need to label cells manually (*e.g.* using cell hashing oligos^20^) and enable pooling at earlier stages of processing. Existing genotype-based demultiplexing tools are beset by long run times and reduced accuracy in the face of ambient RNA^31^, while genotype-free methods often require prior knowledge of the number of individuals in the pool and lack methods for validating results.

We have developed two tools to address these shortcomings: demux_vcf, a fast genotype-based individual demultiplexing program (Figure 3A), and demux_mt, a program that can simultaneously infer the number of individuals in a pool and assign cells to individual-of-origin by clustering mitochondrial haplotypes (Figure 3B). These tools are complementary: demux_mt can be used with a variant caller to identify individuals and their nuclear genome variants, which can then be used to more robustly assign cells to individuals using demux_vcf (Online Methods). Their results can also be compared for quality control.

**Figure 3.**
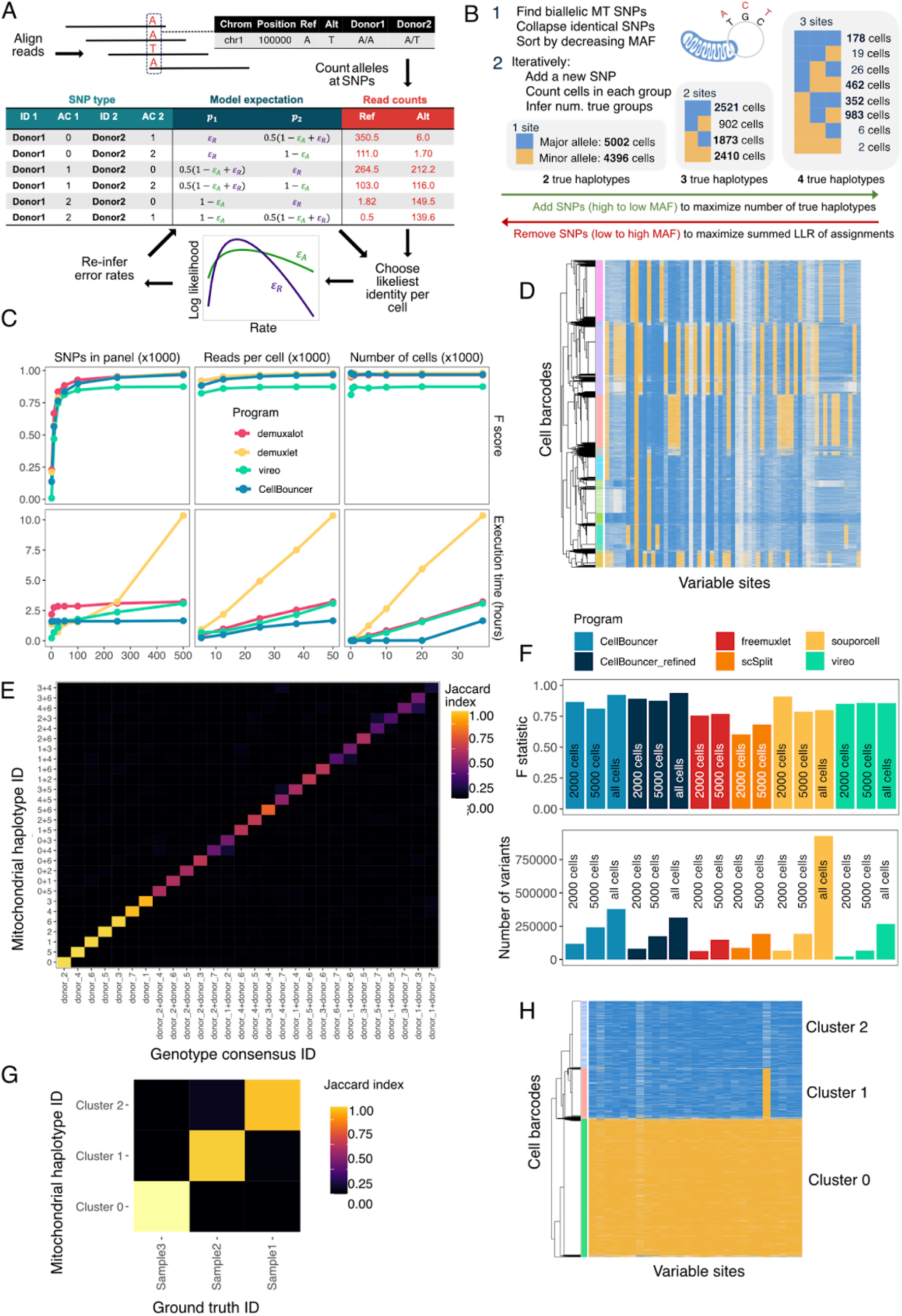
Rapid genotype-based and genotype-free mitochondrial demultiplexing tools in CellBouncer. **A**: schematic of demux_vcf, which demultiplexes cells using preexisting genotype data. “AC” in table headings means alternate allele count (1 indicates heterozygous genotype). **B**: schematic of demux_mt, which demultiplexes cells without genotype data by clustering mitochondrial haplotypes. **C**: Accuracy measured by F score and execution time (bottom row) of genotype-based demultiplexing tools on a test data set^41^, downsampled in various ways. All x-axis values are in thousands; reads per cell is an expectation based on 50,000 targeted reads per cell in the complete data set. **D**: Hierarchical clustering of inferred mitochondrial haplotypes on full data set shown in B (rows are cells, columns are SNPs, blue = major and yellow = minor allele). Colored bars on the left indicate demux_mt cluster assignment. **E**: consensus cell identities from genotype and cell hashing-based assignments (X-axis) vs identities assigned by demux_vcf (Y-axis). Doublets are two IDs separated by “+”. **F**: Comparison of genotype-free demultiplexing tools. Two downsampled (2000 cells, 5000 cells) and the full (all cells) datasets are shown. CellBouncer = MT-to-VCF pipeline; CellBouncer_refined = MT-to-VCF pipeline with genotype refinement. Top row: F score, with consensus IDs as truth. Bottom row: output number of SNPs segregating among IDs. CellBouncer outperformed all other methods on the 5,000 cell and full datasets, and identified the most variants, except souporcell on the full data set. **G**: Ground truth IDs from separately-processed donors vs mitochondrial identities using a scATAC-seq data set^42^. **H**: Hierarchical clustering of mitochondrial haplotypes shown in G. Mitochondrial cluster labels shown in G and H are the same.

Demux_vcf groups SNPs by their allelic state in each pair of individuals, counts reference and alternate alleles in each cell across these groups, and identifies cells by comparing the likelihood of each pair of candidate identities (Figure 3A). The rationale behind this strategy is that different sets of SNPs are informative for distinguishing between different individuals; using all available SNPs for every comparison risks allowing genotyping error at fixed sites to influence decisions when there is deep population structure in the reference panel (Online Methods). To account for ambient RNA, demux_vcf also infers and incorporates the rates at which reads erroneously mismatch expected genotypes (Figure 3A).

Demux_mt is unique among programs that identify cells without prior knowledge of genotypes in that it identifies SNP sites and infers the number of pooled individuals while assigning cells to individuals of origin. In pooled culture studies, the latter property may be a useful way to account for sample dropouts and cell line contamination that is frequent in cell culture^43–45^. By limiting its scope to mitochondrial SNPs, demux_mt also avoids the need to consider heterozygosity and the challenges it poses for genotype clustering. Demux_mt employs a heuristic algorithm that iteratively grows candidate mitochondrial haplotypes by a single SNP, stopping when it has discovered all individuals in the pool (Figure 3B); its core assumption is that the counts of cells with each mitochondrial haplotype follow a single Poisson distribution (Supplementary Methods).

To benchmark CellBouncer, we obtained 10X Genomics single-cell RNA-seq data from non-small cell lung cancer dissociated tumor cells from seven donors, with 40,000 targeted cells and with donors of origin labeled using 10X Genomics CellPlex membrane-attached oligos^41^ (10X NSCLC data set). We called variants specific to each donor and assessed the execution time and accuracy (F-score) of demux_vcf, along with three alternative programs (demuxlet^14,41^, demuxalot^16^, and Vireo^15^). We then downsampled the data to test the effect of the size of the genotype reference panel, the number of cells, and the sequencing depth on each program. Because of noise inherent in the CellPlex data, we used as ground truth the most commonly-assigned identity per cell from all three programs plus CellPlex, choosing the CellPlex identity in case of ties (Online Methods). CellBouncer, demuxalot, and demuxlet converged to similar accuracy as SNP panel size and sequencing depth increased toward levels typical in experiments, and CellBouncer achieved the highest accuracy when there were few cells (Figure 3C, Table S2). Additionally, CellBouncer’s demux_vcf also runs faster than the alternatives (Figure 3C).

We then ran demux_mt on the full data set, recovering the 7 expected individuals. Visual comparison of demux_mt identities to hierarchically clustered mitochondrial haplotypes shows high concordance (Figure 3D). We also compared mitochondrial haplotype assignments to the ground truth identities, matching identities by choosing reciprocal maximum Jaccard indices (Online Methods). This resulted in 88.7% precision and 72.0% recall for mitochondrial labels (Figure 3E). The lower recall reflects the presence of unlabeled cells, due to the fact that high-quality droplets tend to contain a low fraction of mitochondrial reads^46^. To address this, we devised a pipeline to label reads by mitochondrial haplotype, find nuclear SNPs segregating between mitochondrially-identified individuals, and use demux_vcf to assign cells to individuals using these variants. This approach, referred to as the MT-to-VCF pipeline, improved both precision (to 92.5%) and recall (to 92.3%) on the same data (Figure 3F, Table S3, Figure S1). For further improvement, we created a tool, refine_vcf, that alters genotype calls to maximize the likelihood of the read counts given cell-individual assignments (Online Methods). Incorporating this tool modestly increased precision (to 93.9%) and recall (to 93.7%) (Figure 3F, Table S3, Figure S2).

We compared the MT-to-VCF pipeline, with and without genotype refinement, to other existing genotype-free identification methods: freemuxlet^19^, souporcell^17^, scSplit^18^, and Vireo^15^ (in genotype clustering mode). Two of these methods, scSplit and freemuxlet, failed to run on the full dataset (Online Methods). To make the data tractable for these tools, we randomly sampled cells to create two smaller datasets. CellBouncer with genotype refinement largely outperformed the other methods and identified the most genomic variants (Figure 3F, Table S3). The other programs made two common errors avoided by CellBouncer: misidentifying one true individual as multiple clusters (freemuxlet, scSplit, and souporcell, Figure S3, Figure S4, Figure S5, Table S3) and mislabeling doublet identities (scSplit and Vireo, Figure S4, Figure S6, Table S3).

To assess whether demux_mt could identify mitochondrial haplotypes from single-nucleus data, in which few mitochondrial reads are expected, we downloaded single-nucleus ATAC-seq data from blood cells from three donors that were processed separately,^42^ providing ground truth identities. We recovered all three haplotypes, and they corresponded to donors of origin with 97.0% precision and 88.7% recall (Figure 3G), the latter of which would improve using the MT-to-VCF pipeline. Mitochondrial SNP coverage in the ATAC data appeared more uniform than in RNA-seq (Figure 3G vs Figure 3D), showing that this type of data is well suited for our method. Overall, CellBouncer provides methods for demultiplexing individuals using genetic variation, either with or without genotype data, that meet or exceed the capabilities of existing tools, while increasing processing speed.

### Modeling ambient RNA contamination

Droplet-based single cell RNA-seq studies are prone to ambient RNA contamination introduced by the processing required to generate single cell or nucleus suspensions, which can affect cell demultiplexing^31^ and experimental results^33^ if not properly modeled. In addition, disproportionate contribution of some individuals to ambient RNA can mask biological differences when multiple experimental conditions are pooled. However, existing methods rely on empty droplets^27,29^ and/or expression heterogeneity ^27,28^, making them susceptible to errors in cell calling and cell type identification and limiting their use in studies of a single cell type. Additionally, no method exists to account for the relative contribution of distinct individuals to ambient RNA. We present a CellBouncer program, quant_contam, that makes use of the ground truth contained in external genotype data to infer ambient RNA contamination from the rate at which cells mismatch expected genotypes.

CellBouncer’s quant_contam program uses read counts and cell identifications from demux_vcf, modeling each cell’s allelic counts as a weighted mixture of those expected from the cell’s true identity and those expected from ambient RNA. Further, ambient RNA is modeled as a weighted mixture of RNA from all individuals in the pool (Supplementary Methods). Using this inferred ambient RNA profile, it estimates cell-specific contamination rates and then re-assigns cells to individuals of origin, given knowledge of ambient RNA. Quant_contam can then optionally adjust single-cell gene expression count data to remove the effect of ambient transcripts (Figure 4A).

**Figure 4.**
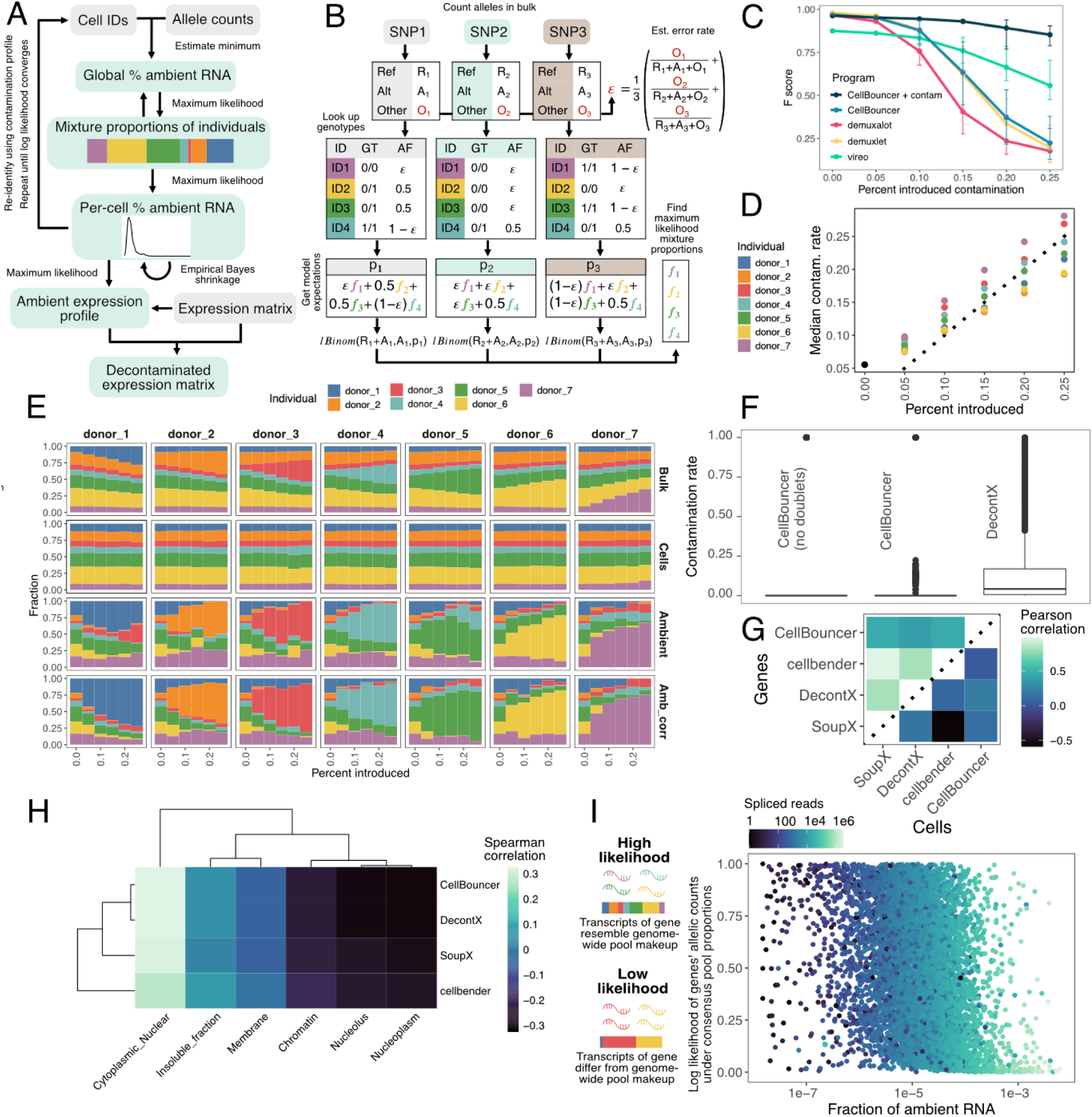
Ambient RNA profiling and bulk proportion inference using sequence variation. **A**: schematic of quant_contam, a program that uses output of demux_vcf to model ambient RNA as a mixture of individuals, infer ambient RNA per cell, and optionally infer the gene expression profile of ambient RNA and remove ambient-origin gene counts from a single cell expression matrix. **B**: schematic of bulkprops, a program that models a set of reads as a mixture of genotyped individuals without considering cell barcodes. Bulkprops only considers SNPs with no missing genotypes and infers the percent of reads originating from each individual via maximum likelihood. **C**: Accuracy (measured by F score) of CellBouncer, with and without ambient RNA modeling, compared to alternative programs, in identifying the true identity of cells in which a set amount of synthetic ambient RNA (X-axis) originating from a single individual was introduced to every cell in the data set. Data consists of 40,000 NSCLC cells from 7 donors^41^. **D**: After introducing a set amount of synthetic ambient RNA (X-axis) originating from a single individual (colored points) to every cell in a subsample of the NSCLC data set, the CellBouncer-inferred mean percent ambient RNA per cell is shown (Y-axis). The dotted line is y = x. **E**: Columns denote donors of origin for synthetic ambient RNA introduced to every cell in the data set, in increasing proportions (X-axis). Colors represent CellBouncer-inferred donor identities. Top row: inferred overall pool proportions from bulkprops. Second row: inferred cell identities. Third row: inferred ambient RNA profile. Fourth row: inferred ambient RNA profile if true cell identities are used (instead of CellBouncer-inferred identities that were less accurate due to contamination). **F**: Inferred mean contamination rate per cell using CellBouncer (with and without the doublet rate set to zero) versus DecontX on a SMARTseq2 data set, in which there is no true ambient RNA contamination. SoupX and cellbender, which require data from empty droplets that do not exist in this type of data, were not run. **G**: Pearson correlation coefficients between the gene expression profile (top/left half) and per-cell contamination rates (bottom/right half) inferred by each pair of ambient RNA inference tools. **H**: Spearman correlation coefficients between gene expression profiles inferred by ambient RNA inference tools (rows) and mean relative concentration index (RCI) reported by lncAtlas^47^ for transcripts of genes in different cell compartments. **I**: CellBouncer-inferred gene expression profile of ambient RNA (X-axis) versus the log likelihood of allele counts across individual genes, given the genome-wide maximum likelihood proportions of each donor, converted into percentiles (Y-axis). Points are colored by the total number of spliced reads from each gene.

We developed another program, bulkprops, that models total allele counts at each SNP as the sum of a weighted mixture of individual genotypes and outputs pool proportions; this is applicable to both bulk and single-cell data and can be applied to Census-seq experiments^48^ (Figure 4B). Through bootstrapping, both bulkprops and quant_contam can fit Dirichlet concentration parameters to pool proportions (Supplementary Methods), which allows for significance testing when comparing two sets of proportions. The combination of tools enables cross-checking single cell ambient RNA results with the overall pool proportions.

To test quant_contam and bulkprops, we created datasets with simulated ambient RNA of known quantity and origin. We chose 1000 random cells from the 10X Genomics NSCLC data set^41^ and used each of the 7 donors as a source of artificial contamination, replacing 5%, 10%, 15%, 20%, or 25% of each cell’s sequence data with reads sampled from the contaminant donor. We note that this is a likely more challenging test for cell identification than real ambient RNA, as contamination stemming from a single donor risks pushing programs to identify cells as doublet combinations of the true genotype and the contaminant individual.

We ran demux_vcf and quant_contam on each of these 35 datasets to identify individuals, profile ambient RNA, and refine cell-individual assignments by incorporating ambient RNA profiles. Although the accuracy of cell-individual assignments decreased as contamination increased, CellBouncer’s quant_contam-refined cell-individual assignments were significantly more accurate than those from other programs for high-contamination data (Figure 4C). CellBouncer quant_contam’s mean inferred contamination rates also agreed with introduced contamination rates (Figure 4D). Although cell identifications stayed mostly constant, ambient RNA profiles and bulk proportions changed in line with expectation (Figure 4E), with significant (p < 0.05) differences in the proportion of the contaminant individual observed in moderate-to-high contamination datasets (Table S4).

As a negative control, we obtained single cell RNA-seq data from multiple donors processed separately,^42^ prepared using SMART-seq2, a plate-based technology that does not produce ambient RNA^49^. After calling variants and running demux_vcf, quant_contam inferred a mean contamination rate of 0.48%. For comparison, we ran the ambient RNA inference tool DecontX^28^, which does not require the empty droplet information specific to droplet-based technology. This tool inferred 14% mean ambient RNA (Figure 4F), suggesting that CellBouncer may be less susceptible to false positive contamination inference, due to its use of genotype data as an external ground truth.

To measure consistency between ambient RNA inference tools, we ran quant_contam, cellbender^29^, DecontX, and soupX^27^ on the 10X Genomics NSCLC data set, computing Pearson’s r^2^ between each pair of program’s per-cell contamination rate inferences, and between each pair of programs’ ambient RNA gene expression profiles. Gene expression profiles were more consistent across programs than were per-cell ambient RNA estimates, with the highest r^2^ (0.980) between cellbender and SoupX, which both model ambient gene expression using empty droplets. Surprisingly, these programs’ per-cell contamination estimates were negatively correlated (r^2^ = -0.590). Conversely, the highest agreement between per-cell contamination rates was between CellBouncer and DecontX (r^2^ = 0.206), suggesting that ambient RNA inferences using genotypes at least partly agree with those made using expression patterns (Figure 4G). Ultimately, the lack of concordance between existing methods argues in favor of CellBouncer’s use of external ground truth data.

Ambient RNA has been found largely to consist of highly expressed cytoplasmic transcripts^33^, and we tested each program’s inferences for consistency with this observation. We correlated each gene’s inferred prevalence in ambient RNA according to each program with its relative concentration index (RCI) values from lncAtlas^47^, which measure transcript enrichment in subcellular locations; this comparison quantifies the extent to which ambient transcripts are associated with different cell compartments. Consistent with expectation, all programs’ ambient expression profiles correlated strongly with cytoplasmic versus nuclear RCIs, weakly with insoluble fraction versus other cytoplasm, and strongly negatively with nuclear compartments. CellBouncer and DecontX showed this pattern most starkly (Figure 4H), also showing the highest correlations with another feature associated with cytoplasmic transcripts, spliced transcript counts (Spearman’s rho for cellbender = 0.865, CellBouncer = 0.886, DecontX = 0.887, SoupX = 0.862, CellBouncer: Figure 4I). Further, we found significant agreement between ambient expression profiles and a published list of genes enriched for ambient RNA in brain tissue^33^ (Supplementary Text), suggesting that genic sources of ambient transcripts are cell type-independent.

Finally, we tested our model’s assumption that the allelic makeup of ambient RNA can differ from the bulk pool composition. To this end, we computed the log likelihood of allele counts for each gene, given the genome-wide consensus proportions of individuals in the pool (Supplementary Methods). A low log likelihood for a gene indicates that its mixture of genotypes differs from the genome-wide norm, suggesting either allele-biased expression or ambient RNA. Supporting ambient RNA as the cause of this mismatching, we found these low log likelihood genes are more likely to be spliced (Figure 4I), consistent with cytoplasmic contamination. Each program’s vector of genes contributing to ambient RNA correlated significantly negatively with gene log likelihoods, indicating that the origins of ambient transcripts are likely biased toward particular donors; CellBouncer and DecontX show this pattern most (Spearman’s rho for cellbender = -0.264, CellBouncer = -0.281, DecontX = -0.282, SoupX = -0.273) (Figure 4I).

Collectively, ambient RNA detection using genetic variation as a ground truth improves individual demultiplexing, recovers the contribution of distinct individuals to the ambient RNA pool, and further informs the nature and subcellular origin of ambient transcripts.

### Processing custom tags

Several technologies exist to associate custom barcode sequences with cells; they involve sequencing molecules that link custom tags to cell barcodes and computationally assigning user-defined labels to cells. These technologies can also detect heterotypic doublets and even distinguish cell-containing from empty droplets^20,50^. Clustered regularly interspaced short palindromic repeat (CRISPR) single guide RNA (sgRNA) capture^32^ produces similar data, excepting that cells may receive multiple sgRNAs, especially in high-multiplicity-of-infection (MOI) screens, and the presence of two labels in a cell can therefore not be used to identify cell doublets. We will refer to custom barcode-derived cell labels of all types, together with captured sgRNA sequences, as “tags.”

Existing methods for assigning cells to tag-based identities can err when data are sparse^51^, and others do not sufficiently account for contamination from ambient tags when making assignments. To address these issues, we introduce demux_tags, which models tag-based identities as multinomial distributions over each tag’s counts, informed by total tag depth per cell. Demux_tags learns mean “foreground” and “background” tag counts by fitting a 2-component mixture model to each tag independently. It then simulates cells with each possible identity by sampling random foreground or background tag counts corresponding to the identities, and fits multinomial linear models to predict the expected fraction of each tag type in a cell with a given identity, conditional on the total “foreground” tag count in the cell. This allows demux_tags to build a multinomial model for each possible identity for every cell, and then choose the likeliest model (Figure 5A). Demux_tags makes several simplifications to speed up processing of sgRNA count data, in which there are more identities to consider (Supplementary Methods).

**Figure 5.**
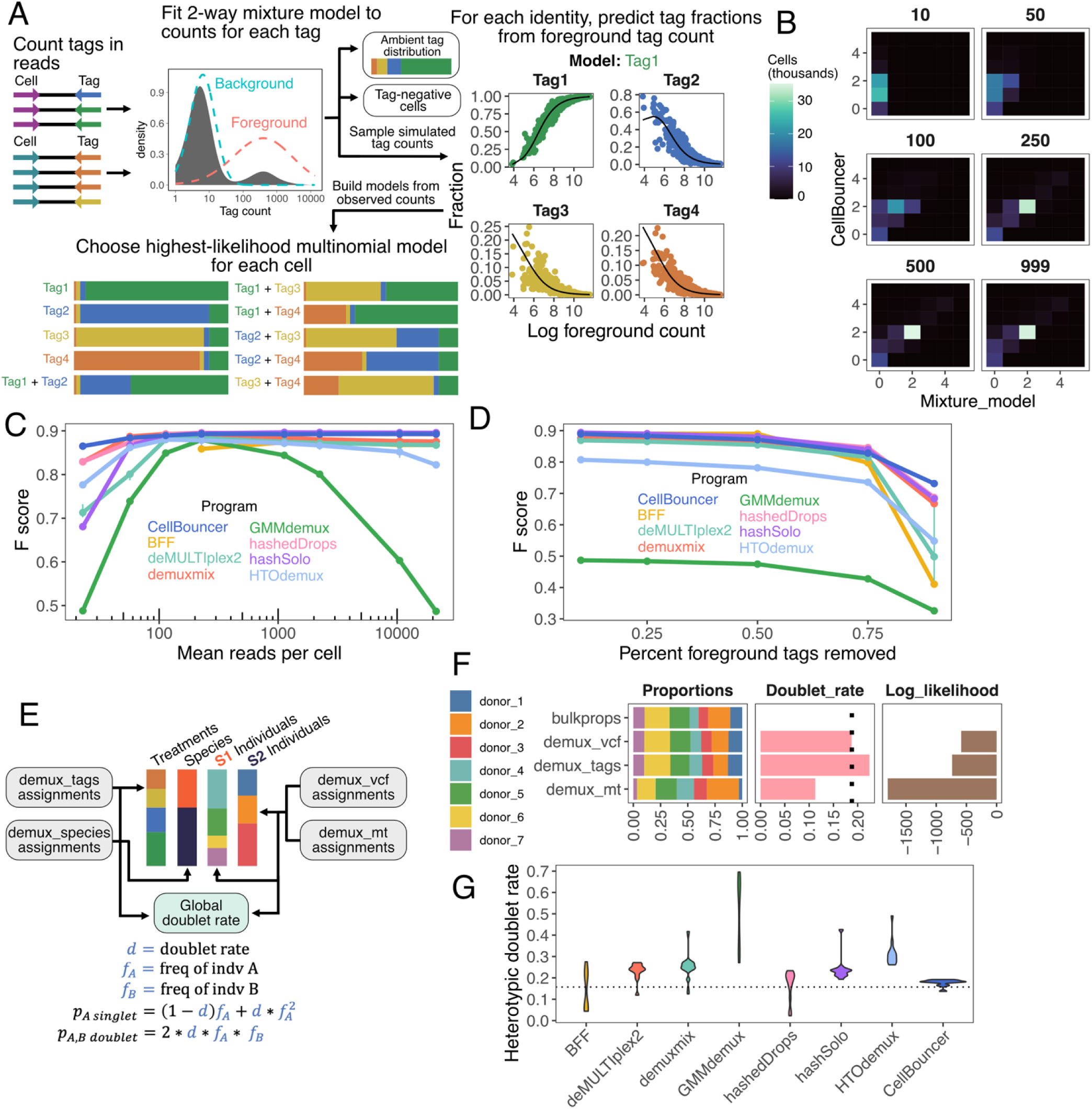
Tools for assigning custom label-based identities (or guide RNAs) to cells, and for inferring a consensus doublet rate and pool proportions from one or more CellBouncer programs. **A**: schematic of demux_tags. **B**: Using data from a published CRISPR screen that targeted 2 sgRNAs per cell^32^, box headings show the number of expected sgRNAs per cell after downsampling the data to a range of tag read counts, the X-axis shows the assigned number of sgRNAs per cell using the source publication’s sgRNA assignment strategy, and the Y-axis shows the number using demux_tags, and the color denotes the number of cells receiving the indicated number of sgRNAs from both strategies. We excluded counts of cells that received 5 or more sgRNAs, which correspond to the 99.9^th^ percentile of all cells. **C**: Downsampled number of 10X Genomics CellPlex reads per cell versus F score (harmonic mean of precision and recall) of CellPlex-based identities inferred by programs shown. For true cell identities, we used the consensus of all VCF-based tools examined in this study, using 10X-provided CellPlex identities to break ties. Values shown are means of three trials; error bars when visible show minimum and maximum values. Missing BFF points indicate failed runs. **D**: similar to C, but in each trial, the indicated fraction of counts of the tag representing each cell’s true identity were removed, in order to reduce the signal to noise ratio. **E**: Schematic of doublet_dragon. The program determines which input cell labels correspond to the same groups of labels (*e.g.* species, treatments, or individuals) and infers a consensus set of label proportions per label group, along with a global doublet rate that includes homotypic doublets. S1 individuals and S2 individuals refer to individuals belonging to species 1 and species 2, after species demultiplexing. **F**: Pool proportions, doublet rate, and log likelihood of cell identity counts inferred by doublet_dragon after running three CellBouncer programs on different aspects of the 10X NSCLC data set^41^. Maximum likelihood pool proportions from bulkprops are also shown for comparison. **G**: Heterotypic doublet rates inferred by different tag assignment programs, along with the maximum likelihood consensus heterotypic doublet rate (dotted line), from running doublet_dragon on output from all VCF-based demultiplexing tools examined in this study. Data shown are all downsampled datasets (three per X axis value) from C and D.

We tested the effect of low sequencing depth on demux_tags’s sgRNA assignments by downsampling published sgRNA capture data^32^ to various expected guide counts per cell. For each downsampled set of sgRNA counts, we compared the number of sgRNAs assigned to each cell by demux_tags, and by the original mixture model strategy (Online Methods), to the study’s target of two sgRNAs per cell. Demux_tags assigned two sgRNAs per cell more often than the mixture model on the full data set and at all levels of downsampling (Figure 5B), with a mode of two assigned sgRNAs per cell even with only 50 mean sgRNA counts per cell (5% of the data).

We then tested demux_tags against alternative programs for assigning cell labels from custom barcode tags, with different levels of sequencing depth. We obtained 10X CellPlex reads from the 10X NSCLC data set^41^, which were used to label cells’ donors of origin. We then counted CellPlex tags using demux_tags and then repeated this process on three random samples each of 0.1%, 0.25%, 0.5%, 1%, 5%, 10%, and 50% of the reads. Next, we used demux_tags, BFF^21^, deMULTIplex2^22^, demuxmix^23^, GMMdemux^24^, hashedDrops^26^, hashSolo^25^, and HTOdemux^20^ to assign identities to cells from each set of counts, measuring accuracy (F-score) using ground truth labels derived from the majority vote (with no ties) of all tested genotype-demultiplexing programs run on the same cells. Some tools, such as GMMdemux and HTOdemux, over-identified doublets as the number of tags per cell passed an optimum, but most programs performed well with 100 or more tag counts per cell (Figure 5C). CellBouncer, deMULTIplex2, hashedDrops, and hashSolo performed best; CellBouncer scored highest on the sparsest data (F 3.57% higher than the next best) and was within 0.44% of the top scorer on all other datasets (Figure 5C, Table S5).

To model experiments with “leaky” labeling leading to high background tag counts, we used the same data to simulate lower signal-to-noise ratio by removing 10%, 25%, 50%, 75%, or 90% of counts belonging to tags representing each cell’s true identity. CellBouncer performed best on the noisiest data set (4.4% higher than the next alternative) and comparably to the top method (within 1.8% of the highest F) for other datasets (Figure 5D, Table S6).

### Inferring the consensus doublet rate

To aid in quality control, we developed a tool, doublet_dragon, that uses sets of cell identities from other CellBouncer programs to estimate consensus pool proportions and a global doublet rate that encompasses homotypic as well as heterotypic doublets. Doublet_dragon can take a single or multiple sets of inferred identities for the same cells, which can overlap in meaning (e.g. genotype and mitochondrial identification of donors) or not (e.g. donor identities and treatment conditions). Doublet_dragon models the number of cells with each identity as a sample from a binomial distribution, with the probability of an identity determined by its prevalence in the pool and a global doublet rate (Supplementary Methods, Figure 5E). After inferring these parameters by maximum likelihood, it computes the log likelihood of each set of cell identities under the consensus model. This can be used to compare the relative confidence of each set of identities.

We used doublet_dragon to estimate the doublet rate in the NSCLC data set, using the genotype-based IDs from demux_vcf, demuxlet, Vireo, and demuxalot, at 18.8%, close to the expected rate of 16% given 40,000 recovered cells. We then compared doublet_dragon inferences from CellBouncer’s genotype-, mitochondria, and tag-based identities for these cells, using 18.8% as the expected doublet rate and bulkprops inferences as expected pool proportions. Genotype-based identities appear most accurate, followed by tag-based, and then mitochondrial identities (Figure 5F). This reflects the amount of data used for each set of identifications: read counts at hundreds of thousands of SNPs, tens of thousands of tag counts, or read counts at a handful of mitochondrial sites. Finally, we computed a heterotypic doublet rate from each tag-assignment program’s output on each set of tag counts we produced; estimates from demux_tags were more consistent and closer to expectation than those from other programs (Supplementary Text, Figure 5G).

Together, demux_tags improves sgRNA and other tag-based assignments, particularly in cases of low read counts or low signal to noise ratios from leaky barcodes, while doublet_dragon can use all available demultiplexing data to assess their relative quality and infer a global doublet rate, to aid in quality control and inform downstream analyses.

### Investigating mitochondrial inheritance and gene regulation in inter-species composite cells

We next applied CellBouncer to address an open biological question about nuclear-mitochondrial interactions among hominids. Hominid cell fusions bring two species’ genomes into a shared regulatory environment^53–56^, allowing the study of *cis* versus *trans* gene regulatory divergence and nuclear-mitochondrial interactions. Previous studies have hinted at human-biased mitochondrial inheritance relative to chimpanzees, but these findings were limited by a small number of contributor pairs and did not explore the extent of this bias, the effects at the single cell level, or causal mechanisms^53^. To systematically investigate this phenomenon, we generated 24 polyclonal intra- and inter-species tetraploid composite induced pluripotent stem cell (iPSC) lines^57^ from 12 hominids (Online Methods). We then collected single-cell RNA-seq to explore whether mitochondria from one or both contributor cell lines persist in composite cells (Figure 6A).

**Figure 6.**
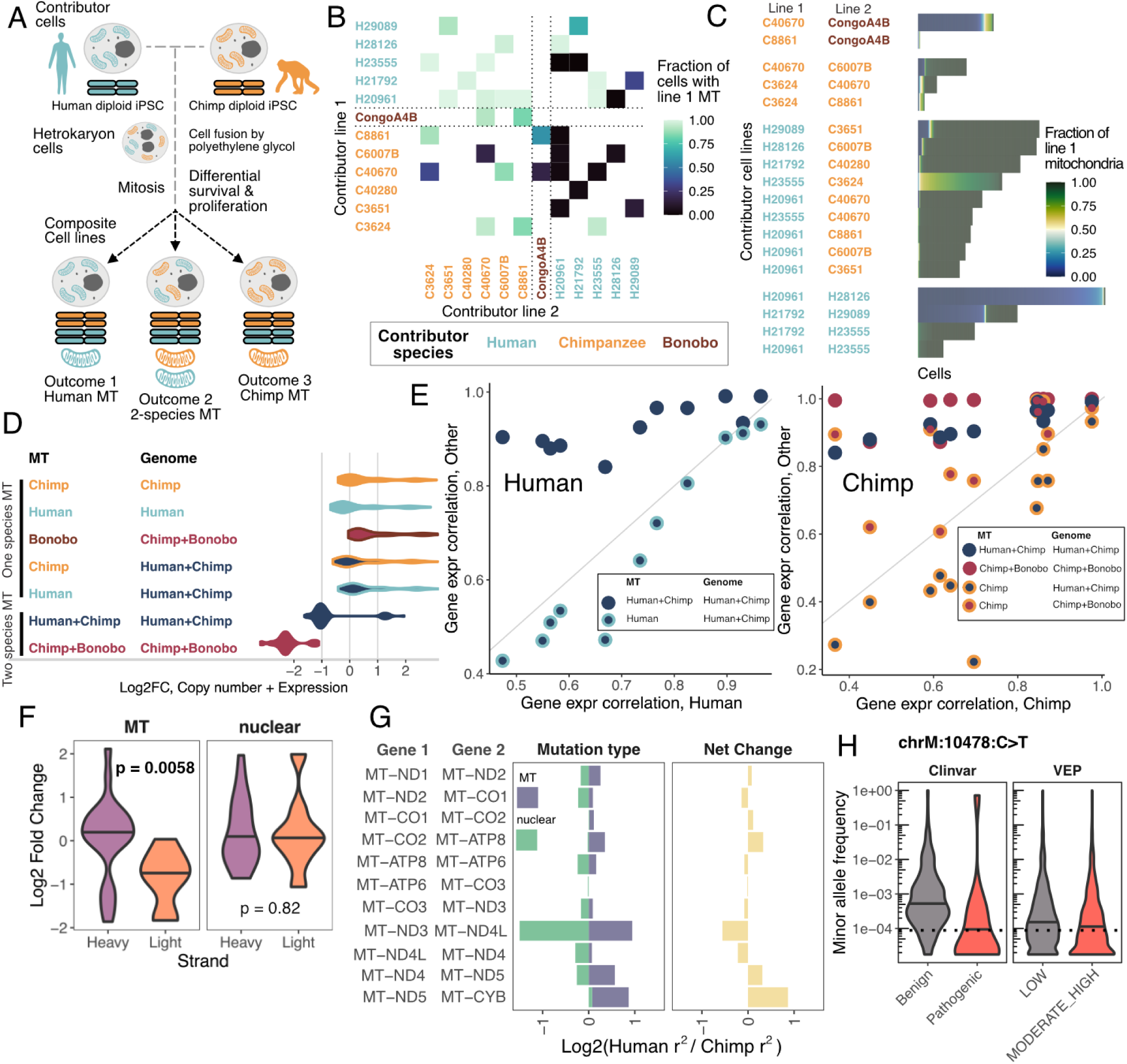
Investigation of mitochondrial inheritance and gene regulation in inter-species composite cell lines. **A**: construction of composite cell lines and possible mitochondrial inheritance outcomes. **B**: For each pair of contributor cell lines (X and Y axes) that were fused to form composite cell lines, the fraction of composite cells containing mitochondria from the contributor line on the Y axis is shown. A fraction of 1.0 for both iPSC lines indicates that cells all received mitochondria from both contributor lines. **C**: data from B, but showing individual cells’ fractions of mitochondrial reads stemming from contributor lines. **D**: For protein coding mitochondrial genes, log_2_ fold changes due to mitochondrial copy number and expression level (and excluding post-processing) are shown. Distributions encompass all mitochondrial genes and all comparisons of the given mitochondrial/genomic identity shown against all others. Only cell classes for which there were at least 10 cells with at least 100 informative allelic mitochondrial reads are shown (Chimp+Bonobo with Chimp MT is excluded). **E**: Mitochondrial post-processing differences, measured as correlation between adjacent pairs of protein-coding genes on the heavy strand. Each point shows Pearson’s r^2^ between gene pairs in cells of the identity given in the legend (Y axis) against single-species cells, where single-species cells in the left panel are composites of two human cell lines and single-species cells in the right panel are composites of two chimpanzee cell lines. **F**: Log_2_ human vs chimpanzee fold change in expression of all mitochondrial genes by strand, decomposed into mitochondrial and nuclear changes (where MT refers both to proximal and distal mutations on the mitochondrial genome). **G**: Log_2_ human vs chimpanzee fold change in post-processing, measured as Pearson’s r^2^ between adjacent pairs of heavy strand protein-coding genes, decomposed into mitochondrial and nuclear changes (left panel) and showing the net change (right panel). **H**: GnomAD^52^ minor allele frequency (MAF) of a back-mutation at a fixed, human-derived allele at a candidate cleavage site. Distributions show MAFs of other mitochondrial mutations with the functional (VEP) and pathogenic (ClinVar) annotations shown.

Addressing this biological question in single cells presented particularly challenging demultiplexing problems of identifying the combination of individuals-of-origin for each cell, confirming results with tags, profiling ambient RNA, and determining the combination and fraction of mitochondrial haplotypes in each cell. After collecting single-cell RNA-seq data, we assigned cell line identities using demux_tags on MULTI-seq labeling data^50^ and demux_vcf with genomic variant data generated from high-coverage whole genome shotgun sequencing. Treating MULTI-seq labels as a ground truth for cell line identity, we found that demux_vcf more accurately identified composite cell lines of origin than alternatives (Table S7) and that quant_contam produced the best ambient RNA inferences (Figure S7, Supplementary Text).

We then identified mitochondrial haplotypes present in each cell (Supplementary Text, Online Methods, Figure 6B,C). We found that inter-species composite cells tend to retain mitochondria from only one species, with chimpanzee/bonobo cells biased toward bonobo mitochondria, and human/chimpanzee cells biased toward human mitochondria. There were many exceptions, however: 2.5% of human/chimpanzee cells had only chimpanzee mitochondria, and 13.9% had mitochondria from both species; 8.0% of chimpanzee/bonobo cells had only chimpanzee mitochondria and 9.9% had mitochondria from both species. Of the 10 human/chimpanzee composite cell lines, only one retained both species’ mitochondria in most of its cells (Figure 6B,C), suggesting a contributor-specific influence on this outcome. These findings were robust to false-discovery rates observed for within species cell fusions (Online methods).

We also observed species differences in mitochondrial inheritance for within-species composites: compared to human/human lines, chimpanzee/chimpanzee lines more frequently retained the mitochondria from both contributors, albeit in unequal ratios (Figure 6 B,C), and this observation was robust to species-specific differences in mitochondrial diversity (Figure S13). Studies of mitochondrial heteroplasmy in mice and humans have shown that cells can selectively promote certain mitochondrial haplotypes over others in a tissue- and age-dependent manner^58,59^; If the efficiency or dynamics of this process differ across species, particularly in stem cells, it could help explain the interspecies patterns observed in our data.

The presence of cells with all combinations of mitochondrial inheritance patterns allowed us to examine the influence of hominid nuclear-mitochondrial interactions on gene expression. Mitochondrial genes are expressed as polycistronic transcripts, which are later cleaved and selectively degraded by post-processing machinery; it is therefore possible to isolate the effects of the combined influence of copy number and expression level from the effects of post-processing differences^60^. We computed overall differential expression (DE) across groups, reflecting aggregate differences in copy number, expression level, and post-processing. We then computed DE normalized to total mitochondrial gene expression rather than genome-wide expression, reflecting post-processing effects, and the differences between these two DE measures, reflecting DE due to copy number and expression level (Online Methods). Strikingly, cells with mitochondria from two species show decreased copy number and/or expression level across all mitochondrial genes when compared with all other types of cells (Figure 6D), suggesting gene-regulatory crosstalk between allospecific mitochondria.

Canonical mitochondrial mRNA post-processing involves cleavage at junctions where tRNA genes separate protein-coding sequences, with cuts occurring at the 5’ and 3’ ends of the tRNA genes^61^. We therefore sought to measure mitochondrial post-processing efficiency in different composite classes (species combinations) by calculating the correlation in expression between adjacent pairs of protein-coding genes on the mitochondrial heavy strand. We observed increased correlation at each junction in cells with mitochondria from two species relative to all other types of cells (Figure 6E), indicating lower levels of cleavage and degradation in composite cells of this class.

Comparing human/chimpanzee cells with both species’ mitochondria to those with only one, a significant proportion of DE genes are involved in apoptosis and cell cycle arrest (Table S8) and similar enrichments are seen when comparing chimpanzee/bonobo composite cells with both species’ mitochondria to those with only bonobo mitochondria (Table S9, S10), Additionally, these gene sets significantly overlap (Figure S14) and are enriched for protein-protein interactions (Figure S15), further suggesting incompatibility between allospecific mitochondria in the same cell. A prior study found that human cells cannot take up chimpanzee mitochondria and survive, even when human mitochondria are functionally impaired; full ablation of human mitochondria is required for survival^62^. This suggests the involvement of nuclear mechanisms. We found several candidate genes by testing the effects of contributor lines on gene expression in human/chimpanzee cells. H23555, the human contributor to the fusion with the highest survival of both species’ mitochondria, was associated with downregulation of several genes involved in p53 signaling (Figure S15), which had higher-than-normal connectivity in the interaction network (Figure S16): CDKN1A, GDF15, PHLDA3, and ZMAT3. This provides an exciting first glimpse at candidate mechanisms underlying this process.

We next used our data to disentangle regulatory divergence between human and chimpanzee mitochondrial genes stemming from mutations in mitochondrial DNA (henceforth “local mutations”) from those found in the nuclear genome (henceforth “nuclear mutations”; Online Methods). We reasoned that nuclear mutations can only contribute to divergence in mitochondria gene expression by acting in *trans*, while local mutations in mitochondria could act in *cis* or *trans*. We first found local downregulation of all genes on the light strand in humans relative to chimpanzees (Figure 6F), resulting in significantly altered MT-ND6 expression. Prior work has found species differences in light strand promoter usage, including a chimpanzee-specific transcription start site at an A/T rich locus in the MT-TE gene^63^, that could explain this difference.

Considering mitochondrial RNA post-processing efficiency, we observed large-effect, compensatory nuclear and mitochondrial species differences working in opposition at the MT-ND3/MT-ND4L locus, resulting in little net change in expression correlation (Figure 6G). These transcripts are partly cleaved by an unknown non-canonical mechanism that targets position chrM:10479^64^, which is immediately adjacent to a fixed, *Homo*-specific derived T-C mutation. Population sequencing data^52^ suggest that a C-T back-substitution at this site (rs1603222851), which results in a synonymous change in MT-ND4L, persists at a frequency lower than typical synonymous mitochondrial mutations and more characteristic of coding and disease-associated variants (Figure 6H). Additionally, RNA secondary structure modeling around this site suggests that the derived allele increases the probability of base pairing in a hairpin loop structure (Figure S17), potentially making the position less accessible to nucleases. Although beyond the scope of this study, this mutation may act together with a human-specific upregulation of nuclease activity to comprise a hybrid incompatibility between the *Homo* and *Pan* lineages. Similar compensatory mutations involving nuclear regulation of mitochondrial genes have been found in multiple models for early species divergence^65–67^ and may make up the bulk of Dobzhansky-Muller hybrid incompatibilities found in nature^68^. Further experiments using composite cell lines may uncover more examples of such otherwise-invisible regulatory changes.

The CellBouncer suite enabled this analysis by assigning cells to their composite lines of origin, identifying mitochondrial haplotypes, and quantifying ambient RNA, allowing us to profile many different composite cells. This represents an advance over both single-cell sequencing studies of individual cell lines, and bulk sequencing studies that cannot detect cell-specific differences in mitochondrial haplotype.

## Discussion

We introduce CellBouncer, a versatile toolkit for processing of pooled single cell genomics data. It supports flexible cell demultiplexing by species- and individual-of-origin, with or without genotype data or prior knowledge of the number of pooled individuals while modeling the extent and composition of ambient RNA to refine demultiplexing assignments and correct gene expression values. Its functionality extends to analyzing bulk sequencing data as a mixture of genotyped individuals, assigning treatments or sgRNAs to cells based on cell hashing or CRISPR guide capture data, and inferring a global doublet rate including homotypic doublets. We showed that CellBouncer outperforms existing methods where they apply, while introducing new capabilities beyond their scope. It improves computational performance (species and genotype-based individual demultiplexing), accuracy (noisy cell hashing data, low-depth CRISPR guide capture data, genotype-free individual demultiplexing, genotype-based demultiplexing with high ambient RNA), and adds novel functionality (ambient RNA profiling, doublet rate inference).

CellBouncer expands the possibilities for single cell study design, increasing throughput while reducing biases and batch effects. It enables species mixing experiments that were previously impractical due to the computational burden imposed by composite reference genomes for more than two or three species. By leveraging mitochondrial clustering, it provides a powerful alternative for demultiplexing individuals when genotype data are unavailable, as well as an independent validation method for other cell identification approaches. CellBouncer’s demux_vcf program scales efficiently to large datasets and SNP panels while accounting for population structure, even allowing for pooling of intra- and inter-species composite cell lines. The quant_contam module enhances pooled study integrity by profiling the origins of ambient RNA, flagging problematic donors or cell lines. Demux_tags enables accurate tag-based identity and CRISPR sgRNA assignments from noisy data, expanding the usable portion of datasets that might otherwise be discarded. Finally, doublet_dragon will enable users to calibrate cell loading density and improve doublet detection, guiding removal of heterotypic and homotypic doublets, with possible implications for compressed sensing studies^69^.

Given the risks of demultiplexing^31^ errors and biological misinterpretation^31,33^ due to ambient RNA contamination, CellBouncer’s genotype-based demultiplexing program and its accompanying ambient RNA inference tool offer a valuable advance over existing methods. Whereas existing methods require heterogeneity in gene expression or empty droplets to infer contamination, our method compares data to the external ground truth contained in genotype data, making it suitable for pooled cell data regardless of cell type diversity or empty droplets. Additionally, as current ambient RNA detection methods show limited agreement, the external ground truth of genotype data enables precise modeling and correction of ambient RNA contamination, improving both accuracy and interpretability.

Using CellBouncer tools in combination to process data from a mixture of intra- and inter-species composite iPSCs, we found that inter-species composite cells tend to favor one species’ mitochondria over the other, with human/chimpanzee cells preferring human mitochondria and chimpanzee/bonobo cells preferring bonobo mitochondria. However, a subset of cells deviated from this pattern, allowing us to explore the impact of each species’ mitochondrial and nuclear genomes on global gene expression. Cells retaining mitochondria from both species show signs of cell cycle arrest and apoptotic signaling, globally decreased mitochondrial gene expression and/or copy number, and impaired mitochondrial mRNA post-processing relative to cells with a single species’ mitochondria. These findings suggest that cross-species mitochondrial interactions disrupt normal gene expression regulation, triggering a nuclear apoptotic response . Our findings here agree with a prior study that discovered that human cells cannot take up great ape mitochondria unless human mitochondria are first eliminated^62^.

We also found evidence for opposing mitochondrial and *trans*-nuclear mutations altering post-processing of two neighboring mitochondrial genes. While more work remains to be done to elucidate this mechanism, the changes we found at these loci may be an early glance at an example of nuclear-mitochondrial hybrid incompatibility loci separating humans from our closest extant relatives.

## Acknowledgements

The authors thank Reed McMullen and Jenelle Wallace for feedback on the software and manuscript, Richard E. Green and Yoav Gilad for providing cell lines, and Helena Pinheiro for illustrations in Figure 1. Sequencing was performed at the UCSF CAT, supported by UCSF PBBR, RRP IMIA, and NIH 1S10OD028511-01 grants. This work was supported by the following funding sources: Weill Neurohub Fellowship to NKS, National Institutes of Health 1R01MH134981, DP2MH122400, National Science Foundation #2134955, the Schmidt Futures Foundation, and the Keck Foundation to AAP. This project was funded in part by the Emory National Primate Research Center Grant No. ORIP/OD P51OD011132. AAP is a New York Stem Cell Foundation Robertson Investigator.

## Contributions

N.K.S., B.J.P., and A.A.P conceived the project and experimental design. N.K.S., designed the algorithms and performed all analysis. B.J.P. designed the cell fusions and performed all experiments. A.A.P supervised the study and secured the funding. All authors discussed the results and contributed to the final manuscript.

## Ethics declarations

All studies were approved by UCSF GESCR (Gamete, Embryo, and Stem Cell Research) Committee.

## Code availability

The CellBouncer repository is available at https://github.com/nkschaefer/cellbouncer.

## External data

Species demultiplexing benchmarks used data from Gene Expression Omnibus (GEO) accession GSE178265 and 10X Genomics: https://www.10xgenomics.com/datasets/5k-hgmm-3p-nextgem.

Mitochondrial demultiplexing, genotype-based demultiplexing, tag-based demultiplexing, and ambient RNA benchmarks used data obtained from 10X Genomics: https://www.10xgenomics.com/datasets/40-k-mixture-of-nsclc-dt-cs-from-7-donors-3-ht-v-3-1-3-1-high-6-1-0.

Mitochondrial demultiplexing using ATAC data used data obtained from ArrayExpress ID E-MTAB-9068.

The negative control experiment for ambient RNA inference used data obtained from ArrayExpress ID E-MTAB-9067.

Data quantifying enrichment of transcripts in sub-cellular compartments were obtained from lncAtlas: https://lncatlas.crg.eu/.

CRISPR guide capture assignment benchmarking used data obtained from GEO accession GSE146194.

## Online Methods

For algorithms behind CellBouncer programs, see “Descriptions of algorithms” section in Supplementary Text.

### Species demultiplexing

#### Benchmarking

##### Labeled data set

We downloaded single-cell RNA-seq data from three species (*Macaca mulatta*, *Rattus norvegicus*, and *Tupaia belangeri*) produced as part of a prior study^38^ and processed separately, so that the true species identity of each cell was known. We obtained the data from NCBI GEO (ID GSE178265) and used reads from sample IDs SAMN19715216 (*R. norvegicus*), SAMN19715217 (*T. belangeri*), and SAMN19715221 (*M. mulatta*). To build reference lists of unique k-mers, we obtained reference transcriptomes from each species from Ensembl release 112: Rattus_norvegicus.mRatBN7.2.cds.all.fa, Tupaia_belangeri.TREESHREW.cds.all.fa, and Macaca_mulatta.Mmul_10.cds.all.fa.

For each k-mer size we tested (15, 20, 25, 35, and 45), we counted k-mers in each transcriptome using FastK^70^ with the -t option set to 1. We then used the CellBouncer program get_unique_kmers to iterate through each transcriptome in sorted order, identify k-mers present in only one transcriptome, and then apply the DUST algorithm^70,71^ to each unique k-mer, discarding any with DUST score greater than 2. For k=20, we also tested the effects of sampling 1 million, 5 million, 10 million, and 20 million unique k-mers per species. We then split pairs of transcriptomic read files into 100 chunks each (using the CellBouncer program split_read_files), counted k-mers of each length using demux_species in batch mode, and joined together all resulting count files using the CellBouncer program combine_species_counts. Next, we summed all counts per cell across data from each SRA run ID corresponding to the same SRA sample ID.

Because the data came from multiple separate 10X Genomics runs, rather than a single pooled experiment, we randomly chose cells from which to create artificial inter-species doublets. For this, we downloaded the lists of cell barcodes that passed the authors’ quality control filtering (from GEO) and randomly sampled two sets of 100 cells from each species to represent doublets. Each set of 100 cells served as half of an artificial doublet. For each artificial doublet, we then scaled species-specific k-mer counts from both component cells to the same sequencing depth. To accomplish this, taking *A* to denote the first species, *B* to denote the second species, *c_A_* to denote the total species-specific k-mer counts in the cell from species *A* and *c_B_* to denote the total species-specific k-mer counts in the cell from species *B*, we computed a scaling factor for all counts from species 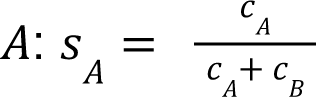 and for species *B*: 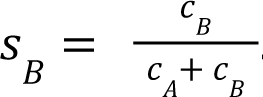. We set each species-specific k-mer count in the artificial doublet 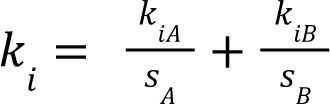 where *k_iA_* is the k-mer count indexed by *i* in the cell from species *A* and *k_iB_* is the k-mer count indexed by *i* from species *B*.

To avoid barcode collisions when merging data from each species, we randomly generated a set of 4.5 million unique 16-base barcode sequences and replaced each original barcode sequence with a random one, guaranteed to be unique across the data set. For each value of k, we then compiled species-specific k-mer counts from all species into a single file and ran demux_species without read demultiplexing (-d option enabled). We compared the output cell barcode-species assignments to the true identity of each cell, treating each original barcode + species combination as a unique key. For each value of k, we computed the sensitivity of our species assignments (the total true assignments divided by the total number of cell barcodes) as well as the specificity (the total true assignments divided by the number of cell barcodes assigned to a species). We also considered the sensitivity and specificity of assignments for cell barcodes within the filtered set that passed the authors’ QC, as well as the sensitivity and specificity of assignments for cell barcodes that demux_species identified as non-empty droplets.

##### Unlabeled data set

To benchmark computational performance and assess our method’s ability to identify true inter-species doublets, we also downloaded a public data set of a mixture of approximately 2,500 human and 2,500 mouse cells from 10X Genomics, which includes cell-species assignments made from counting reads that aligned to each species’ reference genome^39^. We obtained the reference human genome version hg38 and mouse version mm10 from UCSC^72^, and the human annotation (version 36, GTF format), and the mouse annotation (version 27, GTF format) from GENCODE^72,73^. We then extracted transcriptomes in FASTA format using gffread^74^ with arguments -g [genome FASTA] –gtf [annotation GTF] -w [output transcriptome FASTA]. For each desired k-mer length, we then counted unique k-mers in each transcriptome using FastK^70^ with option -t1 and generated unique k-mer lists using CellBouncer’s get_unique_kmers program. We made k-mer lists using the minimum accurate value of k learned in the last experiment (20), along with higher values to profile memory usage and execution time (25, 35, and 45). For each k, we sampled 20 million unique k-mers per species. We then ran demux_species with k = 20 and read demultiplexing disabled (-d option) after splitting reads into 50 files, counting k-mers using demux_species in batch mode, and joining counts using CellBouncer’s combine_species_counts program.

After inferring species identities of cells, we counted identities and compared these counts to those expected given a 1:1 human:mouse pooling ratio and 5,000 recovered cells. To calculate the observed doublet rate, we divided the inferred number of human+mouse doublets (106) by the total number of cell barcodes CellBouncer identified as cell-positive (4696) times the probability of sampling a human and mouse cell in either order (0.5*0.5*2). We also performed a chi-squared goodness-of-fit test of our observed human (2323), mouse (2267), and human+mouse doublets (106) against expected numbers, given 4688 total cells, even species proportions, and a doublet rate of 0.04, using the chisq.test() function in R. Finally, we computed the likelihood of our observed numbers of cells of each type under a multinomial distribution with parameters expected under the same assumptions (human = mouse = 0.49, human+mouse doublets = 0.02) and divided by the maximum likelihood possible given the same total counts.

##### Memory profiling

We investigated CellBouncer’s memory usage with multiple values of k, as well as STAR^40^ v2.7.10b’s memory usage when aligning to a composite reference genome, using the memory_profiler utility (https://github.com/pythonprofilers/memory_profiler). To profile memory usage and execution time while building unique k-mer lists, we wrote a script that runs FastK^70^ to count mouse transcriptomic k-mers, runs FastK again to count human transcriptomic k-mers, and then runs CellBouncer’s get_unique_kmers program to produce unique k-mer lists; we then ran mprof on this script. To profile memory usage and execution time while building a STAR reference, we ran mprof on STAR with –runmode genomeGenerate. To profile memory usage when running demux_vcf, we ran the program on one million forward/reverse read pairs, using each different value of k (20, 25, 35, and 45), and sampling 20 million k-mers per species. To profile memory usage when aligning with STAR, we ran mprof on the command STAR with –clipAdapterType CellRanger4 –outSAMType BAM SortedByCoordinate –soloType CB_UMI_Simple –soloCBstart 1 –soloCBlen 16 –soloUMIstart 17 –soloUMIlen 10 –outFilterScoreMin 30 –soloCBmatchWLtype 1MM –soloUMIfiltering MultiGeneUMI_CR –soloUMIdedup 1MM_CR –soloCellFilter EmptyDrops_CR –soloBarcodeReadLength 0 –soloFeatures GeneFull_Ex50pAS –soloMultiMappers EM. All memory profiling was done on an Ubuntu system with an AMD EPYC 7343 16-Core Processor and 500 GB total RAM. To align data from memory_profiler, we selected lines beginning with “MEM” and subtracted from all timestamps (in Unix epoch format) the minimum timestamp corresponding to that run, then converted to minutes by dividing by 60. We also converted memory values from megabytes to gigabytes by dividing by 1024.

### Demultiplexing using genotype data

#### Benchmarking

We downloaded a data set of 10X Genomics single-cell RNA-seq data from non small-cell lung cancer dissociated tumor cells from seven donors, with 40,000 targeted cells sequenced to a targeted depth of 50,000 reads per cell, and with donors labeled using 10X Genomics CellPlex labels, aligned and demultiplexed using CellRanger^41^. Because this data set did not include genotype data, we called autosomal and X-chromosome variants using FreeBayes^75^ with default parameters, using as input a copy of each donor-separated BAM file in the data set, with read groups added to mark the correct donor (via the “SM” field). We then filtered the resulting variants to biallelic SNPs minimum variant quality of 100 and fewer than 50% missing genotypes per SNP (using bcftools^76^ view -v snps -m 2 -M 2 -i “QUAL > 100 & F_MISSING < 0.5”)^62^. This left us with 511,405 variants for demultiplexing.

We created each downsampled SNP set by randomly sorting variants (Linux sort -R) and taking the number desired from the top of the list (Linux head). We downsampled reads to 10%, 25%, 50%, and 75% of the original number present using the -s option to samtools view^62^. To downsample cells, we randomly sampled barcode lists (using Linux sort -R and Linux head), converted these to CellBouncer-format cell-individual assignments files (using awk to add a placeholder sample name, a column identifying all cells as singlets, and a column indicating a Bayes factor of 100), and then used the bam_split_bcs CellBouncer utility program to create new BAM files limited to the selected barcodes.

We then ran each program on each BAM file, with default arguments. For demuxlet, we specified that genotypes should be read from the “GT” field in the VCF (--field GT). For Vireo, we specified that cell barcodes are “CB” tags in the BAM (--cellTAG CB argument to cellsnp-lite) and that genotypes should be read from the “GT” field (-t GT argument to Vireo). For demuxalot, we wrote a Python program to run the routine, as specified in the Github instructions, with default parameters, except joblib_n_jobs=1 to count_snps(), so that execution time for all programs was measured on a single core. We modified output of all programs to be in the same format as CellBouncer “.assignments” files for ease of assessing accuracy. We measured all execution times using the “real” time measured by the Linux time command. All programs were run on their own core on an Ubuntu system with an AMD EPYC 7343 16-Core Processor and 500 GB total RAM. Execution times included allele counting/single cell genotyping as well as cell-individual assignment and IO for all programs.

After running all programs, we compiled all output assignments for all cells, including the 10X Genomics CellPlex assignments (for doublets, we took the two highest-count oligos to be indicative of the doublet combination, since the CellRanger multi pipeline did not report specific doublet identities). We took the majority-vote identity for each cell to be the ground truth, and we broke all ties by choosing the 10X CellPlex assignment. Given these true identities, we computed the precision of each set of assignments as the number of assignments that matched consensus assignments divided by the number of assignments, and the recall as the number of assignments that matched consensus assignments divided by the total number of cells with consensus assignments (cells not assigned an identity by a program penalize precision but not recall). We then computed the F-score as 2*(precision *recall)/(precision + recall).

### Demultiplexing using mitochondrial haplotype clustering

#### Benchmarking

For mitochondrial demultiplexing on the full 10X Genomics NSCLC 40k data set^41^, we ran demux_mt with default parameters on a BAM file containing all data merged together. We used the same ground truth set composed of majority-vote VCF-based assignments described earlier. For all comparisons of mitochondrial to genotype-based assignments, which have different labels, we used the CellBouncer script plot/compare_assignments.R, which matches labels between sets by computing the Jaccard index between mitochondrial and genotype-based labels and choosing as a match any pair of labels for which each is the reciprocal highest Jaccard index-producing label in the other category. The script also notifies the user when the highest-Jaccard match in one set is not the reciprocal highest-Jaccard match in the other, which indicates that one or more inferred individuals likely do not correspond to true individuals. It also reports precision, recall, and the F score if either set of labels is used as ground truth. We define precision as the number of matching labels divided by the total number of test labels, and recall as the number of matching labels divided by the total number of true labels.

We downsampled the number of cells in this data set as described previously, with one exception. Since demux_mt relies on the presence of low-quality droplets containing mitochondrial reads for clustering, we wished not only to sample the desired number of high-quality cell barcodes, but also a corresponding number of low-quality droplets. On the full data set, we counted 32,669 high-quality barcodes and 903,635 barcodes with mitochondrial data that were not included in the filtered set. Assuming a linear relationship between filtered and low-quality droplets with mitochondrial reads, we decided to sample approximately 25 low-quality mitochondrial barcodes for every 1 sampled high-quality barcode in both sets. This led to our inclusion of mitochondrial data from 49,890 randomly chosen low-quality barcodes in the 2,000 cell set and 114,934 randomly chosen low-quality barcodes in the 5,000 cell set (we sampled random cells from barcode lists using Linux sort -R and Linux head, subsampled chosen cells from BAMs using CellBouncer’s utils/split_bam_bcs program, and merged BAMs with high-quality barcodes with mitochondrial-only BAMs containing low-quality barcodes using samtools^62^ merge).

We ran demux_mt on each BAM file with default parameters, then used the CellBouncer program utils/bam_indiv_rg to add read group tags using the singlet identifications in the resulting assignment file, and then ran FreeBayes^75^ with default parameters. We also ran FreeBayes on the full BAMs with no sample tags in read groups, to find variant sites for use by the other programs being tested. For all variant sets produced from the full data set, we filtered to include only biallelic variants of minimum quality 100, with fewer than 50% missing genotypes per site: bcftools^76^ view -m 2 -M 2 -v snps -I “QUAL > 100 & F_MISSING < 0.5”. For all variant sets produced from downsampled data (both the 2,000 cell and 5,000 cell set), we lowered our stringency due to data sparsity and required minimum variant quality 50 and fewer than 75% of genotypes missing per site: bcftools view -m 2 -M 2 -v snps -i “QUAL > 50 & F_MISSING < 0.75”.

For the CellBouncer MT-to-VCF pipeline, we ran demux_vcf using the filtered variant set FreeBayes created from the BAM file with mitochondrial haplotype-derived sample tags. We also ran the CellBouncer program utils/refine_vcf (described in the next section) on this variant set, with the cell-individual assignments just created and with variant site filtering disabled (--p_thresh 0), to produce a refined set of genotypes that maximize the likelihood of the reads together with the cell-individual assignments. We then ran demux_vcf again with this new genotype set to produce “CellBouncer_refined” cell-individual assignments.

We then ran freemuxlet^19^, scSplit^18^, souporcell^17^, and Vireo^15^ on each data set, with default parameters and 7 expected individuals for each program, using the filtered variant sets from FreeBayes (with no sample tags) as loci to genotype and cluster for all programs except souporcell, which selects variant sites as part of its pipeline. Two programs we tested, freemuxlet and scSplit, did not successfully run on the full data set. This was due to excess memory usage (freemuxlet killed in the dsc-pileup stage, running alone on a system with 500 GB of RAM), and execution time (scSplit ran for over 71 hours without completion).

We converted each program’s output to a CellBouncer-format assignments file, to evaluate results against consensus truth assignments using plot/compare_assignments.R. We used the F score computed by this script and took note of when the highest-Jaccard match to an identity was not reciprocal.

Because variants where genotypes are identical across all sites are useless for distinguishing between individuals, counts of output variants from each program (or those that remained after filtering FreeBayes output, in the case of CellBouncer’s MT-to-VCF pipeline) only included sites where more than one genotype exists among all inferred individuals.

To test whether demux_mt could recover mitochondrial haplotypes from scATAC-seq data, we downloaded ATAC-seq reads from a study of hematopoiesis that sorted and sequenced cells from three donors, processed separately^42^ (ArrayExpress ID E-MTAB-9068). Because these data came from a plate-based method, we generated a random 16 bp cell barcode sequence to correspond to each cell we downloaded, ensuring that each barcode was unique across the data set. We then inserted these barcodes in the format of SAM tags into FASTQ sequence comments (in the format CB:Z:[barcode sequence]). We aligned reads to the hg38 reference genome with minimap2^77^ in short read mode, outputting SAM and inserting sequence comments as SAM tags (minimap2 -y -a -x sr), and then compressed to BAM, sorted, and indexed using samtools^76^. We merged BAMs for all cells from each donor (using samtools merge), extracted mitochondrial sequences from each donor (using samtools view -bh chrM), and merged and indexed mitochondrial mappings from all three donors. We then ran demux_mt on this merged BAM file with the doublet rate set to zero (-D 0), recovering three haplotypes. Finally, we created a CellBouncer-format assignments file to denote the known true identity of each cell (using the downloaded metadata) and compared the true assignments to mitochondrial clusters using the CellBouncer script plot/compare_assignments.R.

### Ambient RNA quantification, profiling, and removal

#### Synthetic ambient RNA data set

For our main test of our ambient RNA inference and bulk proportion profiling tools, we created a data set contaminated with synthetic ambient RNA from a known source and in a known amount. For this, we first randomly sampled 1000 cells from the 10X Genomics NSCLC data set^41^ to use as a baseline data set. We then counted the number of reads per cell in this data set and created a data set of candidate contaminant reads by eliminating the 1000 sampled cells from the larger data set. We then calculated the number of contaminant reads required to replace the desired amount of each cell’s RNA: if *n* is the total read count in a cell and *p* is the desired fraction of ambient RNA contamination, we sampled 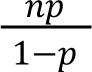 contaminant reads, skipping those missing cell barcodes, duplicate reads, unmapped reads, and secondary alignments. Each contaminant read’s cell barcode was adjusted to match the desired target for contamination.

We repeated this process 35 times, introducing 5%, 10%, 15%, 20%, or 25% contamination into each cell, originating from one of the seven donors per file, to create 35 new datasets. On each data set, we ran CellBouncer’s demux_vcf, along with demuxlet, demuxalot, and vireo with default parameters. We then ran CellBouncer’s quant_contam on each with -D 0.1, to allow it to make doublet identities less likely when they make up too much of the data set. F-scores of all resulting cell-individual assignments, including refined assignments from quant_contam, were computed by comparing the results to the consensus true identities used earlier (the majority vote ID of each cell from running demux_vcf, demuxalot, demuxlet, and vireo on the full data set and including the 10X CellPlex-derived labels, using the CellPlex IDs to break ties).

To assess how negatively affected the inferred ambient RNA profile was by cell mis-identifications in the high-ambient RNA datasets, we also ran quant_contam again on each data set, using the cell-individual assignments from demux_vcf run on the baseline data set in place of the individual assignments from demux_vcf run on the contaminated data set. Since these cell-individual assignments were more likely to be correct, we expected the inferred ambient RNA profile to be more accurate, and the degree of difference to help illuminate the extent to which quant_contam’s accuracy suffers in the face of high ambient RNA data.

We also ran bulkprops on each of these datasets, using the same SNP set used to run the SNP-based demultiplexing tools. Each bulkprops run used -e 0.001 to ensure a consistent error rate in the model across runs.

#### SMART-seq2 data set

We downloaded gene expression data from a published study that performed SMART-seq2 RNA-seq on human fetal liver and bone marrow blood cells^42^. To make the data amenable to processing using 10X Genomics’ CellRanger pipeline, we selected a random 16-base barcode from the CellRanger barcode whitelist for every cell in the data set. We then split apart read pairs, using every real read as the reverse read in a pair, and generating for each a synthetic forward read consisting of the selected cell barcode sequence followed by 34 random bases (with “?” quality characters for each) to achieve 50bp forward reads containing completely random UMIs. We aligned these data to the CellRanger-provided GRCh38 reference version 2024-A using CellRanger version 8.0.1.

To obtain a high-quality set of variants, we used STAR^40^ v2.7.2a (with –outSamType BAM sortedByCoordinate) to align all downloaded read pairs to the CellRanger-provided GRCh38 reference version 2020-A, added read groups to tag individuals of origin using the bam_dummy_rg program from https://github.com/nkschaefer/bio_misc, and called variants on the autosomes and X chromosome using FreeBayes^40,75^ with default parameters. We then used bcftools^76^ to filter the resulting variant file to biallelic SNPs with minimum total read depth of 100 using the command bcftools view -m 2 -M 2 -v snps -I ‘INFO/DP >= 100’ and, to further improve variant accuracy, we ran CellBouncer’s refine_vcf tool on the resulting file, with –p_thresh 0.01 and using the true identity of each cell as the cell-individual assignments.

Using this variant call set, we ran demux_vcf and quant_contam twice, once with default parameters and once with -D 0 for both programs, to reflect that doublets are not possible with this technology. We then ran DecontX^28^ on the raw_feature_bc_matrix with default parameters.

#### Full NSCLC data set

We ran each contamination inference tool on the full NSCLC data set. To run cellbender^29^, we first subset the raw gene expression matrix to include only gene expression data, using the CellBouncer tool subs_mex_featuretype.py. We then ran cellbender remove-background with the arguments –expected-cells 3300 –model full –epochs 150. We extracted per-cell contamination rates from the /droplet_latents/background_fraction field in the H5 output, and the ambient gene expression profile from the /global_latents/ambient_expression field. To run DecontX^28^, we used the filtered_feature_bc_matrix data, extracted per-cell contamination estimates from the contamination field in the result object, and extracted the ambient RNA expression profile with the R command rowSums(metadata(sce)$estimates$all_cells$eta), where sce is the result object, and then normalized all elements of this vector by dividing them by the vector’s sum. For SoupX^27^, we used the clusters inferred by DecontX for the sake of consistency, extracted the ambient expression profile from the soupProfile field in the result object, and measured each cell’s contamination rate as 1 – colSums(afterMatrix)/colSums(beforeMatrix), where beforeMatrix was the filtered_feature_bc_matrix data and afterMatrix was the output of adjustCounts. To run CellBouncer, we used default parameters for demux_vcf and quant_contam, but provided DecontX-inferred clusters to quant_contam for the sake of consistency. We then compared each pair of ambient gene expression profiles, and every pair of per-cell ambient RNA fractions, using the cor function in R.

#### Assessing subcellular localization of genes expressed in ambient RNA

We downloaded data from lncAtlas^47^, which is a database of results from bulk RNA-seq experiments in which the relative concentrations of different genes in different subcellular locations were measured. lncAtlas reports results as relative concentration indices (RCIs), which are defined as the log_2_ RPKM of a given gene in one cellular fraction minus the log_2_ RPKM in another. The database includes measurements of relative concentrations of transcripts in the cytoplasm versus nucleus in multiple cell lines, as well as additional subcellular localization data in K562 cells. After downloading, we calculated the mean RCI per gene and intersected all RCI data with ambient gene expression vectors from all programs by excluding any gene with a HGNC symbol appearing more than once in our data set (the 10X Genomics NSCLC data) and matching lncAtlas with ambient expression vectors using Ensembl gene IDs. We then computed the Spearman correlation between each ambient expression vector and lncAtlas RCIs using the R function cor with method=’spearman’ and use=’pairwise.complete.obs’.

#### Comparing with previously published ambient RNA gene list

We downloaded a list of genes found by a prior study^33^ to be highly expressed in ambient RNA and intersected this list with each of our inferred ambient expression vectors. We performed a series of Mann-Whitney U tests to compare the ambient expression vector loading from each program for genes not in the list against genes in the list, using the wilcox.test function in R.

#### Comparing with numbers of spliced reads and gene log likelihoods

We counted spliced and unspliced transcripts in the 10X NSCLC data set using velocyto^78^ and extracted the spliced counts from the output .loom file, intersecting these counts with all inferred ambient expression profiles for all genes with HGNC names appearing only once in the data set.

We obtained mean log likelihood values per gene under the consensus, data set-wide donor mixture proportions, by running bulkprops “in reverse” as described in Supplementary Methods. To make these values easier to visualize, we rank-normalized them using the ecdf function in R.

### Processing custom tag and CRISPR guide capture sequencing data

#### Downsampling sgRNA counts from Perturb-Seq data

We downloaded sgRNA capture data from Replogle *et al* (2020)^32^ and extracted counts for the iPSC experiment, in which the authors targeted two guides per cell, in MEX format. We then split the MEX data into a separate matrix per library using the CellBouncer program utils/split_mex_libs.py. For the full-coverage data set (average 999 guides per cell), we ran CellBouncer demux_tags on each library, with the –sgRNA option enabled. We also assigned guides for each library using the mixture model-based strategy proposed by the study authors, via code adapted from https://github.com/josephreplogle/guide_calling. After assigning guides using each method to each library separately, we combined the output .table files using the CellBouncer program utils/combine_sgrna_tables.py after appending unique library identifiers to cell barcodes. We then merged together the two tables in R and computed the number of cell barcodes that received each combination of guides assigned by the two methods, using the table command in R. We then repeated this process after downsampling counts to each desired level. To achieve this, we chose a sampling probability (1%, 5%, 10%, 25%, or 50%, yielding an expected 10, 50, 100, 250, or 500 counts per cell respectively) and adjusted each count in the MEX file by choosing whether or not to keep each read given the sampling probability. For example, with a sampling probability of 50% and a count of 10, we would draw 10 random uniformly distributed numbers between 0 and 1 using python’s random.random() function, and set the downsampled count to the number of these samples for which the random number was below the sampling probability (0.5 in this case).

#### Benchmarking using 10X Genomics CellPlex data

We downloaded CellPlex reads from the 10X Genomics NSCLC data set^41^ and counted tags in the reads using demux_tags. We then randomly downsampled reads three times per desired sequencing depth using seqkit sample^79^, with a seed of 100, 200, and 300 for the three samples and sampling probability (-p argument) set to 0.001, 0.0025, 0.005, 0.01, 0.05, 0.1, and 0.5 to produce all required samples. We ran demux_tags to count tags in each downsampled data set as well, and we ran each program being tested (demux_tags, BFF^21^, deMULTIplex2^22^, demuxmix^23^, GMMdemux^24^, hashedDrops^26^, hashSolo^25^, and HTOdemux^20^) on these tag count tables produced by demux_tags. Because not all of these programs provide information about specific doublet types when cells are assigned doublet identities, we treated all doublet types assigned by all programs as a single “multiplet” identity, and we removed all cells assigned a “negative” or undetermined identity. We then compiled a ground truth for each cell by taking the majority vote identity assigned to each cell by demux_vcf, demuxlet^14^, Vireo^15^, and demuxalot^16^ using genotype data, again converting specific doublet identities to a single “multiplet” identity and discarding cells for which the majority-vote identity was a tie. For each program’s identifications on each data set, we computed precision as the number of matching identifications divided by the number of inferred identities, recall as the number of matching identifications divided by the number of true identities (excluding cells whose identities were not inferred), and the F score as 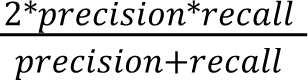.

To simulate datasets with decreased signal-to-noise ratio, we loaded the full-count data set and, using the consensus ground truth identities, transformed each foreground/true tag count to remove 10%, 25%, 50%, 75%, or 90% of the desired fraction of counts, leaving all background counts unaltered. We then repeated the same process as above, running each program on the transformed counts and assessing accuracy using the ground truth labels for each cell. We note that, while the three replicates of each downsampled data set allowed stochasticity in each program’s method as well as in the downsampling process to alter results, the three replicates of each low signal-to-noise data set used identical datasets (there was no randomness used in producing these datasets), meaning that the differences in program output here were entirely due to stochasticity in non-determinism in the programs.

### Inferring consensus pool proportions and global doublet rate

#### Assessing NSCLC data set labels using doublet_dragon

To obtain an estimate of the true doublet rate in the NSCLC data set^41^, we ran doublet_dragon on the genotype-based identities assigned by demux_vcf, demuxlet^14^, demuxalot^16^, and Vireo^15^ together, inferring a consensus global doublet rate of 0.187848. We then computed an expected heterotypic doublet rate by subtracting the expected fraction of homotypic doublets, 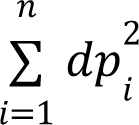 using the same variable names as in the last section. This gave us an expected heterotypic doublet rate of 0.1572569.

To compare the output of demux_tags, demux_vcf, and demux_mt run on this data set, we ran doublet_dragon on the output of each program separately, comparing the pool proportions to those from running bulkprops on the full data set with default parameters, and to the consensus global doublet rate inferred above.

We also computed the heterotypic doublet rate from running each HTO assignment program on each downsampled and low signal-to-noise ratio data set we created for benchmarking demux_tags. To this end, we divided all cells receiving doublet or multiplet identities by the total number of cells assigned any identity. We then compared these numbers to the consensus heterotypic doublet rate inferred using genotype data above.

#### Generation and growth of episomally-reprogrammed iPSC lines

Human and Chimp iPSC lines previously described^80,81^ were used for all cell fusion experiments. We also generated an additional Bonobo (Congo) and chimpanzee (40670) iPSC line as described previously^80^. Briefly, fibroblasts were grown at 5% CO_2_/atmospheric O_2_ in α-MEM supplemented with 10% FBS, NEAA, GlutaMAX, 64 mg/L L-ascorbic acid 2-phosphate trisodium salt and Primocin until 70-80% confluence and released by trypsinization for transfection. 1.5×10^6^ cells were transfected with 1.25 μg per episomal vector containing the following genes: OCT3/4, SHp53, SOX2, KLF4, LIN28, and L-MYC (Addgene plasmids 74947, 27078, and 74944)^82,83^. Transfected cells were seeded at 15,000/cm^2^ on tissue culture plates precoated with 1 μg/cm^2^ vitronectin. Cells were grown in Essential 6 media (made in-house as previously described^84^ without TGFβ1, supplemented with 0.5 mM sodium butyrate and 100 nM hydrocortisone (Sigma Aldrich). Hydrocortisone was used between days 1-12, or until cell density exceeded >70% confluence. Colonies began to form at days 18-22 and were picked between days 24-30 onto dishes coated with γ-irradiated CF-1 derived mouse embryonic fibroblasts (MEF) and subsequently grown in 50/50% ratio of iDEAL/E8 feeder-free medium^85^, iPSC lines were expanded 4-8 passages on feeders before being transitioned to feeder-free conditions. Higher rates of spontaneous differentiation were routinely observed when growing Bonobo iPSC clones on MEF, to reduce rates of spontaneous differentiation, 50 nM Dorsomorphin was added to the medium while expanding on MEF as recommended previously^86^, Feeder-free iPSC lines were expanded on Growth Factor Reduced Matrigel using EDTA-based cell release solution^84^ and grown in a custom iPSC media made in-house (v3.1 medium): 65 ng/ml FGF2-G3, 2 ng/ml TGFβ1, 0.5 ng/ml NRG1, 20 μg/ml insulin, 20 μg/ml holo-transferrin, 20 ng/ml sodium selenite, 200 μg/ml ascorbic acid 2-phosphate, 2.5 mg/ml bovine serum albumin (BSA), 30 ng/ml heparin, nucleoside supplement (15 μM adenosine, 15 μM guanosine, 15 μM cytidine, 15 μM uridine, and 6 μM thymidine) in Dulbecco’s Modified Eagle’s Medium (DMEM)/Ham’s F-12. During routine passaging on both feeders and feeder-free conditions, media was supplemented with 2 μM thiazovivin (MedChemExpress, HY-13257) to boost survival.

#### Generation of Fusion Reporter Lines

To facilitate rapid selection and visualization of tetraploid cell fusions, we generated reporter lines coexpressing a fluorescent protein and drug selectable marker. Piggybac transposon constructs expressing a whole cell localized mNeonGreen and puromycin resistance cassette (mNEON-Puro), or a nuclear localized mCherry2 and Zeocin resistance cassette (mCherry-Zeo) were generated using Gibson mediated cloning of gBlock fragments (IDT) containing the optimized cDNA sequences into a piggybac expression vector derived from addgene plasmid# 133379^87^. For each reporter line, 1 million cells were electroporated with 5 μg of the piggyback expression plasmid using a Lonza 4D nucleofector device, and program DN100. To facilitate insertion of the piggybac cassettes, an expression vector for the piggybac transposase was coelectroporated, at a ratio of insert: transpose of 5:1. Following electroporation, cells were recovered in V3.1 media containing CEPT^88^ for 24 hours. Electroporated cell lines were selected using 0.25-2 μg/mL puromycin or 1-5 μg/mL Zeocin, the concentration of each drug used for selection was started at the lowest amount specified and gradually increased (by 0.25-0.5 μg/mL increments for puromycin and 1-2 μg/mL increments for Zeocin) until reaching the upper limit of the drug used.

#### Generation of Tetraploid Cell Fusion Lines

Tetraploid cell fusions were generated using an on plate DMSO-PEG1500-mediated cell fusion approach^89^. Briefly, 100,000 cells from two different reporter lines (mNEON-Puro or mCherry-Zeo) were mixed and plated into one well of a 24-well glass bottom plate (Cellvis) and allowed to recover overnight in V3.1 iPSC media supplemented with CEPT^88^. Cells were washed once with DPBS followed by addition of 300 μL 45% PEG 1500 (Roche#10783641001) / 10% DMSO (25-950-CQC), After a 2 minute incubation, cells were washed twice with DPBS (500 μL per wash) and once with iPSC media before final media replacement. Selection of tetraploid cell fusions was performed starting 48 hours post-fusion by using a drug step up pulsing approach starting with 0.25 μg/mL puromycin (1 μg/mL) and 0.75 μg/mL Zeocin, selection pressure was increased as the cells grew by doubling the concentrations of the respective drugs until the maximum concentration of 1 μg /mL Puromycin/ 3 μg /mL Zeocin was reached. This approach resulted in well-formed double positive colonies with close to 100% success across a broad range of fusions. Selection pressure was maintained for 2-3 passages and removed on the final passage for cell banking.

#### 10X#single cell RNA-seq library construction

Tetraploid iPSC lines were grown in V3.1 media until 80-100% confluent and subsequently dissociated with Accutase. To provide orthogonal labeling for cell line of origin, samples were labeled using MULTI-seq barcodes as described previously^50^. 500,000 cells were resuspended in PBS and transferred to 96 well plates containing DNA barcodes (200 nM) and lipid modified oligo (LMO) anchors (200 nM) for 5 minutes on ice, then transferred to plates containing the LMO co-anchor (200 nM) for an additional 5 minutes on ice. Excess LMO/barcodes was removed by two washes using PBS + 50 mg/mL, followed by a wash using PBS + 25 mg/mL BSA. Labeled samples were resuspended in PBS + 10 mg/mL BSA, pooled together in sets of 8-12 samples, and adjusted to 4 million cells/mL before sample loading. Single-cell RNA sequencing was collected by loading approximately 20,000 cells per lane and captured using the 10x Genomics Chromium X controller with version 3.1 RNA capture kits. Library preparation was performed according to manufacturer’s instructions and sequenced on Illumina HiSeq and NovaSeq X platforms.

#### Read alignment and quality control

To avoid bias that would be introduced by aligning data to any one species’ reference genome, we obtained a genome representative of the most recent common ancestor of humans, chimpanzees, and bonobos, imputed as a byproduct of a whole-genome progressive Cactus alignment^90^ of multiple great ape genomes^91,92^ and annotated using the Comparative Annotation Toolkit^93^, produced as part of a recent study^91^. We also produced an ancestral mitochondrial genome by aligning the human (hg38), chimpanzee (panTro6^92^), bonobo (panPan3^91^), and gorilla (gorGor6^92^) mitochondrial sequences using MUSCLE^94^ after shifting the circular mitochondrial genome breakpoints to align with the human reference’s linear coordinates. At each position, we chose the human base if it matched either bonobo or chimpanzee, or the gorilla base otherwise, ignoring positions where the chosen base was a gap. We then annotated this genome using liftOff^95^, with the GENCODE^73^ v36 human mitochondrial annotation as source. Finally, we used a custom pipeline (https://github.com/nkschaefer/litterbox) to remove types of transcripts known to be problematic for comparative analysis, such as readthrough transcripts, and to rename genes to the current HGNC recommendations.

Additionally, to avoid losing mitochondrial reads to alignments with nuclear mitochondrial insertion (NUMT) sequences, we masked NUMTs according to a procedure outlined previously^96^ and incorporated into a pipeline: https://github.com/nkschaefer/numty-dumpty. The procedure entails simulating mitochondrial reads, aligning to a version of the nuclear genome lacking the mitochondrial sequence, calling peaks with MACS2^97^, and converting sequence within the peaks to “N” using bedtools maskfasta^98^.

We aligned data using 10X Genomics CellRanger v8.0.0 with default parameters. After alignment, we used Scanpy^99^ for visualization and quality control before pseudobulk and differential expression analyses. From visual inspection, we chose to discard cells with more than 20% mitochondrial reads from pseudobulk for differential gene expression analysis.

#### Sample demultiplexing and QC

Using variants for all contributor cell lines produced from deep whole genome sequencing data as part of a prior study^34^, we ran demux_vcf on each library, along with demuxalot^16^, demuxlet^14^, and Vireo^15^ with default parameters. Although the other programs ran quickly, the large number (∼12M) of SNPs, owing to inclusion of many sites fixed between species, prevented demuxlet from running to completion within the allotted time of 12 days. In this data set, each program reported allotetraploid composite cells as doublets; we were therefore unable to detect true doublets of composite cells using genotype-based demultiplexing. We provided an allowed list of “doublet” identities to demux_vcf using the -I argument, which resulted in higher accuracy by preventing the program from choosing identities that were not included in the experiment.

We then used demux_mt to count MULTIseq tags in reads and assign identities (composite cell lines of origin) to each cell. We note that relatively shallow sequencing of these tags, as well as at least some evidence for MULTIseq tag bleeding into neighboring wells, likely lowered the accuracy of MULTIseq-inferred identities, but the MULTIseq data still provided a helpful way to identify and discard doublets that would otherwise be invisible to genotype-based demultiplexing.

We noticed an issue with our data for library B that resulted in low agreement between the MULTIseq label for fusion line 27 and its expected genotype, resulting in relatively poor genotype-MULTIseq agreement on this library (Table S7). Because of this, we discarded any cells for which the genotype-based assignment (from demux_vcf) disagreed with the MULTIseq-based assignment (from demux_mt) from all downstream analyses.

We assessed the accuracy of genotype-based identities using MULTIseq tags as ground truth after discarding all cells identified as doublets from the MULTIseq data.

Next, we inferred ambient RNA contamination using quant_contam on the demux_vcf output on each library, after discarding cells from the “.assignments” file that were inferred to be doublets from MULTIseq data. We also ran SoupX^27^, DecontX^28^, and cellbender^29^ on each library, and extracted cell-specific ambient RNA fractions from the output of each as described earlier. SoupX failed to run successfully on one of our libraries (A1), and so this library was excluded from results.

To obtain an independent estimate of ambient RNA contamination per library, we ran bulkprops on each using the same variant data. We then computed bulk proportions from our inferred cell-individual assignments. Because our libraries consisted of a single cell type likely to be uniform in RNA content, we reasoned that the discrepancy between cell-individual assignments and inferred bulk proportions could be explained as arising from ambient RNA. To get a rough estimate of ambient RNA contamination, we made the simplifying assumption that ambient RNA originated only from cell lines overrepresented in the bulkprops output compared to the cell assignments. Then, if *b_i_* is the fraction of bulk RNA originating from individual *i* according to bulkprops, *_i_* is the fraction of bulk RNA originating from individual *i* according to cell-individual assignments from demux_vcf, *c* is the global contamination rate, and *f_i_* is the fraction of ambient RNA composed of individual *i*, then for each *i* where *b* > *a_i_*, we assume the excess RNA must originate from cells not assigned to individual *i*:

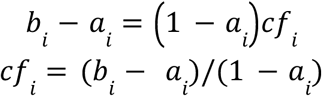

Since by definition, 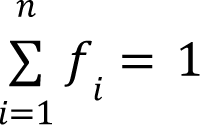, and 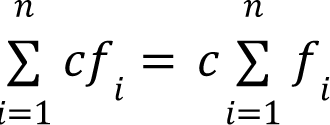, then

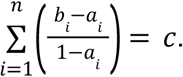

We compared the per-cell ambient RNA estimates from each program in each library to estimates produced this way.

#### Assigning mitochondrial haplotypes to cells

In datasets such as ours, in which very divergent mitochondrial haplotypes can exist, one run of demux_mt risks recovering one consensus haplotype per group (in our case, species), rather than finding all haplotypes in the data set. For this reason, we created a CellBouncer program called mt_subcluster.py, which takes as input the results from one run of demux_mt, as well as the IDs of clusters to sub-cluster. It then re-runs demux_mt on each chosen cluster, limiting analysis to cell barcodes assigned to that cluster in the previous run. After all runs are complete, it then merges all resulting variants together, creates a consensus haplotype for each cell, and identifies each cell as belonging to one of the final haplotypes. One run of mt_subcluster.py was enough to recover most mitochondrial haplotypes in our technical replicates of the same library (A1 and A2), but recovering all human haplotypes took two additional rounds of subclustering in the other library (B). This is likely due to uneven representation of the haplotypes; demux_mt assumes counts of cells belonging to all haplotypes follow a single Poisson distribution, and this assumption can be violated when haplotypes exist at very uneven proportions.

To obtain a more complete set of mitochondrial haplotypes, we called variants on our WGS data using FreeBayes^75^ on the mitochondrial genome with ploidy set to 1. We then converted these data from VCF format into the format used by demux_mt.

We then added an option to demux_mt to allow counting of allele-specific reads in tetraploids (and cells identified as doublets of two identities). This option takes a set of mitochondrial haplotypes and cell-individual assignments (*e.g.* from demux_vcf) and outputs, at each site segregating between the two mitochondrial haploypes for each doublet/tetraploid cell, the number of reads carrying the allele from each individual. For all of our human/human and chimpanzee/chimpanzee composite cells, we used this feature to calculate allele specific reads at SNPs, then intersected these counts with mitochondrial gene coordinates (in BED format) using BEDTools^98^ intersect, and summed allele-specific counts across genes in R.

To avoid noise arising from low-frequency sites, we chose to use only fixed species-specific mitochondrial variants for our inter-species comparisons. To this end, we selected (using bcftools^76^ view) only variants fixed between human and chimpanzee cell lines and produced demux_mt-format mitochondrial haplotype files at these sites, with one haplotype for humans and one for chimpanzees, with the “0” allele at each site representing human and the “1” allele at each site representing chimpanzee. We repeated this to create another haplotype file for sites fixed between chimpanzees and bonobos. We then counted allele-specific mitochondrial gene expression using the same procedure as above, but to help us calibrate our methods, we computed human/chimpanzee mitochondrial allele specific counts in human/human and chimpanzee/chimpanzee composite cells, as well as in human/chimpanzee composite cells.

To assign a mitochondrial haplotype to every cell using these allele-specific counts, we used version 0.15.0 of the Pomegranate mixture model fitting library^100^ with custom code to allow fitting beta-binomial distributions. For each set of allele-specific counts, we first assigned every gene in every cell to a distribution of origin using a mixture model. For inter-species allele-specific counts, we fit a three-way beta-binomial mixture model, with parameters initialized to (0.1, 9.9) (species 1 mitochondria), (5, 5) (both species mitochondria), and (9.9, 0.1) (species 2 mitochondria). We then summed the log probabilities of each gene belonging to each distribution, across cells, and assigned cells to identities by choosing the highest-probability distribution.

Intra-species allele counts were much lower than inter-species allele counts, owing to there being far fewer segregating SNPs between each pair of haplotypes. For this reason, we first summed allele counts across all genes for each cell before fitting distributions. We then split cells into groups by species, ensured each pair of allele counts had the higher count first, and fit a two-way mixture model representing cells with two mitochondrial haplotypes (initial parameters (10*m, 10*(1-m))) and those with one mitochondrial haplotype (initial parameters (9.9, 0.1)), where m was the mean fraction of (individual 1 count)/(individual 1 count + individual 2 count) across all cells. We did this after realizing that many intra-species cells showed evidence for retaining two mitochondrial haplotypes, and that the allelic ratios of haplotypes often differed significantly from 0.5. After fitting the mixture model, we assigned each intra-species cell to both mitochondrial haplotypes, or to the mitochondrial haplotype with the higher read count, according to the probability obtained through fitting the mixture model.

Once all cells were assigned to one or more mitochondrial haplotype, we sought to assess our method’s reliability by also assigning cells known to be of human/human origin to human and/or chimpanzee mitochondrial haplotypes. Given prior literature, and our observation that human mitochondria are more common than chimpanzee mitochondria in human/chimpanzee composite cells, we sought to calculate the rate at which our method would need to falsely identify cells with only human mitochondria as having either only chimpanzee mitochondria, or human and chimpanzee mitochondria, if in fact all human/chimpanzee cells contain only human mitochondria.

Denoting cells that truly contain only human mitochondria as *T_H_*, cells that truly contain only chimpanzee mitochondria as *T_C_*, cells that truly contain both mitochondria as *T_HC_*, cells labeled as having only human mitochondria as *L_H_*, cells labeled as containing only chimpanzee mitochondria as *L_C_*, and cells labeled as having both types of mitochondria as *L_HC_*, we defined our false discovery rate of interest as *FDR* = *P*(*L_HC_*) + (*T_H_*|*L_C_*). Using Bayes’s theorem:

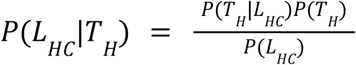

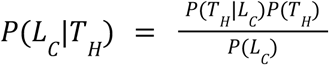

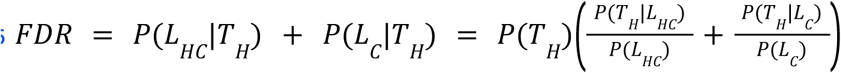

We calculated *P*(*L_HC_* |*T_H_*) and *P*(*L_C_* |*T_H_*) by dividing the number of human/human cells assigned both human and chimpanzee mitochondria, or only chimpanzee mitochondria, by the total number of human/human cells, and we computed *P*(*L_HC_*) and *P*(*L_C_*) by dividing the total occurrences of those labels across all cells by the total number of cells. To discover the false discovery rate of interest, we set *P*(*T_H_*) = 1 to represent the possibility that our observation was wrong and all human/chimpanzee cells contain only human mitochondria.

According to our data, we misidentified 2.4% of human/human cells as having human and chimpanzee mitochondria and none as having chimpanzee mitochondria. Using the method above, we calculated that a false identification rate of 26.18% would be necessary to explain all of our results, if no human/chimpanzee cells truly inherited chimpanzee mitochondria

#### Comparing mitochondrial allelic ratios to divergence between haplotypes

We measured the divergence between each pair of within-species mitochondrial haplotypes by loading the aforementioned mitochondrial haplotypes we constructed, which consist of biallelic SNPs, with the two alleles at each arbitrarily encoded as “0” and “1”, and computing Euclidean distances using the dist function in R.

#### Differential expression analyses

After assigning each cell to one or more mitochondrial haplotype, we assigned every cell to a pseudo-bulk bin based on its library of origin, the species of origin of both of its contributor cell lines, and the species of origin of its mitochondria. After discarding all cells with more than 20% mitochondrial reads, as described earlier, we summed expression of all genes within each pseudobulk bin and performed differential expression analysis with DESeq2^101^ twice: once including all genes in the annotation and a second time including only mitochondrial genes. Because the size factors computed in the first analysis normalized expression genome-wide, while those in the second analysis normalized expression to mitochondrial totals, the second analysis should reflect only differences in mitochondrial post-processing, while the first reflects differences in mitochondrial copy number, expression, and post-processing. We then assumed these different drivers of expression differences to be multiplicative and approximated differences due only to copy number and expression level by subtracting log_2_ fold change values from the second analysis from the first. When reporting significant DE values, we used p = 0.01 as a cutoff.

#### Correlation of neighboring genes

Considering only each pair of adjacent protein-coding genes on the mitochondrial heavy strand, we loaded single-cell gene expression matrices in R (using the readMM command in the Matrix package) and grouped cells based on the species of origin of their nuclear genomes as well as their mitochondria. In each group, we then computed the Pearson correlation coefficient between the expression of genes in each pair using the corr function in R.

#### Gene Ontology enrichment

We performed Gene Ontology enrichment analyses using the R package GOfuncR version 1.22.0, built on the FUNC^102^ package, with the included human gene ontology data. Enrichments were calculated using the go_enrich function with test=’hyper,’ and all genes with adjusted p < 0.01 included in gene sets.

#### Separating mitochondrial from nuclear regulatory changes

Both for comparing DE of genes on the mitochondrial light vs heavy strand, and for comparing the correlation of expression of neighboring gene pairs in human versus chimpanzee, we sought to use all data we collected to disentangle regulatory changes stemming from differences in mitochondria from regulatory changes stemming from differences in nuclear genomes. To this end, we reasoned that measurements from human/human cells come from interactions between human nuclear genes and human mitochondria, while those from chimpanzee/chimpanzee cells reflect interactions between chimpanzee nuclear genes and chimpanzee mitochondria; comparisons between human/human and chimpanzee/chimpanzee cells therefore reflect both mitochondrial and nuclear differences.

Since human/chimpanzee cells contain both species’ nuclear genomes, measurements from these cells with a single species’ mitochondria reflect the influence of both nuclear genomes on a single species’ mitochondria. Therefore, comparing human/chimpanzee cells with human mitochondria to human/chimpanzee cells with chimpanzee mitochondria is a way to isolate mitochondrial differences: differences reflect the influence of a common nuclear background on the two different mitochondria.

To isolate nuclear differences, we could compare human/chimpanzee cells with a single species’ mitochondria to same-species composite cells with matching mitochondria. For example, human/chimpanzee cells with human mitochondria reflect human and chimpanzee nuclear factors with human mitochondria, so comparing these to human/human cells should isolate the influence of chimpanzee nuclear factors. If we abbreviate human as H and chimpanzee as C with normal script used to denote mitochondrial species and subscript used to denote nuclear genome species, and working in log space, then:

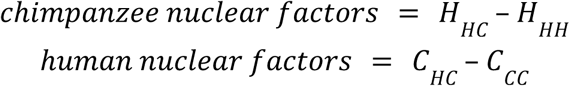

We can then compare human to chimpanzee nuclear factors as

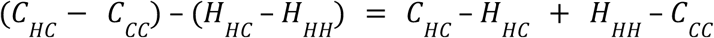

Since we already defined nuclear + mitochondrial differences as *H_HH_* − *C_CC_* and mitochondrial differences as *H_HC_* − *C_HC_*, this is consistent with the expectation that nuclear + mitochondrial = *C_HC_* − *H_HC_* + *H_HH_* − *C_CC_* + *H_HC_* − *C_HC_* = *H_HH_* − *C_CC_*.

We chose to omit data collected from human/chimpanzee cells with both species’ mitochondria from this analysis, due to our observation of cross-species mitochondrial interactions inhibiting normal gene expression.

#### Assessing allele frequency of human-specific mutation

We identified a fixed, human-specific derived allele on the mitochondrial genome at the position adjacent to a proposed MT-ND3/MT-ND4L cleavage site, using mitochondrial genomes aligned as described earlier when building the reference genome. In this case, however, we also included the Neanderthal reference mitochondrion^103^. We used the gorilla allele to infer the ancestral state.

Using the GnomAD population sequencing database^52^, we found a back-mutation at the site from the derived T allele to the ancestral C allele (rs1603222851). To compare the frequency of this mutation to the frequencies of other types of mutations, we downloaded all mitochondrial variants from GnomAD and removed all with homoplasmic allele frequency == 0. We then converted all Variant Effect Predictor (VEP) ^104^ annotations into the broad categories used by VEP: high (start lost, stop lost, frameshift), moderate (inframe deletion, missense), low (incomplete terminal codon, start retained, stop retained, synonymous, coding sequence), and modifier (non coding transcript exon). Because of the small size of the mitochondrion resulting in relatively few high-impact mutations, we chose to compare mutations with either moderate or high predicted impact to those with low predicted impact.

We then similarly grouped annotations from ClinVar^105^ into two broad categories: pathogenic (encompassing affects, likely pathogenic, pathogenic/likely pathogenic, pathogenic, likely pathogenic; drug response, and pathogenic; drug response) and benign (association not found, benign/likely benign, benign, likely benign). We then compared the homoplasmic allele frequencies of mutations across both sets of categories.

## Supplementary Text

### Pitfalls of mitochondrial haplotype clustering

Although demux_mt works in many cases, we note that factors such as cellular stress and tissue degradation can greatly influence the number of mitochondrial reads, which will affect the ability of demux_mt to recover mitochondrial haplotypes. Further, some genomes, such as those of nonhuman primates, contain many nuclear mitochondrial insertions (NUMTs)^106^, which are likely to reduce the number of true mitochondrial reads aligned to the mitochondrial genome; finding and masking these sequences might be necessary, as has been done previously for other studies^96^. Additional pitfalls include using tissue or cell lines derived from closely related laboratory animals that might share a recent mitochondrial ancestor, or cell populations that exist at very different proportions in the pool (demux_mt assumes counts of all haplotypes follow a single Poisson distribution; see Online Methods).

We attempted to cluster a 15-donor RNA-seq data set^42^ produced using the plate-based method SMARTseq2 but were unsuccessful, probably because demux_mt relies on the many mitochondrial reads scattered across the empty and low-quality droplets present in droplet-based datasets. Since high-quality droplets often contain few mitochondrial reads, clustering can rely on information from “empty” droplets. Excluding low-quality droplets from droplet-based sequencing data therefore also risks failed clustering.

Additionally, single-nucleus datasets should generally have lower mitochondrial read coverage than those produced from whole cells, although in our hands, single nucleus ATAC is often suitable for clustering due to the high accessibility of the mitochondrial genome in ATAC-seq assays (Figure 3H). We do not expect single nucleus RNA-seq to be suitable for mitochondrial clustering, owing to uneven coverage of the mitochondrial genome combined with the mitochondrial depletion expected in single nucleus data, and we expect some datasets, especially those produced from cells under severe stress or from degraded post-mortem tissue samples, to be similarly unsuitable.

In all cases, hierarchically clustering mitochondrial haplotypes and comparing the results with identifications from demux_mt should provide a helpful quality control check. cells that receive the same mitochondrial identity but do not appear adjacent in hierarchical clustering were likely identified with very little information, and clustering results should therefore be considered unreliable.

### Ambient RNA analysis

#### Testing ambient expression values against published ambient gene lists

After running CellBouncer’s quant_contam, cellbender^29^, DecontX^28^, and SoupX^27^ on the 10X NSCLC data set^41^, we extracted the ambient gene expression vector from each program. This vector has an entry between 0 and 1 for every gene, with all entries summing to 1, where every entry describes the relative contribution of a gene to the ambient RNA pool.

We then compared each program’s ambient gene expression vector to a set of genes previously reported to comprise a significant portion of ambient RNA in several brain-specific single cell RNA-seq datasets^33^. Although this study focused on different cell types than were present in our test data, we found high concordance between the list of contaminant genes and all inferred ambient expression profiles (Mann-Whitney p < 2.2^-16^ for ambient expression values of genes in and not in the list, for all programs), with CellBouncer’s slightly higher than the other methods (Mann-Whitney U = 18943956 compared to cellbender 18705538, DecontX 18859634, and SoupX 18789108). We note that the source study found ambient RNA removal with cellbender, DecontX, and SoupX to be inadequate in cells of the predominant type; CellBouncer may remedy this limitation by ignoring cell type and gene expression data altogether when inferring contamination.

### Processing custom tags

#### Comparing heterotypic doublet rates implied by alternative programs

As part of our benchmarking strategy for tag-based demultiplexing programs, we created tag count sets from randomly-downsampled CellPlex reads from the 10X NSCLC data^41^, and by removing various percentages of foreground tag counts from each cell to simulate low signal-to-noise (Online Methods). We then assigned cells to tag-based identities using CellBouncer’s demux_tags and alternative programs BFF^21^, deMULTIplex2^22^, demuxmix^23^, GMMdemux^24^, hashedDrops^26^, hashSolo^25^, and HTOdemux^20^.

As a final test of demux_tags and doublet_dragon, we computed the heterotypic doublet rate from the tag-based identities from each program we tested, on each altered version of the NSCLC data we used for benchmarking. Since none of our alterations changed the underlying doublet rate, any variation in heterotypic doublet rate inferred by each program is due to technical noise. To get an expected “true” heterotypic doublet rate, we then subtracted the expected rate of each type of homotypic doublet from the consensus doublet rate inferred using doublet_dragon on genotype-based identities from the full data set. We found that demux_tags inferred heterotypic doublet rates that were both more consistent and closer to the true value than the other programs.

### Demultiplexing inter-species composite cells

#### Assessing genotype demultiplexing accuracy

After collecting and aligning data from interspecies composite cell lines, we tested several alternative tools for identifying cell lines of origin. As ground truth, we collected MULTIseq^50^ tag data that was used to label each cell line of origin. We also collected whole genome sequencing data for each diploid cell line produced for a prior study^34^. We then ran demux_vcf, demuxalot^16^, demuxlet^14^, and Vireo^15^ on composite cells to assess the accuracy of each by comparing with the MULTIseq-based identities.

Since composite cells contain two genomes, successful identifications identify allospecific composite cells as doublets of two cell lines, meaning that we could not identify true doublets using genotype demultiplexing. Because of this, we first discarded all cells identified as doublets by running demux_tags on the MULTIseq data before assessing the accuracy of genotype-based identifications. We also added an option to demux_vcf to provide an allowed list of doublet identities; this limits the set of possible doublet identities a cell can receive and therefore improves accuracy.

Using MULTI-seq labels as ground truth, we found that demux_vcf was more accurate in identifying composite cell lines of origin than other tools, with demuxalot second best but considerably worse on one library, Vireo the least accurate, and demuxlet unable to run on the large SNP set we used (Table S7).

We then used CellBouncer’s quant_contam tool to profile ambient RNA contamination using genotype mismatching; we note that this tool is better suited to detect ambient RNA than alternatives that rely on gene expression heterogeneity, as our data set contained many individuals but only one cell type. We found ambient RNA profiles to be fairly consistent across the two technical replicates (Figure S7) and that CellBouncer’s per-cell contamination rate estimates were closer to mean values we estimated from the discrepancy between inferred bulk proportions and cell line assignments than per-cell contamination rate estimates from cellbender, DecontX, and SoupX (Figure S8, Online Methods). Contamination rate estimates from cellbender^29^ were tightly correlated to the number of genes detected per cell (Figure S9), suggesting a possible pitfall of using data from empty droplets to model ambient RNA.

#### Identifying mitochondrial haplotypes

Although we were able to infer most but not all mitochondrial haplotypes in these cells using hierarchical runs of demux_mt (Figure S10, Figure S11, Figure S12), we further constructed a full mitochondrial haplotype for each contributor line using variant calls from our WGS data.

This allowed us both to recover low-frequency mitochondrial haplotypes and to have knowledge of all SNPs segregating among mitochondrial haplotypes.

We modified demux_mt to count reads specific to each contributor haplotype in every allotetraploid cell, given an identity derived from genotype-based demultiplexing (Online Methods). This run mode outputs allele-specific counts at every SNP that segregates between contributor mitochondrial haplotypes. For identifying mitochondrial haplotypes in inter-species composite cells, we chose to use species-consensus mitochondrial haplotypes consisting only of SNPs fixed and segregating between species; for intra-species allotetraploid composite cells, we used individual-specific mitochondrial haplotypes.

## Supplementary Methods

### Descriptions of algorithms

#### Species demultiplexing (demux_species)

CellBouncer’s demux_species requires as a starting point lists of k-mers unique to each species’ transcriptomes. To accomplish this, it builds k-mer lists specific to a chosen value of k and set of species to be pooled. Our pipeline requires k-mer counts from FastK^70^, which are stored in a custom file format in sorted order. CellBouncer’s program get_unique_kmers takes sorted k-mer counts from FastK and outputs lists of k-mers present in one but not other species’ transcriptomes, removing low-complexity k-mers with DUST score^56^ greater than 2. It can also optionally sample a set number of k-mers from each species, which are chosen in descending order of frequency, randomly choosing those with tied frequencies.

To hold k-mers in memory, we developed a data structure that represents k-mers as 64-bit unsigned integer keys, using the binary encoding scheme A = 00, C = 10, G = 01, T = 11. These are in turn stored in a hash table. This means that any length of k-mer up to 32 bases can fit in one integer and will result in a similar overall memory usage (and execution time). If a k-mer length greater than 32 is chosen, a second 64-bit unsigned integer must be stored per k-mer, with a second layer of hash tables, resulting in a leap in memory usage and execution time. In our tests, k = 32 was more than sufficient to obtain high accuracy.

Because k-mer sets are expensive to store in memory, CellBouncer includes the option to make one pass through the reads per species, in order to decrease memory usage at the cost of higher execution time. In our benchmarking tests, this was unnecessary, but may become necessary if many species are included and memory is limited.

We chose to perform exact cell barcode matching during the k-mer counting step, because fuzzy matching introduces higher memory usage and slows down execution time. In the interest of keeping all relevant data, however, one mismatch is allowed per cell barcode when demultiplexing reads. CellBouncer also corrects any mismatched bases in cell barcodes when writing out demultiplexed reads, in order to speed up downstream processing. When counting k-mers, we also discard counts from reads with duplicate UMIs, but we require UMI matches to be exact in order to collapse them.

Once counts are compiled, demux_species fits a multinomial mixture model to infer the species of origin of each cell, using the expectation-maximization (EM) algorithm. Each species identity is represented by a multinomial distribution over the vector of k-mer counts from each species, and each possible combination of two species (doublet identity) is represented by a distribution whose parameters are the average of the parameters of the two “parent” species’ distributions. To ensure that each doublet distribution continues to model the desired doublet type, we introduce a tweak to the EM algorithm: after each maximization step, each doublet distribution’s parameters are set to the mean of the parameters of its two parent distributions.

After assigning cell barcodes to species of origin, we use a two-component mixture model to infer which barcode sequences are likely to represent empty droplets, and which droplets were likely to have contained cells. For this, we assume that cell-containing droplets will have more transcriptomic reads, and therefore more species-specific k-mer counts as well as more confident species assignments. To reflect this, we use compound distributions whose likelihoods are the product of a Poisson distribution over total species-specific k-mer count per cell and a Gamma distribution over the log likelihood ratio of assignment (assignment LLR) in the same cell. We take cell barcodes whose likelihood under the high-count, high-LLR model is greater than under the low-count, low-LLR model to be putative cell-containing barcodes.

Once cells are assigned to the likeliest species of origin, demux_species makes a final pass through all provided reads to write each to a separate file according to its species of origin, facilitating downstream processing. The assignments used in this step include the full set of droplet barcodes, not just the smaller filtered set of putative cell-containing barcodes found in the last step, providing the user flexibility to incorporate later filtering steps taken after reference genome alignment to remove cells with few or poor-quality transcriptomic reads. Users can provide multiple types of reads for the same library, provided that barcodes are shared, or that a mapping from each RNA-seq barcode to other data type’s barcodes exist. Because genes are under more selective constraint than other genomic regions, the space of potential transcriptomic k-mers is much smaller than that of the genomic k-mers that would be required to demultiplex using ATAC-seq or other genomic reads. Because of this, we rely solely on transcriptomic reads for k-mer counting and species assignment.

#### Demultiplexing using genotype data (demux_vcf)

When a reference panel contains individuals from deeply divergent groups, such as different subspecies or species, not all SNPs are equally useful in discriminating between possible cell identities. This is because many SNPs in the panel will be fixed differences between groups. If there are many such fixed SNPs, then slight differences between individuals in missingness and genotype error at these SNPs can significantly influence the overall rate at which a cell matches the expected alleles from its individual of origin. When this happens, the wrong individual from the correct group can be chosen over the correct individual.

To remedy this situation, we chose a strategy in which a different set of SNPs is used to compare each pair of candidate cell identities. In the case described above, this choice allows all fixed between-group differences to inform comparisons between individuals belonging to different groups, but not comparisons between members of the same group. Our approach assumes individuals to be diploid and considers only biallelic SNPs. For each pair of individuals in the reference panel, we track all SNPs where the two individuals have different genotypes, and we then count reference and alternate alleles observed in each cell at each type of SNP. In other words, if (x,y) denotes SNPs at which individual 1 has x alternate alleles and individual 2 has y alternate alleles, we compile 6 counts of reference and alternate alleles in each cell in order to choose between these individuals, corresponding to SNPs of type (0,1), (0,2), (1,0), (1,2), (2,0), and (2,1). Furthermore, instead of adding 1 for each reference or alternate allele observed in a read, we add the probability of each read being correctly mapped: 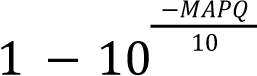. We also skip all reads marked as duplicate, unmapped, or non-primary alignments.

Once all alleles of interest have been counted in all cells, we use each (reference, alternate) allele count pair in each SNP type to compare the likelihood of the cell deriving from the first versus the second individual. To this end, we choose an initial rate at which we expect reference alleles to be misread as alternate alleles, ε_*R*_, and a rate at which we expect alternate alleles to be misread as reference alleles, ε_*A*_. Then, the expected alternate allele matching rate *p* at sites where a cell’s individual of origin is homozygous for the reference allele is ε_*R*_, at heterozygous sites is 0. 5 * (1 − ε_*A*_ + ε_*R*_), and at sites homozygous for the alternate allele is 1 − ε_*A*_. Each pair of (reference, alternate) allele counts can then be assigned an expected alternate allele matching probability given that the cell comes from individual 1, *p*_1_, and another given that the cell comes from individual 2, *p*_2_. For each category of SNPs, then, the reference and alternate allele counts in a cell can be used to produce a binomial log likelihood ratio of the cell deriving from individual 1 versus individual 2. For example, if a cell has *r* reference alleles and *a* alternate alleles observed at SNPs of type (0,1), then the log likelihood ratio of individual 1 versus individual 2 identity is where *lBinom*(*r* + *a, a, ε_R_*) − *lBinom*(*r* + *a, a*, 0.5 * (1 − ε_*A*_ + ε_*A*_)), where *lBinom*(*n, k, p*) is the log binomial likelihood of *k* successes in *n* trials with probability *p*.

This framework allows us to compare directly every pair of possible single-individual identities, but it does not include cell doublets, where two cells enter the same droplet and are assigned the same barcode sequence. Doublets pose a challenge: although this method tracks all types of SNPs necessary to compare the likelihood of a doublet identity to both of its component singlet identities, it does not allow us straightforwardly to compare a doublet identity to any other possible identities. To illustrate this problem, we know that a doublet of individual 1 and 2 has an expected alternate allele fraction of 0.5 * (1 − ε_*A*_ + ε_*R*_) at SNPs of type (2,0) in individuals 1 and 2, since half of the reads should originate from individual 1 and half from individual 2, and we can average the expected alternate allele fractions from both individuals at SNPs of this type. We do not, however, know the expected alternate allele fraction in doublets of this type at sites of type (2,0) in individuals 1 and 3. This is because we have not computed the expected alternate allele fraction in individual 2 across all sites where individual 1 is homozygous for the alternate allele.

To solve this problem, we first compute the log likelihood ratio of each doublet identity versus each of its component singlet identities, using the same sets of SNPs used to compare the two singlet identities. As mentioned above, the expected alternate allele fraction in the doublet is the average of the expected alternate allele fraction in each of the two component identities, or more generally, *f* − *f* * ε_*A*_ + (1 − *f*)* ε_*R*_, where *f* is the fraction of chromosomes possessing the alternative allele. This gives the following expected alternative allele fractions: 0. 25 − 0. 25ε_*A*_ + 0. 75ε_*R*_ for (0,1) or (1,0) SNPs, 0. 5 * (1 − ε_*A*_ + ε_*R*_) for (0,2) or (2,0) SNPs, and 0. 75 − 0. 75ε_*A*_ + 0. 25ε_*R*_ for (1,2) or (2,1) SNPs. We then take advantage of the fact that all computed values are ratios. Suppose that *LL*(*x*) denotes the log likelihood that a cell comes from individual *x*, and *S_1_* denotes a singlet containing a cell from individual 1, *S*_2_ denotes a singlet containing a cell from individual 2, and *D*_12_ denotes a doublet containing a cell from individual 1 and a cell from individual 2. We have already computed *LL*(*D*_12_ − *LL*(*S*_1_) and *LL*(*D*_12_) − *LL*(*S*_2_) as described above. Suppose we also are considering a third individual and have already computed *LL*(*S*_1_) − *LL*(*S*_3_) and *LL*(*S*_2_) − *LL*(*S*_3_). We can then compare the likelihood of the doublet *D*_12_ to the singlet *S*_3_: *LL*(*D*_12_) − *LL*(*S*_3_) = *LL*(*D*_12_) − *LL*(*S*_1_)) + *LL*(*S*_1_) − *LL*(*S*_3_)) = (*LL*(*D*_12_) − *LL*(*S*_2_) − (*LL*(*S*_2_) − *LL*(*S*_3_)). We compute each such log likelihood ratio both ways and take the mean of the two (giving the log of the geometric mean likelihood ratio).

We use the same logic to compare each pair of doublet identities. In cases where the two doublet identities share a member, there is one way to compute the log likelihood ratio, by subtracting the log likelihood ratio of each doublet versus the shared member singlet: *LL*(*D*_12_) − *LL*(*D*_13_) = (*LL*(*D*_12_) − *LL*(*S*_1_) − (*LL*(*D*_13_) − *LL*(*S*_1_). When the two doublets do not share a member, there are four ways to compute the log likelihood ratio, taking advantage of the log likelihood ratios already computed between doublets that share a member:

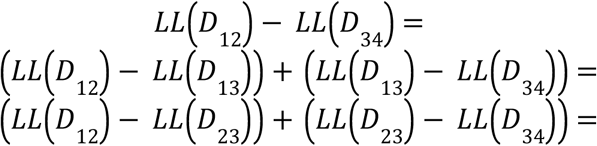

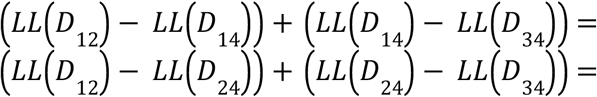

We compute these log likelihoods each way and take the mean of all four estimates.

All log likelihood ratio computations described above leave us with a table of comparisons between each pair of candidate identities, which still needs to be transformed into a single most likely identity. To accomplish this, we find the comparison giving the largest-magnitude log likelihood ratio in the table and eliminate the identity deemed less likely by this comparison (if there are multiple log likelihood ratios of equal magnitude, we eliminate all of their less-likely identities). We then repeat this process until there are only two possible identities left; the likelier of the two identities is taken as the true identity, and the final remaining log likelihood ratio, which describes the degree of preference for the likeliest over the second likeliest identity, is reported as a measure of assignment confidence.

In practice, the table of all pairwise log likelihood ratio comparisons can become very large when there are many individuals in the reference panel, due to the existence of 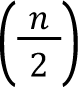 possible doublet combinations when there are *n* individuals. To remedy this, after we compute all log likelihood ratios for singlet/singlet comparisons, we eliminate from consideration all but the ten likeliest individuals. This limits the number of potential doublet identities under consideration to 45 and greatly improves performance when there are many individuals in the genotype reference panel.

At this point, assignments can still be inaccurate if the initial values of ε_*R*_ and ε_*A*_ were inaccurate. This might be the case if many genotype calls were incorrect, or if there is a high amount of ambient RNA contamination. To account for this, we first assign each cell an identity using the initial values of ε_*R*_ and ε_*A*_ and then re-infer these parameters from the data, conditional on the first-round assignments. We accomplish this by maximizing the un-normalized posterior log probability density of each cell’s reference and alternate allele counts under the binomial distribution, using the initial ε_*R*_ and ε_*A*_ values as means of normally distributed priors, with standard deviation values that can be supplied by the user (and decreased to minimize the influence of the data on the posterior estimates). We compute posterior estimates using the L-BFGS algorithm for numeric optimization, with prior estimates as initial guesses. We also constrain both variables to the domain (0,1) by applying the logistic transformation to both, and then maximizing the probability density of the back-transformed variables.

With the maximum *a posteriori* estimates of ε_*R*_ and ε_*A*_, we re-visit each cell and compute a final assignment and log likelihood ratio; these are all written to disk together with some summary statistics and the allele count data, which can then be used to profile ambient RNA by CellBouncer’s quant_contam program.

#### Demultiplexing using mitochondrial haplotype clustering (demux_mt)

The first step of mitochondrial demultiplexing is to locate candidate variant sites on the mitochondrial genome. These will inevitably include some examples of sequencing error or mis-aligned reads in addition to true variant sites. Our method identifies mitochondrial sites where at least two alleles appear in the pileup, takes the two most common alleles at each such site, and sorts these in decreasing order of minor allele frequency (MAF), so that the sites most likely to be true SNPs appear first.

At this step, we then apply an optional minimum coverage threshold. The threshold is determined by creating a histogram of the number of sites that would pass every possible candidate minimum coverage threshold, and then finding a relatively low threshold that would exclude many low-coverage sites by finding the “knee” point of this histogram. We took the knee to be the point of maximum curvature, where curvature is defined by 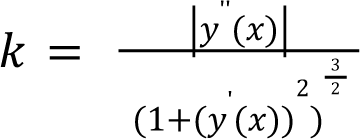. We calculate derivatives using the slopes of line segments between points, assigning the derivative at each point to the mean of the slope of the two adjacent line segments, and then smooth values by averaging each with the value before and after it. We apply the same smoothing to the curvature values. We also require that the chosen filter does not exclude more than 25% of the data.

Next, we read through the BAM and store the major and minor allele count at each candidate SNP seen in reads for every cell barcode, skipping non-primary alignments. Importantly, we include all cell barcodes, even if a list of filtered cell barcodes corresponding to true cells is available. This is because high-quality cell barcodes tend to have few mitochondrial reads, and the mitochondrial reads from droplets uninformative for RNA-seq analyses can still be informative for mitochondrial clustering. Next, we assign each cell barcode the major allele, minor allele, or missing data at each candidate SNP as follows: first, compute

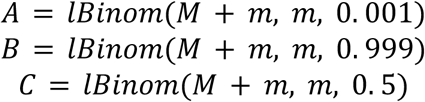

where *M* is the major allele count in the cell, and *m* is the minor allele count in a cell, and *lBinom*(*n, k, p*) is the log binomial likelihood of *k* successes in *n* trials with probability *p*. The cell is assigned the major allele at the site if *A* > *B* & *A* > *C*, the minor allele if *B* > *A* & *B* > *C*, and missing data otherwise (or if *M* = *m* = 0). We also track the sums of *B* and *C* per site, for all cells not assigned the major allele. After all cells are processed, if for any site ∑ *C* ≥ ∑ *B*, then we take this to mean that the overall evidence that the site is erroneous and does not behave like a biallelic SNP outweighs the evidence that it does, and the site is eliminated.

The variable coverage in RNA-seq data also leads to high numbers of SNPs with missing data in individual cells. To confront this issue, we made use of the fact that true mitochondrial clades are often defined by multiple mutations that arose on the same branch of the phylogeny. This manifests in our model as multiple minor alleles that are shared by approximately identical groups of cells (but missing in some cells). We reasoned that if we could share information across these linked sites, effectively collapsing multiple co-varying SNPs into groups, we could share information across cells and overcome some of the challenge posed by missing sites. To accomplish this, when we are comparing two sites and *A* denotes the set of cells with the minor allele at the first site (with higher MAF) and *B* denotes the set of cells with the minor allele at the second site (with lower MAF), we compute the following:

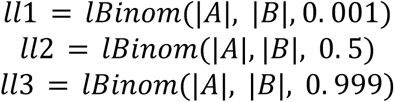

Then, if *ll*1 > *ll*2 & *ll*1 > *ll*3, then the sites have such different MAFs that the second site looks more like an error to the first site than a potential equivalent SNP, and we choose not to collapse the two. Otherwise, we go on to compute

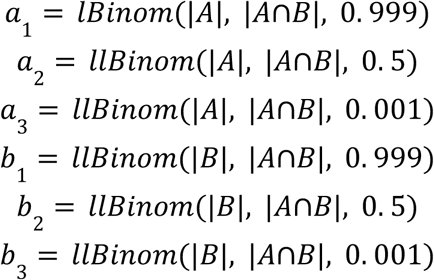

If *a*_1_ > *a*_2_ & *a*_1_ > *a*_3_ and *b*_1_ > *b*_2_ & *b*_1_ > *b*_3_, then we take that as evidence that the number of cells common to both groups is approximately equal to both groups separately, and we collapse the SNPs into a single group. This is done repeatedly until there are no more SNPs to collapse, allowing groups to contain arbitrarily large numbers of SNPs.

Next, we compile two sets for each SNP group: cell barcodes with the major allele and cell barcodes with the minor allele. If any SNP group has zero cell barcodes in one of these two sets, it is eliminated. For those remaining, the size of the set of cell barcodes with the minor allele for the SNP group serves as the minor allele frequency of the SNP group; we sort SNP groups in decreasing order of this set size.

We then iteratively construct haplotypes by including SNP groups in decreasing order of MAF, and inferring the number of true haplotypes at each step, as follows:

> `SET num_true_haps = -1`
>
> `SET haplotypes = ∅`
>
> `FOR group in SNP_groups:`
>
> `SET maj = cells with major allele at SNP group`
>
> `SET min = cells with minor allele at SNP group`
>
> `SET haplotypes_new = ∅`
>
> `IF first SNP group:`
>
> `haplotypes_new = [maj, min]`
>
> `ELSE:`
>
> `FOR haplotype in haplotypes:`
>
> `ADD (haplotype ∩ maj) to haplotypes_new`
>
> `ADD (haplotype ∩ min) to haplotypes_new`
>
> `ENDIF`
>
> `SET haplotypes = haplotypes_new`
>
> `SET num_true_haps_new = CALL compute_num_haps(haplotypes)`
>
> `IF num_true_haps_new < num_true_haps:`
>
> `BREAK`
>
> `ELSE:`
>
> `SET num_true_haps = num_true_haps_new`
>
> `ENDIF`
>
> `ENDFOR`

The program tracks groups of cell barcodes belonging to every potential mitochondrial haplotype, given the current set of SNPs. Haplotypes are only removed from consideration when their membership drops to zero. Since this will result in many of the potential haplotypes under consideration being inauthentic, and having counts of cells that reflect noise rather than true haplotype membership, the compute_num_haps routine is used to determine which cell counts reflect true mitochondrial haplotypes and which reflect noise:

> `ROUTINE compute_num_haps(haplotypes):`
>
> `SET sizes = numbers of cells belonging to each haplotype`
>
> `SORT sizes largest to smallest`
>
> `SET llmax = 0`
>
> `SET ntrue_max = -1`
>
> `FOR num_true from 1 to number of haplotypes:`
>
> `SET mean_true = mean of num_true largest sizes`
>
> `SET mean_false = mean of other sizes`
>
> `SET llsum = 0`
>
> `FOR size in sizes:`
>
> `IF size is among num_true largest sizes:`
>
> `ADD Poisson log likelihood(size, mean_true) to llsum`
>
> `ELSE:`
>
> `ENDFOR`
>
> `ADD Poisson log likelihood(size, mean_false) to llsum`
>
> `ENDIF`
>
> `IF ntrue_max is -1 OR llsum > llmax:`
>
> `SET llmax = llsum`
>
> `SET ntrue_max = num_true`
>
> `ELSE IF ntrue_max is not -1 AND llsum < llmax:`
>
> `BREAK`
>
> `ENDIF`
>
> `END FOR`
>
> `RETURN ntrue_max`
>
> `END ROUTINE`

After running this algorithm, we are left with a set of candidate mitochondrial haplotypes. We define a method similar to that used by demux_vcf for assigning a cell to a mitochondrial haplotype: for each pair of haplotypes, we find all sites that differ between the two haplotypes. For each such site, we take the major allele count *M* and minor allele count *m* in the cell and perform a likelihood ratio test:

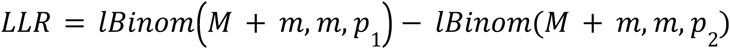

Where *lBinom*(*n*, *k*, *p*) is the log binomial PDF of *k* success in *n* trials with probability *p*, *p*_1_ is the expected minor allele fraction in the first haplotype, and *p*_2_ is the expected minor allele fraction in the second haplotype (0.001 if a haplotype has the major allele and 0.999 if it has the minor, or 0.5 if considering a doublet with both the major and minor allele). We then sum the *LLR* for each such comparison and enter it into a two-way log likelihood ratio table between each candidate set of identities. Our program optionally takes a prior probability of doublet identities, which is incorporated into LLRs between singlet and doublet identities. We iteratively remove the least likely identity from the table until only one comparison remains; this comparison tells us the true identity and the assignment LLR, or confidence in the assignment of that identity.

After obtaining the set of candidate mitochondrial haplotypes that maximizes the number of inferred haplotypes and assigning all cells to their likeliest haplotype, there remains an issue to solve. There is a risk of having added too many SNPs: if an erroneous SNP is added, it can scramble existing mitochondrial haplotypes, shrinking the membership of each and resulting in the compute_num_haps routine inferring a higher number of clusters (the routine assumes that all true clusters will have counts coming from the same distribution; if true cluster sizes shrink to the point where they are similar to the sizes of erroneous clusters, true and erroneous clusters will be counted together as true).

To account for this, we first sum all assignment LLRs, to have a measure of the overall confidence of all cell-haplotype assignments across the data set. We then walk backwards through all added SNP groups, lowest to highest MAF, and for each, exclude the SNP group from all haplotypes, re-assign all cells to the modified haplotypes, and compute the new total summed assignment LLR. We repeat this process while the summed assignment LLR increases; we count as final the set of SNPs and haplotypes that maximizes summed assignment LLR.

Our rationale here is that many erroneous sites will not segregate at the same frequencies as true mitochondrial SNPs (0% or 100% within every cell). Additionally, the inclusion of erroneous sites will “break up” true mitochondrial haplotypes into multiple erroneous haplotypes; each haplotype will only be distinguishable from the other using these erroneous SNPs. This will decrease assignment LLRs: the difference in log likelihood of a cell belonging to one of the other erroneous haplotype will depend only on data from the erroneous site, for which there will be few reads.

#### Refining genotype calls (refine_vcf)

The CellBouncer MT-to-VCF pipeline involves finding variants that segregate among individuals using cell-individual assignments that may be unreliable, *e.g.* when two mitochondrial haplotypes differ by a single segregating site, identifications can hinge on a small number of reads overlapping that site. Additionally, when labeling reads in the BAM by individual of origin prior to variant calling, we only make use of singlet identifications, although doublets can comprise a significant proportion of the data when loading density is high. This can result in some noise in the variant calls that are used to re-identify cells. To address this, we developed a program, utils/refine_vcf, that updates a set of genotypes to maximize the likelihood of the read data and cell-individual assignments. The resulting genotype calls can then be used for another round of assignment using demux_vcf (or any alternative program), which can improve accuracy.

Whereas demux_vcf sums reads per cell across all SNPs of a given allelic state, utils/refine_vcf sums reads across all cells per identity for a given SNP, summing each read’s probability of being correctly mapped 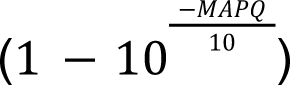 rather than counting each read as 1. Then, for each SNP, it re-infers genotypes by maximum likelihood, using a binomial model and the L-BFGS algorithm.

When re-genotyping a SNP, we infer genotypes for any individuals where the genotype was non-missing in the input SNP set, or where at least 1 total read count was observed either from cells assigned to singlets of the individual, or from cells assigned to at least two different doublet combinations involving the individual. Because L-BFGS is a continuous optimization algorithm, we infer maximum likelihood alternate allele frequencies rather than discrete genotypes. The initial guess for each alternate allele frequency is the genotype in the input file (if non-missing) or the number of alternate alleles divided by alternate plus reference alleles across all identities where the individual is present (both singlets and doublets). We constrain each allele frequency to lie in the range (0,1) by applying the logistic transformation and computing the log likelihood using the back-transformed (expit) values.

We then compute the likelihood of each pair of reference and alternate alleles as *lBinom*(*r* + *a*, *a*, *p*), where *lBinom*(*n*, *k*, *p*) is the log binomial likelihood of *k* successes in *n* trials with probability *p*, *r* is the reference allele count, and *a* is the alternate allele count. If a cell is a singlet, then *p* is the alternate allele frequency in its individual of origin, and if it is a doublet, *p* is the mean of the alternate allele frequencies in the two component individuals. After inferring the maximum likelihood allele frequency in each donor, we convert these to discrete genotypes by rounding to the nearest 0.5 and assigning 0 to homozygous reference, 0.5 to heterozygous, and 1 to homozygous alternate. Only sites for which we infer more than one genotype across all individuals are included in output, and variant quality values in the output data are preserved from the input file.

Because some variant sites reflect sequencing and alignment errors rather than true sequence polymorphism, we also added an optional filtering step to remove sites where the allele count data does not agree well with the maximum likelihood genotypes. After re-inferring all genotypes, we count the total number of expected reference and alternate alleles at the site.

To this end, denoting the expected total alternate allele count as *A* and the expected total reference allele count as *R*, we visit each individual with a non-missing genotype and add (*r* + *a*) * *g* to *A* and (*r* + *a*) * (1 − *g*) to *R*, where *r* is the reference allele count for that individual, *a* is the alternate allele count for that individual, and *g* is the fraction of the individual’s chromosomes expected to have the alternate allele (*g* = 0 for homozygous reference, *g* = 0. 5 for heterozygous, and *g* = 1 for homozygous alternate). We then compute the expected alternate allele fraction at the site *p*, using a pseudocount of 1 to prevent *p* = 0 or *p* = 1: *p* = (*A* + 1)/(*R* + *A* + 2). We then compute the two-tailed binomial p-value of the observed total number of reference alleles at the site, given the total observed reference plus alternate alleles (counting reference and alternate alleles only for individuals with non-missing genotypes) and *p*. Sites with this p-value below a user-supplied threshold are removed from the set of variants.

#### Ambient RNA profiling, quantification, and removal (quant_contam)

CellBouncer’s ambient RNA profiling tool, quant_contam, considers rates at which cells mismatch expected allele frequencies and is thus sensitive to the accuracy of the cell-individual assignments from demux_vcf.

The model used by quant_contam assumes that any individual cell’s reference and alternate allele counts at a given type of SNP are a weighted sum of draws from two binomial distributions: one representing ambient RNA and one representing the cell’s true identity. If *R* is the observed reference allele count and *A* is the observed alternate allele count, then

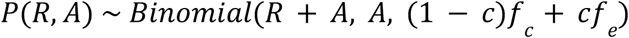

Where *Binomial*(*n*, *k*, *p*) is the binomial distribution with *k* successes in *n* trials with success probability *p*. In this parameterization, *c* is the fraction of the cell’s transcripts originating from ambient RNA, *f_e_* is the expected rate of alternate allele matching at SNPs of a given type in endogenous RNA, conditional on the cell’s identity, and *f_c_* is the rate of alternate allele matching at SNPs of the same type in ambient RNA. Both *f_c_* and *f_e_* are adjusted to reflect sequencing error rates, which are user-adjustable parameters set to 0.001 by default.

To adjust the alternate allele matching frequencies to account for sequencing error, we use the formulae

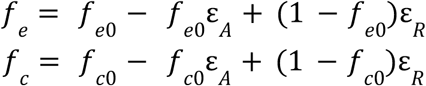

Where ε_*R*_ is the rate at which reference alleles are misread as alternate alleles and ε_*A*_ is the rate at which alternate alleles are misread as reference alleles (both of these error rates are also inferred by demux_vcf as part of the process of assigning cells to individual identities).

The value *f*_*e*0_ is the expected alternate allele frequency at SNPs of a given type, given a cell’s identity, before adjusting for sequencing error. In other words, if a cell comes from donor_1 and *R* and *A* are measured at SNPs where donor_1 is heterozygous, then *f*_*e*0_ = 0. 5. Conversely, if the cell is a doublet of donor_1 and donor_2 and *R* and *A* are measured at SNPs where donor_1 is homozygous for the alternate allele and donor_2 is heterozygous, then *f*_*e*0_ = 0. 25. For cells assigned singlet identities, we consider three types of SNPs: those for which the donor is homozygous for the reference allele, homozygous for the alternate allele, or heterozygous. For doublet cell barcodes, we consider all combinations of SNP types in the two component individuals (*i.e.* 9 types of SNPs based on combinations of three possible allelic states per component individual). Henceforth, we will refer to the types of SNPs relevant to ambient RNA inference when considering a particular cell barcode as the cell barcode’s set of “relevant SNP types.”

The value *f*_*c*0_ is the expected alternate allele frequency in ambient RNA, which is itself a function of each individual’s contribution to the ambient RNA pool. To compute this value, we first visit the genotype data to learn the expected alternate allele frequency in each individual conditional on the allelic state of a SNP in every other individual. This gives us what can be conceptualized as a matrix of conditional alternate allele frequencies *M*, with indices *i*, *j*, and *k*. Indices *i* and *j* together denote types of SNPs (*i* indexes an individual and *j* is a number between 0, 1, and 2 that describes a genotype by denoting the number of alternate alleles in individual *i*) and *k* denotes the individual for which *M_i,j,k_* gives the expected alternate allele frequency at SNPs with the allelic state given by *i* and *j*. Each *M_i,j,k_* is computed directly from the VCF, skipping sites where either individual is missing a called genotype. When *i* = *k*, entries denote expected matching allele frequencies based on the genotype *j*; *e.g. M*_*i*,1,*i*_ = 0. 5 since *j* = 1 denotes heterozygous SNPs (one alternate allele). These values are computed by demux_vcf and written to a file with the extension “.condf.”

We also define a vector *P*, with one entry for each individual in the data set. Each entry *P_*k*_* gives the proportion of ambient RNA originating from individual *k*, with the constraints that 0 < *Pk* < 1 for every individual and (if there are *n* individuals in the data set) 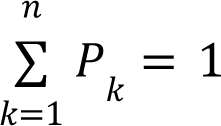 (*P* can be thought of a multinomial distribution over the proportions of individuals in the ambient RNA pool).

With these values, we can compute each value of *f*_*c*0_. For SNPs that have *j* alternate alleles in individual *i*:

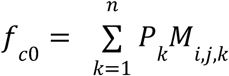

When considering cells with doublet identities, we need to group SNPs by their allelic state in two individuals instead of one. In these cases, for SNPs that have *j* alternate alleles in individual *i* and *y* alternate alleles in individual *x*:

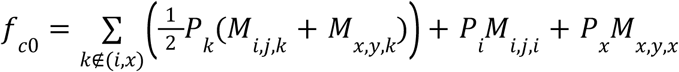

The strategy used by quant_contam is to start with an initial guess of the global contamination rate and infer from it the likeliest mixture of contaminant individuals that could have produced the observed allele matching patterns given the global contamination rate. It then does one round of refinement: it reinfers the global contamination rate conditional on the individual mixture proportions (and resulting alternate allele frequencies in ambient RNA at every type of SNP), then reinfers the mixture proportions conditional on the updated global percent ambient RNA estimate. It then infers the maximum likelihood percent ambient RNA per cell twice, using the first round results as an Empirical Bayes prior. Finally, cell identities are reinferred using the global contamination rate and ambient RNA profile, and this whole process is iterated until the overall log likelihood converges.

At the end of this process, users can provide a gene expression matrix and cell cluster assignments (or the program can use individual identities as clusters), and quant_contam models the expression of every gene in every cell as a mixture of a multinomial distribution representing cluster-specific endogenous gene expression, and another representing gene expression in ambient RNA, with the proportion coming from ambient RNA equal to that cell’s already-inferred percent ambient RNA. Finally, quant_contam subtracts counts expected to come from ambient RNA from the gene expression matrix.

As a first step, CellBouncer must compute an initial guess for the global contamination rate. To this end, we first compute the weighted mean reference and alternate allele count at every type of SNP, given every identity. In other words, for every cell identity in a data set, we compute the mean alternate allele fraction at every type of SNP across all cells for which the SNP type is relevant. When computing means, each cell is weighted by the confidence of its identity (the assignment log likelihood ratio from demux_vcf), divided by the sum of the confidence of all cells assigned to that identity. We then compare each mean observed alternate allele fraction *f_obs_* to its expected value given the cell identity for the SNP type *f_exp_* (for example, for SNPs homozygous in individual 1, *f_exp_* = 0.5 − 0.5ε_*A*_ + 0.5ε_*R*_). Considering our model, *f_obs_* corresponds to 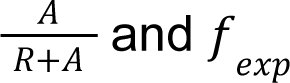 corresponds to *f_e_*, and we wish to estimate *c* without knowledge of *f_c_*. At this stage, for each SNP type, we choose a *f_c_* to get a lower bound on *c*: if *f_obs_* > *f_exp_*, then *c* is minimized when *f_c_* = 1, and if *f_obs_* < *f_exp_*, then *c* is minimized when *f_c_* = 0. Using this idea, we compute *c* using each weighted mean alternate allele frequency at each SNP type as follows: If *f_obs_* > *f_exp_*, then 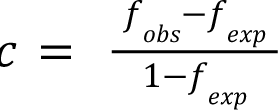, and otherwise 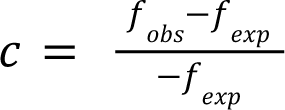.

After computing a minimum *c* this way for each cell, we then compute a weighted mean minimum *c* value for each identity in the data set. We consider these values to reflect both the true, global contamination rate *c* and the makeup of ambient RNA: the mean *c* for an identity depends both on the contamination rate and the proportion of contamination made up of RNA from other identities. In other words, if *c_k_* is the weighted mean minimum *c* estimate for cells assigned identity *k*, *P_k_* is the proportion of ambient RNA originating from identity *i*, and *c* is the true global ambient RNA contamination rate, then for each *k*, *c_k_* = *c*(1 − *P_k_*). By combining these estimates across individuals, we can estimate *c* and each *P_k_*:

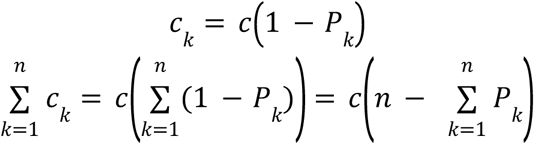

Since 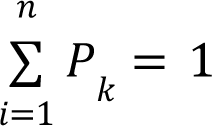 by definition, we can calculate 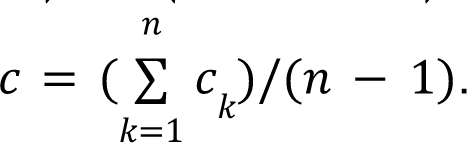. We can then calculate each *P_k_* as *P_k_* = 1 − *c_k_/c*. These estimates serve as initial guesses for the optimization problems detailed below.

We then infer the individual mixture proportions *P* by maximum likelihood, using the initial guesses for each, and the initial guess for *c*. To accomplish this, we consider every cell’s reference and alternate allele count at all relevant SNPs. To handle the constraints on the entries in *P*, we store a vector of logistic-transformed variables: 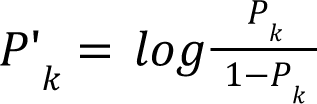 and operate on the back-transformed variables divided by the sum of all back-transformed variables: 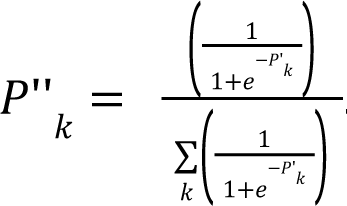. This transformation ensures that every value remains between 0 and 1 and that the sum of all values equals 1. We obtain a maximum likelihood solution using the L-BFGS algorithm^77^ for numeric optimization.

The last time we infer the *P* vector, we bootstrap all observed read count values (100 times by default) and infer *P* in each bootstrap trial, fitting a Dirichlet distribution to all inferred *P* vectors by maximum likelihood. The maximum likelihood Dirichlet concentration parameters are written to disk as the final column in the “.contam_prof” file, which allows us to compute significance when comparing these mixture proportions to any other set of proportions (from cell identities, bulkprop output, or another ambient RNA profile).

Next, we compute from the maximum likelihood *P* entries the value of *f_c_* at every type of SNP, then infer a better estimate of the global contamination rate *c* using these values, using Brent’s method for numeric optimization with first derivative information^107^ with *c* logistic-transformed to constrain it to the range (0,1). We then repeat the last step and re-infer *P* using the improved estimate of *c*.

Next, we infer cell-specific estimates of *c* using the *f_c_* values derived from the maximum-likelihood entries in *P*, using Brent’s method for optimization with first derivative information. After doing this once, we compute the mean and the variance of all cell-specific *c* estimates, fit a beta distribution using the method of moments, and use this as an Empirical Bayes prior in a second round of cell-specific maximum *a posteriori c* inferences, to avoid producing outlier *c* values in cells with low read counts.

With inferred ambient RNA parameters in hand, we then attempt to re-identify cells whose maximum likelihood identity changes with knowledge of the ambient RNA profile. This is achieved using the global *c* estimate and the same procedure and allele counts as in demux_vcf, but adjusting expected alternate allele matching rates using *f_c_* for each SNP type and the global *c*. After re-identifying a cell, we also infer a new maximum *a posteriori c* for that cell, using its new identity and the same beta prior distribution as before. If the likelihood of the cell’s new identity and contamination rate exceeds that of its old identity and contamination rate, the cell’s identity and contamination rate are updated.

Because extensive contamination can significantly bias cell-individual assignments in a way that can alter the global doublet rate (*e.g.* high contamination mostly originating from a single individual might make many cell barcodes falsely appear to be doublets of their true identity and the contaminant individual), we have also built in a way for users to specify an expected fraction of doublets, which is factored into the re-identification process. If the user provides an expected doublet fraction, we first compute the total proportion of cell identities belonging to each individual (singlet identities count twice, and doublet identities count as one of each identity). We name this vector *R*, where *R_i_* is the fraction of all cell identities belonging to individual *i*. We also compute the count of each cell identity in the data set and the total number of cells *t*. If the user-provided expected doublet rate is *d*, then the expected fraction of cells that should be identified as singlets containing cells from individual *i* is 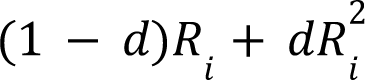 reflecting both true singlets of individual *i* and doublets containing two cells from that individual. Likewise, doublets containing one cell from individual *i* and one from individual *j* should occur at a rate of 2*dR_i_R_j_*. This model is the same used by doublet_dragon when inferring *d*. Denoting the expected rate at which a given identity should occur in the data set as *p*, we can compute a prior likelihood of an identity as *Binomial*(*t*, *n*, *p*) where *n* is the number of cells with that identity. These prior likelihoods are the incorporated into the log likelihood ratio comparisons computed by demux_vcf, where they influence the relative likelihood of choosing one over another identity.

Once cell identities are re-inferred, we compute the log likelihood of the entire data set under the model and inferred parameters. We then iterate all steps described thus far, using the newly re-inferred cell identities and the latest global estimate of *c* in place of the initial guess for *c*, until the log likelihood converges. At this point, quant_contam writes the *P* vector to disk with the extension “.contam_prof,” as well as each estimate of *c* and its Laplace approximation standard error.

If the user also supplies a gene expression matrix, quant_contam can use its inferred per-cell contamination rates and *P* vector to compute an ambient RNA gene expression profile and remove the effect of the ambient gene counts from the matrix. For this procedure, we model every cell’s gene expression as originating from a mixture of two multinomial distributions, similar to DecontX^28^: an endogenous cluster-specific gene expression distribution and another representing ambient RNA. In our case, each cell’s *c* inferred from the allele count data tells us the weight of the ambient distribution versus the endogenous distribution in each cell. We accept external cell-cluster assignments, or in their absence, we assume gene expression to be consistent within each individual and treat cell-individual assignments as clusters

We initialize the ambient expression profile to a weighted average of gene expression in all individuals, with each individual contributing to the ambient profile proportional to its weight in ambient RNA in the *P* vector. We then initialize the endogenous expression profile in each cluster (or individual identity) to the mean across all cells belonging to the cluster or identity. We then jointly infer the fraction of ambient expression originating from each gene, and the fraction of expression in each cluster/identity originating from each gene, by maximizing the multinomial log likelihoods of all distributions, constraining all parameters so that each distribution sums to one as described earlier (in reference to inferring the *P* vector).

After inferring the parameters of the ambient and endogenous gene expression profiles, we write each to disk as a file with the extension “.gex_profile.”

The final step performed by quant_contam is to remove gene counts expected to have come from ambient RNA from the single-cell expression matrix. The sparseness of single-cell expression matrices pose a problem for this task: since many genes inferred to be present in ambient RNA will have zero counts in individual cells, simply multiplying a cell’s total count by the ambient gene expression vector and subtracting these values from the cell’s counts will either result in negative values, or if counts are constrained to be positive, will not remove enough counts to account for the contamination rate inferred through allele mismatching. To get around this, we use the following algorithm to decontaminate each cell’s gene counts in the expression matrix:

> `SET candidate_remove = ∅`
>
> `SET counts_to_remove = cell count total * cell contamination fraction`
>
> `FOR (gene, count) in nonzero gene counts:`
>
> `SET num_remove = ambient gene prop * counts_to_remove`
>
> `SET count_after_removal = count – num_remove`
>
> `ADD (count_after_removal, gene) to candidate_remove`
>
> `ENDFOR`
>
> `SET stop = FALSE`
>
> `WHILE not stop:`
>
> `SORT candidate_remove BY count_after_removal lowest to highest`
>
> `WHILE first count in candidate_remove <= 0:`
>
> `SET (first_count, first_gene) = candidate_remove[0]`
>
> `SET counts_to_remove = counts_to_remove – count of first_gene`
>
> `DELETE candidate_remove[0]`
>
> `ENDWHILE`
>
> `FOR (count_after_removal, gene) in candidate_remove:`
>
> `SET num_remove = ambient gene prop * counts_to_remove`
>
> `SET count_after_removal = gene count – num_remove`
>
> `ENDFOR`
>
> `IF all counts in candidate_remove are positive:`
>
> `SET stop = TRUE`
>
> `ENDIF`
>
> `SET output_expression_matrix = ∅`
>
> `FOR (count_after_removal, gene) in candidate_remove:`
>
> `ADD (gene, count_after_removal) to output_expression_matrix`

We also provide an option, similar to that in SoupX^27^, to round output counts to integers randomly in a way that results in the appropriate number of read counts being removed. In other words, for each count, we take the fractional part of the count above the nearest whole number, and we round up with a probability equal to this number, or down otherwise. This is necessary to achieve the correct number of eliminated reads, *e.g.* if there are many counts of 10.5, simple rounding will change all of these counts to 11, resulting in too many reads in the aggregate, whereas this strategy will randomly round half of these numbers to 10 and half to 11.

#### Inferring bulk proportions (bulkprops)

Our bulk proportion inference tool uses only genotype data for SNPs at which there are no missing individuals. It traverses the input BAM file and counts observed reference (*R*) and alternate alleles (*A*) observed in every cell at these SNPs and uses the following model:

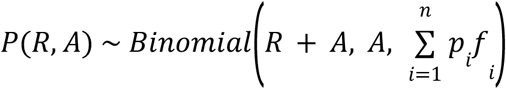

Where *Binomial*(*n*, *k*, *p*) denotes the log binomial probability density function for *k* successes in *n* trials with success probability *p*. In this parameterization, *i* indexes individuals, *n* is the total number of individuals in the data set, and each *p_i_* is an unknown variable that describes the fraction of the reads that originated from individual *i*. Like the *P* vector in our ambient RNA inference tool, each *p_i_* is constrained through variable transformation to lie in the range (0,1) and we require that 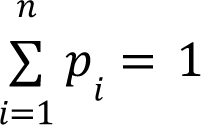. Each *f* is the expected alternate allele frequency at the SNP in individual *i* (which is knowable from the genotype data). Unlike demux_vcf and quant_contam, in which we introduce two sequencing error rates, we model only one error rate here, ε, which is the rate at which any allele is misread as any other. This leads to the following values for *f*_*i*_ depending on the individual’s genotype:

Homozygous for the reference allele: ε

Heterozygous: 0.5

Homozygous for the alternate allele: 1 − ε

Users may provide their own value for ε, or it can be estimated from the data. In the latter case, since our model assumes that all SNPs are biallelic, we track the number of reads that harbor alleles at SNPs that do not match either the reference or alternate allele, *x*, and the total number of reads covering SNPs in the panel, *t*. We treat these non-reference and non-alternate alleles as all being the result of sequencing error and set ε = *x*/*t*. In practice, we find that ε estimated this way is generally on the order of 0.001, which is expected if it mostly describes sequencing error.

Once all allele counts have been compiled, we obtain maximum likelihood estimates for all *p* values using the L-BFGS algorithm. By default, we set initial guesses for all *p_i_* to 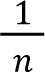, but we provide the option to re-infer maximum likelihood parameters randomly a number of times, using random initial guesses, and to report the highest-likelihood of all these maximum likelihood solutions. This can help avoid reporting local rather than global maximum likelihood solutions.

We also provide the option to bootstrap results by resampling SNPs with replacement and fitting a Dirichlet distribution to the set of all bootstrapped maximum likelihood mixture proportions. This allows us to describe the variance of the maximum likelihood estimates, which in turn allows for the computation of statistical significance of differences between multiple sets of mixture proportions. We achieve this using the same strategy as described in the section about inferring the *P* vector when modeling ambient RNA, and Dirichlet concentration parameters, if inferred, are reported as the final column in the output “.bulkprops” file.

#### Comparing sets of bulk proportions (compare_props.R)

If users wish to compare two sets of individual proportions, we provide an R program called compare_props.R that can find statistically significant differences between two sets of proportions, as long as one has been bootstrapped to infer Dirichlet concentration parameters and can serve as a reference. We allow users to provide cell identifications as a test proportion set, from which mixture proportions are automatically calculated. For each individual in the data set, we measure the probability of the null hypothesis that the individual’s proportion in the “test” (non-reference) set of proportions was generated by the distribution of proportions given by the reference set. To achieve this, we treat the Dirichlet distribution as a set of marginal beta distributions.

If the Dirichlet concentration parameter for individual *i* in the reference proportions is α_*i*_ and there are *n* total individuals, then each individual proportion in the test set 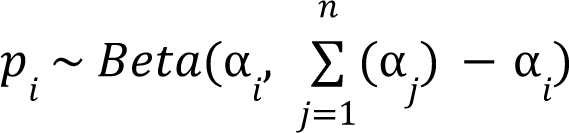. We compute the two-tailed p-value of each *p_i_* using this distribution and then report FDR-corrected p-values for each individual proportion.

We also added an option to run bulkprops “in reverse;” given maximum likelihood mixture proportions from running bulkprops on a full data set, the same program can compute the log likelihood of each input SNP under the consensus mixture proportions using the same model. We also add the option to summarize these log likelihood by gene, mapping reads to genes using the GX and GN tags inserted by aligners like STAR^40^ and reporting mean log likelihood per gene. This can be used as a strategy to pinpoint genes that appear to have donors of origin different from the pool at large, *e.g.* the transcripts that make up ambient RNA. Care must be taken, however, not to overinterpret low log likelihood values reported for genes containing very few informative SNPs.

#### Processing custom tag or CRISPR guide capture sequencing data (demux_tags)

##### Tag counting

Our method counts tags in reads for both sgRNA capture data and HTO data using the Edlib pairwise alignment C++ library^108^, allowing a user-tunable number of mismatches set to 1 by default. For speed, each alignment is set up as a single global alignment between the sequence of interest in a read and a reference of all target sequences concatenated together, padded with runs of N between each sequence. The best match is computed from the position of the best alignment between these sequences. If the input data are sgRNA capture data, demux_tags also allows for variability in the position of the capture sequence in the reverse read by searching the entire read for capture sequences until 100 matches are found. After the 100^th^ match, the most common position in the read at which matches were found is taken to be the position of all future sgRNA capture sequences in reads.

##### Tag assignment

After counting tags in reads, our method fits a two-way mixture model consisting of a negative binomial distribution for the lower/background component and an exponential distribution for the upper component to the un-transformed counts of every tag. We chose an exponential distribution for the upper component of the mixture model based on the observation that means and standard deviations of foreground counts tended to be roughly equal; the high variance of the exponential distribution also reflects counts of foreground tags are sums of two random variables: the true “foreground” count of that tag and the background count of the tag originating from ambient tags.

At this step, we eliminate tags for which both fit distributions appear to represent background or ambient counts. To this end, we first calculate the “separation” between the two distributions for each tag as the cumulative distribution function (CDF) of the negative binomial distribution for the background tags evaluated at the mean of the foreground distribution. This number should be close to 1 for cleanly separated distributions. We then fit a two-component Beta mixture model to all separation values. We compute a similar “separation” metric here: if the CDF of the lower-mean Beta distribution evaluated at the mean of the higher-mean distribution is below 0.01, and the CDF of the higher-mean Beta distribution evaluated at the mean of the lower-mean distribution is above 0.99, then we infer that the distribution separation values are bimodal. This means that tags with low separation values (more likely under the low-mean than the high-mean distribution) likely have no true foreground counts; these tags are then removed. As a final filtering step, we fit a negative binomial distribution to the mean of all background-count distributions that survived filtering, and another negative binomial distribution to the mean of all foreground-count distributions that survived filtering. For any remaining tag, if the mean of its foreground distribution is more likely under the mean background distribution than the mean foreground distribution, then that tag is eliminated as well.

Once the mixture models are fit and filtered, we perform a filtering step for cells, if filtering is enabled or if analyzing sgRNA data. Any cell for which all tag counts are more likely under the background than foreground distribution is considered to be tag-negative and removed from analysis.

After this, we attempt to identify each cell. First, we compute the expected fraction of ambient RNA composed of each tag as 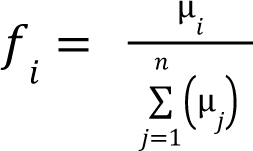, where µ_*i*_ is the background negative binomial distribution mean for tag *i* and *n* is the total number of tags. When identifying a cell, we then compute the fraction of its reads coming from each tag as 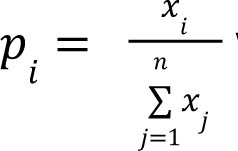 where *x* is the count of tag *i* in the cell. Rather than consider all possible identities for every cell, which would be computationally intractable for high-MOI sgRNA capture data where one cell can be assigned many possible combinations of guides, we sort each tag by decreasing *p_i_* − *f_i_* , prioritizing tags by their degree of enrichment over the background distribution. We then consider the identity of the single most enriched tag, followed by a doublet of the most enriched tag along with the second most enriched tag, and continuing to consider a combination of the three, four, and more most enriched tags in the case of sgRNA data. Because we have sorted identities by decreasing enrichment over background, if we encounter a model with lower likelihood than the previous model considered, we stop looking and accept the model with the highest likelihood encountered thus far. If filtering is enabled or if analyzing sgRNA data, we also include a model representing only background counts; cells for which this is the likeliest model are considered tag-negative.

To build the models that are compared for each cell, we sought to predict expected fractions of all tags under each identity, conditional on the number of observed counts. To accomplish this, we simulate background counts for 1000 “cell” samples by randomly sampling one count from each background distribution for each simulated cell. We also take 1000 samples from each foreground distribution. For any given model, we fit a multinomial linear model to the simulated counts of the form 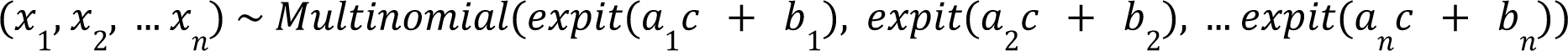, where *c* is the sum of all foreground tag counts for the model, 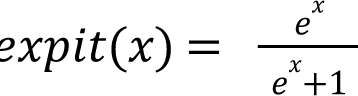, each *a* and *b* are unknown slope and intercept terms in the model, and constraining so that 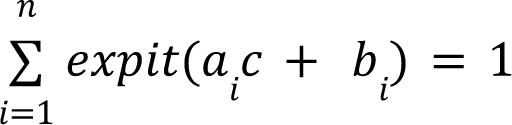. For example, if there are four types of tags Q, R, S, and T, then for the true identity R, we treat each row of (Q, R, S, T) counts as multinomially distributed, with parameters (*p*_1_, *p*_2_, *p*_3_, *p*_4_) that can be calculated as 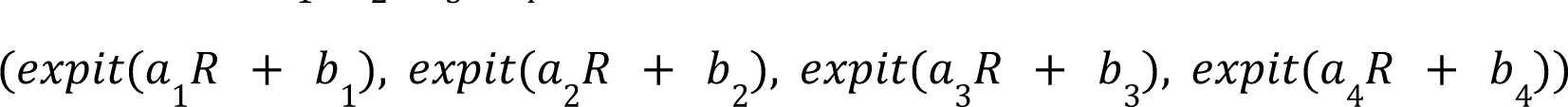. If considering doublets of R and S, then the parameters are determined by 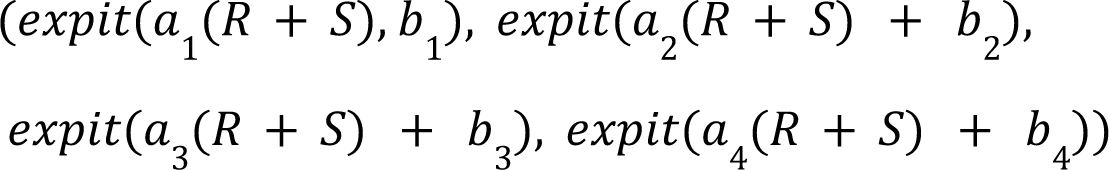.

In the case of sgRNA capture data, the number of possible tags is less restricted, so we adopt a more constrained model that is more computationally feasible. Here, we sum all off-target tag counts into a single background tag count and do not attempt to predict the individual background tag counts using the linear model. In the case of the previous example, when considering the single tag R, we would use the model (*R*, *Q* + *S* + *T*) ∼ *Multinomial*(*expit*(*a*_1_*R* + *b*_1)_, *expit*(*a*_2_*R* + *b*_2_)), and when considering the pair of tags R and S, we would use (*R*, *S*, *Q* + *T*) ∼ *Multinomial*(*expit*(*a*_1_(*R* + *S*) + *b*_1)_, *expit*(*a*_2_(*R* + *S*) + *b*_2)_, *expit*(*a*_3_(*R* + *S*) + *b*_3)_. After predicting the fraction of overall background counts using this model, we then determine each expected individual fraction by multiplying this number by the expected background fractions computed earlier using the means of the fit negative binomial distributions.

Using our simulated counts, we also compute an expectation of foreground counts for the identity being considered by fitting a negative binomial distribution to the simulated foreground counts across all 1000 samples. If we observe tag counts (*Q*, *R*, *S*, *T*) in a given cell, we can then compute the log likelihood of identity *Q* as *lMultinom*((*Q*, *R*, *S*, *T*), (*expit*(*a_1_ Q* + *b*_1_), *expit*(*a_2_ Q* + *b*_2_), + *lNegBinom*(*Q*, µ_*Q*_, ϕ_*Q*_), where *lMultinom* is the log multinomial PDF, *lNegBinom* is the log negative binomial PDF with the µ, ϕ (mean and dispersion) parameterization, each *a* and *b* was inferred by regression as explained in the last step, and µ_*Q*_ and ϕ_*Q*_ came from fitting a negative binomial distribution to 1000 simulated *Q* counts using the method of moments. Conversely, if we are assigning guide RNAs to cells, then the log likelihood would be computed as 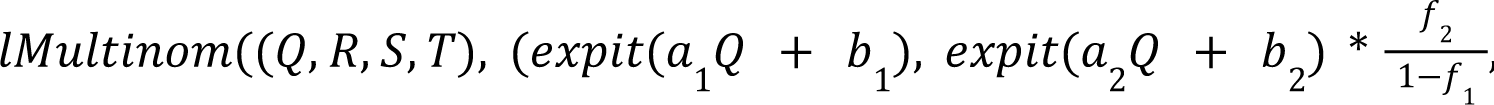, 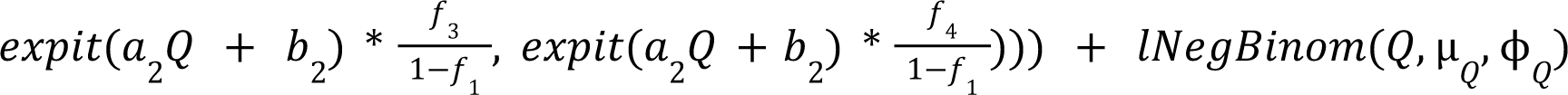, where each *f* is the proportion of all background distribution means coming from a given distribution; these terms are adjusted to account for *f*_1_ being excluded from background due to the first tag (*Q*) being foreground under this model.

As a final step to aid in quality control, we also compute the percent of each cell’s tag counts originating from ambient tags, conditional on the cell’s identity after it has been determined as described above. Here, we define “background” slightly differently: before, each tag count is considered either “background” or “foreground,” meaning that foreground counts are actually the sum of a sample from a true background distribution and a true (but un-modeled) foreground distribution. In this step, we model each count as originating from a weighted sum of two multinomial distributions: one representing foreground and one representing background. We infer the mixing parameter *m* that determines the proportion of counts originating from the background distribution. If a cell has observed counts (*Q*, *R*, *S*, *T*) and was determined to be a singlet of identity *Q*, then we model (*Q*, *R*, *S*, *T*) ∼ *Multinomial*(*mf*_1_ + (1 − *m*), *mf*_2_, *mf*_3_, *mf*_4_), where the *f* vector is the expected background count distribution as described earlier. If the same cell is a doublet of *R* and *S*, we would instead model 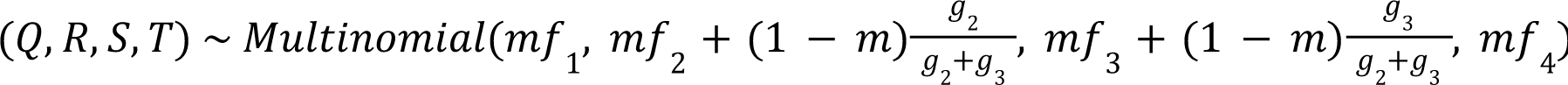, where the *g* vector is the same as the *f* vector but computed using the expectations of foreground exponential distributions, where 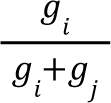 gives the expected fraction of foreground reads originating from tag index *i* in a doublet of tags of index *i* and *j*. If λ is the mean of the foreground exponential distribution for tag index *i*, then 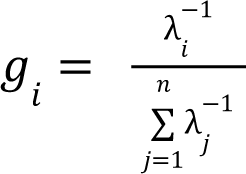. Each *m*, once inferred, is written to a file where these values can be inspected by the user.

### Inferring consensus pool proportions and global doublet rate (doublet_dragon)

We developed the tool doublet_dragon for inferring a consensus doublet rate, which encompasses homotypic as well as heterotypic doublets, from one or more types of CellBouncer-produced cell identifications. Doublet_dragon takes as input one or more CellBouncer-format .assignments files for the same data set; any sets of cell identities that share members are taken to describe the same groups of identities. In other words, if the user has labeled individuals of origin using MULTIseq and is also identifying individuals of origin using genotype data, then the output of demux_tags and demux_vcf should describe the same cell identities. If the cell identities are given the same names in the output files, then doublet_dragon will recognize this and infer consensus proportions of those identities using information from both files. Conversely, if the labels from demux_tags and demux_vcf have different names for cell identities, then doublet_dragon will assume they describe different things (*e.g.* treatments versus cell identities). In this case, information in both files will inform the global doublet rate inference, but two different sets of pool proportions will be inferred. Doublet_dragon places individual identities into groups and infers the proportions within each group separately.

For each group of identities, CellBouncer assumes that the number of cells with each possible identity follows a binomial distribution as follows: for singlets, 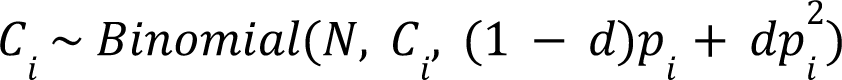 where *Binomial*(*n*, *k*, *p*) is the binomial density of *k* successes in *n* trials with success probability *p*, *C* is the number of cells with singlet identity indexed by *i*, *N* is the total number of cells in the group, *d* is the global doublet rate, and *p* is the probability of sampling identity *i* in the group. Note that our model assumes that cells identified as singlets are actually a combination of true singlets and homotypic doublets, invisible to the CellBouncer program that labeled the cells. For doublets of identity *i* and *j*, the model becomes *C_ij_* ∼ *Binomial*(*N*, *C_ij_*, 2*dp_i_ p_j_*). We then simultaneously infer *d* along with every *p* using L-BFGS. Since initial guesses can influence results by resulting in a local, rather than global, likelihood optimum, we set initial guesses for *d* to a grid of values between 0.05 and 0.95, spaced 0.05 apart, and accept the solution with the highest log likelihood.

Once we have inferred consensus parameters, we also compute the log likelihood of the observed counts in each input file under the consensus model. To this end, we compare the vector of observed counts of each cell identity to the vector of expected proportions under the model (singlets: 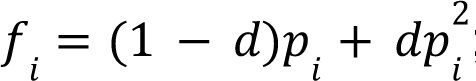; doublets: *f_ij_* = 2*dp_i_ p_j_*) using the multinomial distribution; we report the multinomial log likelihood, which can be useful for comparing likelihoods of different sets of identities computed from the same data.

## Supplementary tables

**Table S1.** Accuracy of demux_species on a labeled data set. Data are from a published study^38^ including cells from three species processed separately (so the true species identities are known). 300 artificial interspecies doublets were also included. Publication-filtered cells refer to cell barcodes that passed quality control after alignment in the published study. CellBouncer-filtered cells refer to cell barcodes that CellBouncer inferred contain real cells based on their species-specific k-mer counts. The “Assignments correct” column includes cell barcodes that do not represent true cells.

| k | k-mers<br>sampled<br>per<br>species | Barcodes<br>assigned | Publication-filtered cells<br>assigned | CellBouncer-filtered cells<br>assigned | Assignments<br>correct | Publication-fi<br>ltered<br>assignments<br>correct | CellBouncer-f<br>iltered<br>assignments<br>correct |
| --- | --- | --- | --- | --- | --- | --- | --- |
| 15 | all | 4499800 | 2889 | 11860 | 29.27% | 30.04% | 30.67 % |
| 20 | all | 1832769 | 2889 | 9798 | 89.91% | 99.97% | 99.86% |
| 25 | all | 1517347 | 2889 | 8117 | 95.24% | 99.97% | 99.84% |
| 35 | all | 1178227 | 2889 | 8479 | 96.87% | 99.97% | 99.86% |
| 45 | all | 901450 | 2889 | 10074 | 97.63% | 99.97% | 99.88% |
| 20 | 1e6 | 945973 | 2889 | 2075 | 98.81% | 100% | 99.95% |
| 20 | 5e6 | 1050689 | 2889 | 2076 | 96.69% | 100% | 99.81% |
| 20 | 10e6 | 1373938 | 2889 | 7989 | 96.01% | 100% | 99.90% |
| 20 | 20e6 | 1373928 | 2889 | 7990 | 96.01% | 100% | 99.90% |

**Table S2.**
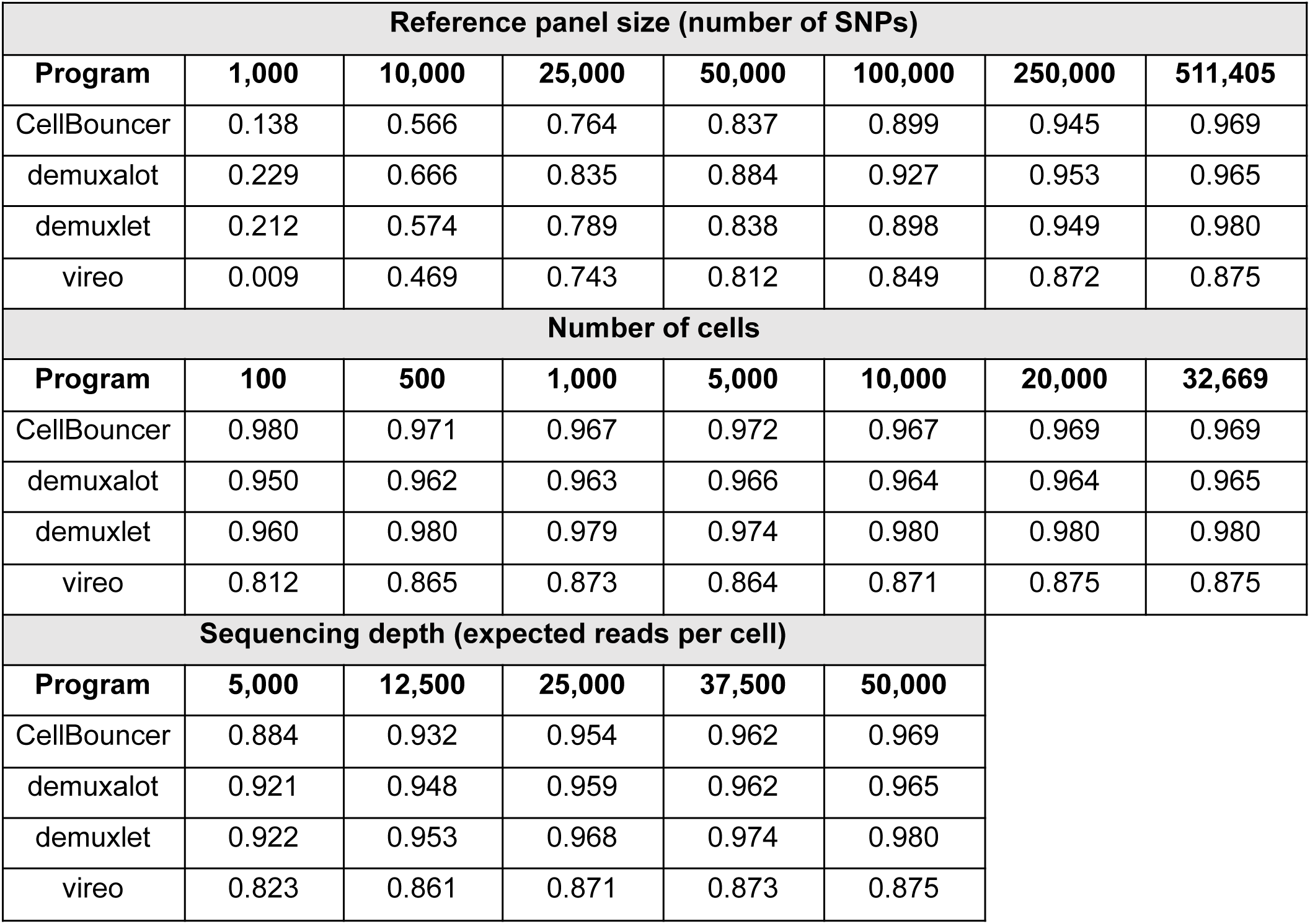
Accuracy (measured by F-score, or the harmonic mean of precision and recall) of CellBouncer against three alternative tools on 10X Genomics 40,000 cell 7-donor NSCLC single-cell RNA-seq data set^41^. Ground truth was the majority vote assignment across all methods plus 10X CellPlex data; ties were broken using CellPlex assignments. Top section: effect of randomly downsampling the genotype data to the reported number of SNPs (right column uses the full panel). Middle section: effect of randomly downsampling the number of cells in the data set (right column uses all cells). Bottom section: effect of randomly downsampling reads to the reported targeted depth per cell (right column uses all reads).

**Table S3.** Precision, recall, F score (harmonic mean of precision and recall), and number of informative variants produced by running several genotype-free individual demultiplexing tools on 10X Genomics 40,000 cell 7-donor NSCLC single-cell RNA-seq data set^41^. Ground truth was the majority vote assignment across all methods plus 10X CellPlex data; ties were broken using CellPlex assignments. Number of variants is the number of output variants that differ among inferred individuals and could be used in downstream analyses. Error meanings: A: crashed (due to memory requirements) or never completed. B: clustering error: two inferred individuals correspond to one actual individual. C: many or all doublets are misassigned.

| Method | Error | Data set | Precision | Recall | F score | Num. variants |
| --- | --- | --- | --- | --- | --- | --- |
| CellBouncer |  | Full | 0.925 | 0.923 | 0.924 | 379,319 |
| CellBouncer + refine_vcf |  | Full | 0.939 | 0.937 | 0.938 | 315,413 |
| freemuxlet | A | Full | N/A | N/A | N/A | N/A |
| scSplit | A | Full | N/A | N/A | N/A | N/A |
| souporcell | B | Full | 0.803 | 0.797 | 0.800 | 927,030 |
| vireo | C | Full | 0.866 | 0.847 | 0.866 | 267,370 |
| CellBouncer |  | 5000 cells | 0.814 | 0.811 | 0.813 | 242,670 |
| CellBouncer + refine_vcf |  | 5000 cells | 0.877 | 0.874 | 0.876 | 175,491 |
| freemuxlet | B | 5000 cells | 0.769 | 0.769 | 0.769 | 148,899 |
| scSplit | B,C | 5000 cells | 0.684 | 0.682 | 0.683 | 193,516 |
| souporcell | B | 5000 cells | 0.792 | 0.782 | 0.787 | 193,869 |
| vireo | C | 5000 cells | 0.866 | 0.850 | 0.858 | 68,411 |
| CellBouncer |  | 2000 cells | 0.868 | 0.865 | 0.866 | 118,994 |
| CellBouncer + refine_vcf |  | 2000 cells | 0.894 | 0.890 | 0.892 | 81,750 |
| freemuxlet | B | 2000 cells | 0.756 | 0.756 | 0.756 | 64,583 |
| scSplit | B,C | 2000 cells | 0.605 | 0.602 | 0.604 | 88,421 |
| souporcell |  | 2000 cells | 0.916 | 0.903 | 0.909 | 67,832 |
| vireo | C | 2000 cells | 0.851 | 0.837 | 0.844 | 23,870 |

**Table S4.** Adjusted p-values for differences of bulk pool proportions from baseline proportions in synthetic contamination experiment. The first two columns denote the individual used as the source of introduced contamination and the percent of contaminant reads introduced to every cell. For each experiment, proportions of each individual were inferred using bulkprops and compared to the baseline proportions, inferred using bulkprops with 100 bootstrap replicates. P-values below a significance threshold of 0.05 are marked with asterisks.

| Source | Fraction | donor_1 | donor_2 | donor_3 | donor_4 | donor_5 | donor_6 | donor_7 |
| --- | --- | --- | --- | --- | --- | --- | --- | --- |
| donor_1 | 0.05 | 7.38E-02 | 4.05E-01 | 4.05E-01 | 4.05E-01 | 4.05E-01 | 4.05E-01 | 4.45E-01 |
| donor_1 | 0.1 | 1.22E-04* | 2.31E-01 | 5.01E-01 | 5.01E-01 | 2.31E-01 | 2.45E-01 | 5.36E-01 |
| donor_1 | 0.15 | 1.04E-08* | 6.93E-02 | 6.22E-01 | 6.22E-01 | 6.93E-02 | 6.93E-02 | 6.82E-01 |
| donor_1 | 0.2 | 1.82E-13* | 5.76E-02 | 9.80E-01 | 9.80E-01 | 8.06E-02 | 3.82E-02* | 9.80E-01 |
| donor_1 | 0.25 | 0.00E+00* | 7.29E-03* | 9.09E-01 | 9.09E-01 | 1.75E-02* | 6.26E-03* | 9.09E-01 |
| donor_2 | 0.05 | 5.90E-01 | 5.90E-01 | 5.90E-01 | 5.90E-01 | 5.90E-01 | 5.90E-01 | 5.90E-01 |
| donor_2 | 0.1 | 7.71E-01 | 2.90E-01 | 7.69E-01 | 7.71E-01 | 2.90E-01 | 4.95E-01 | 7.52E-01 |
| donor_2 | 0.15 | 9.46E-01 | 1.38E-02* | 9.01E-01 | 9.46E-01 | 8.29E-02 | 2.40E-01 | 9.01E-01 |
| donor_2 | 0.2 | 9.79E-01 | 1.44E-04* | 9.79E-01 | 9.79E-01 | 1.03E-02* | 1.33E-01 | 9.79E-01 |
| donor_2 | 0.25 | 8.56E-01 | 2.69E-07* | 8.56E-01 | 8.56E-01 | 5.79E-04* | 5.21E-02 | 8.56E-01 |
| donor_3 | 0.05 | 3.82E-01 | 3.63E-01 | 4.85E-03* | 3.63E-01 | 3.48E-01 | 3.48E-01 | 3.82E-01 |
| donor_3 | 0.1 | 5.38E-01 | 1.29E-01 | 1.53E-08* | 5.23E-01 | 7.54E-02 | 6.18E-02 | 5.38E-01 |
| donor_3 | 0.15 | 7.37E-01 | 3.49E-02* | 0.00E+00* | 6.92E-01 | 8.98E-03* | 2.51E-03* | 7.37E-01 |
| donor_3 | 0.2 | 8.68E-01 | 4.71E-03* | 0.00E+00* | 7.58E-01 | 5.72E-04* | 1.79E-04* | 8.68E-01 |
| donor_3 | 0.25 | 9.32E-01 | 1.72E-03* | 0.00E+00* | 9.32E-01 | 1.16E-04* | 9.71E-05* | 9.32E-01 |
| donor_4 | 0.05 | 4.41E-01 | 4.41E-01 | 4.41E-01 | 2.23E-02* | 4.41E-01 | 4.41E-01 | 4.41E-01 |
| donor_4 | 0.1 | 5.76E-01 | 2.83E-01 | 5.76E-01 | 5.42E-06* | 1.96E-01 | 1.96E-01 | 5.76E-01 |
| donor_4 | 0.15 | 8.32E-01 | 9.16E-02 | 7.95E-01 | 8.82E-12* | 5.36E-02 | 3.17E-02* | 8.32E-01 |
| donor_4 | 0.2 | 9.46E-01 | 1.17E-02* | 9.46E-01 | 0.00E+00* | 5.62E-03* | 3.00E-03* | 9.46E-01 |
| donor_4 | 0.25 | 8.25E-01 | 4.15E-03* | 9.13E-01 | 0.00E+00* | 2.77E-03* | 2.77E-03* | 8.77E-01 |
| donor_5 | 0.05 | 6.70E-01 | 6.61E-01 | 6.61E-01 | 6.61E-01 | 6.61E-01 | 6.61E-01 | 6.61E-01 |
| donor_5 | 0.1 | 9.23E-01 | 4.62E-01 | 9.23E-01 | 9.23E-01 | 1.69E-01 | 4.54E-01 | 9.23E-01 |
| donor_5 | 0.15 | 9.04E-01 | 1.17E-01 | 9.04E-01 | 9.04E-01 | 9.17E-04* | 1.17E-01 | 9.04E-01 |
| donor_5 | 0.2 | 8.94E-01 | 7.88E-02 | 8.94E-01 | 8.94E-01 | 6.37E-07* | 8.04E-03* | 8.94E-01 |
| donor_5 | 0.25 | 6.70E-01 | 2.55E-02* | 6.93E-01 | 6.93E-01 | 2.01E-10* | 8.71E-04* | 6.70E-01 |
| donor_6 | 0.05 | 6.17E-01 | 6.17E-01 | 6.17E-01 | 6.17E-01 | 6.17E-01 | 6.17E-01 | 6.17E-01 |
| donor_6 | 0.1 | 7.60E-01 | 3.51E-01 | 7.60E-01 | 7.60E-01 | 2.50E-01 | 2.50E-01 | 7.60E-01 |
| donor_6 | 0.15 | 9.10E-01 | 7.23E-02 | 8.49E-01 | 8.49E-01 | 3.03E-02* | 6.26E-03* | 9.10E-01 |
| donor_6 | 0.2 | 9.23E-01 | 4.62E-02* | 9.23E-01 | 9.23E-01 | 1.31E-02* | 1.84E-04* | 9.23E-01 |
| donor_6 | 0.25 | 9.52E-01 | 1.29E-02* | 9.52E-01 | 9.52E-01 | 2.35E-03* | 1.07E-06* | 9.52E-01 |
| donor_7 | 0.05 | 4.19E-01 | 2.57E-01 | 3.24E-01 | 3.24E-01 | 2.57E-01 | 2.57E-01 | 4.51E-03* |
| donor_7 | 0.1 | 5.75E-01 | 5.64E-02 | 3.91E-01 | 4.35E-01 | 5.64E-02 | 3.92E-02* | 1.07E-08* |
| donor_7 | 0.15 | 6.92E-01 | 6.42E-03* | 3.99E-01 | 4.83E-01 | 7.60E-03* | 2.69E-03* | 9.33E-15* |
| donor_7 | 0.2 | 9.08E-01 | 2.01E-03* | 7.95E-01 | 9.08E-01 | 3.09E-03* | 8.10E-04* | 0.00E+00* |
| donor_7 | 0.25 | 9.14E-01 | 4.74E-05* | 6.88E-01 | 8.93E-01 | 7.27E-05* | 8.26E-06* | 0.00E+00* |

**Table S5.** F score (harmonic mean of precision and recall) of all HTO-assignment programs compared with CellBouncer demux_tags, on downsampled NSCLC datasets. Three trials were performed at each downsampling level, in which a subset of reads were randomly selected before counting HTO barcodes in reads. True labels came from the majority vote cell identity from running four different genotype-based demultiplexing programs on the same data, and discarding cells for which the majority identification was a tie. The mean, minimum, and maximum F score at each point are reported across the three trials. Results are sorted by decreasing F score within each level of downsampling.

| Mean reads<br>per cell | Program | Mean | Min | Max |
| --- | --- | --- | --- | --- |
| 23 | CellBouncer | 0.8649 | 0.8647 | 0.8652 |
| 23 | GMMdemux | 0.8292 | 0.8285 | 0.8303 |
| 23 | HTOdemux | 0.8291 | 0.8289 | 0.8297 |
| 23 | deMULTiplex2 | 0.7764 | 0.7715 | 0.7848 |
| 23 | demuxmix | 0.7128 | 0.7021 | 0.732 |
| 23 | hashSolo | 0.6807 | 0.6779 | 0.6824 |
| 23 | hashedDrops | 0.4881 | 0.4864 | 0.4913 |
| 57 | CellBouncer | 0.8871 | 0.8869 | 0.8874 |
| 57 | GMMdemux | 0.8839 | 0.8828 | 0.8851 |
| 57 | HTOdemux | 0.8739 | 0.8732 | 0.875 |
| 57 | deMULTiplex2 | 0.8666 | 0.8659 | 0.8674 |
| 57 | demuxmix | 0.8618 | 0.8585 | 0.8671 |
| 57 | hashSolo | 0.8009 | 0.7869 | 0.8087 |
| 57 | hashedDrops | 0.7388 | 0.7378 | 0.7404 |
| 113 | CellBouncer | 0.893 | 0.8924 | 0.8942 |
| 113 | GMMdemux | 0.8927 | 0.8925 | 0.8928 |
| 113 | HTOdemux | 0.8896 | 0.8886 | 0.8911 |
| 113 | deMULTiplex2 | 0.8892 | 0.8887 | 0.89 |
| 113 | demuxmix | 0.8804 | 0.8748 | 0.8836 |
| 113 | hashSolo | 0.8799 | 0.8781 | 0.8813 |
| 113 | hashedDrops | 0.8491 | 0.8484 | 0.8498 |
| 227 | BFF | 0.8959 | 0.8955 | 0.8966 |
| 227 | CellBouncer | 0.8945 | 0.8938 | 0.895 |
| 227 | GMMdemux | 0.8929 | 0.8924 | 0.8937 |
| 227 | HTOdemux | 0.8927 | 0.8918 | 0.8932 |
| 227 | deMULTiplex2 | 0.8834 | 0.8826 | 0.8841 |
| 227 | demuxmix | 0.879 | 0.8786 | 0.8792 |
| 227 | hashSolo | 0.8784 | 0.864 | 0.8875 |
| 227 | hashedDrops | 0.8588 | 0.8575 | 0.8597 |
| 1126 | BFF | 0.8967 | 0.8966 | 0.8968 |
| 1126 | CellBouncer | 0.8936 | 0.8934 | 0.8941 |
| 1126 | GMMdemux | 0.8923 | 0.8918 | 0.8929 |
| 1126 | HTOdemux | 0.8843 | 0.8837 | 0.8847 |
| 1126 | deMULTiplex2 | 0.875 | 0.8741 | 0.8757 |
| 1126 | demuxmix | 0.8737 | 0.8627 | 0.8949 |
| 1126 | hashSolo | 0.8716 | 0.8686 | 0.8753 |
| 1126 | hashedDrops | 0.8441 | 0.8431 | 0.8451 |
| 2235 | BFF | 0.8964 | 0.8962 | 0.8967 |
| 2235 | CellBouncer | 0.8932 | 0.8928 | 0.8937 |
| 2235 | GMMdemux | 0.8924 | 0.8921 | 0.8926 |
| 2235 | HTOdemux | 0.8808 | 0.8803 | 0.8813 |
| 2235 | deMULTiplex2 | 0.875 | 0.8651 | 0.8938 |
| 2235 | demuxmix | 0.8722 | 0.872 | 0.8726 |
| 2235 | hashSolo | 0.8665 | 0.8568 | 0.8729 |
| 2235 | hashedDrops | 0.8007 | 0.8 | 0.8016 |
| 10504 | BFF | 0.896 | 0.896 | 0.8961 |
| 10504 | CellBouncer | 0.8927 | 0.8924 | 0.893 |
| 10504 | GMMdemux | 0.8927 | 0.8924 | 0.893 |
| 10504 | HTOdemux | 0.8767 | 0.8763 | 0.8769 |
| 10504 | deMULTiplex2 | 0.8684 | 0.868 | 0.8687 |
| 10504 | demuxmix | 0.8675 | 0.8674 | 0.8676 |
| 10504 | hashSolo | 0.8519 | 0.8384 | 0.8599 |
| 10504 | hashedDrops | 0.6032 | 0.6028 | 0.6037 |
| 21296 | BFF | 0.8958 | 0.8958 | 0.8958 |
| 21296 | CellBouncer | 0.8927 | 0.8927 | 0.8927 |
| 21296 | GMMdemux | 0.8923 | 0.8923 | 0.8923 |
| 21296 | HTOdemux | 0.8759 | 0.8759 | 0.8759 |
| 21296 | deMULTiplex2 | 0.8674 | 0.8674 | 0.8674 |
| 21296 | demuxmix | 0.8674 | 0.8674 | 0.8674 |
| 21296 | hashSolo | 0.8222 | 0.8222 | 0.8222 |
| 21296 | hashedDrops | 0.4869 | 0.4869 | 0.4869 |

**Table S6.** F score (harmonic mean of precision and recall) of all HTO-assignment programs compared with CellBouncer demux_tags, on decreased signal-to-noise NSCLC data set sets. Three trials were performed at each level, in which a percent of reads from the tag(s) representing the true cell identity were removed (first column). True labels came from the majority vote cell identity from running four different genotype-based demultiplexing programs on the same data, and discarding cells for which the majority identification was a tie. The mean, minimum, and maximum F score at each point are reported across the three trials. Results are sorted by decreasing F score within each level.

| Percent FG removed | Program | Mean | Min | Max |
| --- | --- | --- | --- | --- |
| 0.1 | hashSolo | 0.8941 | 0.8941 | 0.8941 |
| 0.1 | BFF | 0.8929 | 0.8929 | 0.8929 |
| 0.1 | hashedDrops | 0.8897 | 0.8897 | 0.8897 |
| 0.1 | CellBouncer | 0.8896 | 0.8896 | 0.8896 |
| 0.1 | deMULTIplex2 | 0.8759 | 0.8759 | 0.8759 |
| 0.1 | demuxmix | 0.8691 | 0.8691 | 0.8691 |
| 0.1 | HTOdemux | 0.8071 | 0.8071 | 0.8071 |
| 0.1 | GMMdemux | 0.4868 | 0.4868 | 0.4868 |
| 0.25 | BFF | 0.8905 | 0.8905 | 0.8905 |
| 0.25 | hashSolo | 0.8904 | 0.8904 | 0.8904 |
| 0.25 | hashedDrops | 0.8863 | 0.8863 | 0.8863 |
| 0.25 | CellBouncer | 0.8835 | 0.8835 | 0.8835 |
| 0.25 | deMULTIplex2 | 0.8723 | 0.8723 | 0.8723 |
| 0.25 | demuxmix | 0.8654 | 0.8654 | 0.8654 |
| 0.25 | HTOdemux | 0.7997 | 0.7997 | 0.7997 |
| 0.25 | GMMdemux | 0.4838 | 0.4838 | 0.4838 |
| 0.5 | BFF | 0.8899 | 0.8899 | 0.8899 |
| 0.5 | hashSolo | 0.8814 | 0.8814 | 0.8814 |
| 0.5 | hashedDrops | 0.8781 | 0.8781 | 0.8781 |
| 0.5 | CellBouncer | 0.8713 | 0.8713 | 0.8713 |
| 0.5 | deMULTIplex2 | 0.8652 | 0.8652 | 0.8652 |
| 0.5 | demuxmix | 0.8548 | 0.8548 | 0.8548 |
| 0.5 | HTOdemux | 0.7815 | 0.7815 | 0.7815 |
| 0.5 | GMMdemux | 0.4745 | 0.4745 | 0.4745 |
| 0.75 | hashedDrops | 0.8461 | 0.8461 | 0.8461 |
| 0.75 | hashSolo | 0.8402 | 0.8402 | 0.8402 |
| 0.75 | deMULTIplex2 | 0.8377 | 0.8377 | 0.8377 |
| 0.75 | CellBouncer | 0.8284 | 0.8284 | 0.8284 |
| 0.75 | demuxmix | 0.8153 | 0.8153 | 0.8153 |
| 0.75 | BFF | 0.7973 | 0.7973 | 0.7973 |
| 0.75 | HTOdemux | 0.7356 | 0.7356 | 0.7356 |
| 0.75 | GMMdemux | 0.4276 | 0.4275 | 0.4276 |
| 0.9 | CellBouncer | 0.7313 | 0.7313 | 0.7313 |
| 0.9 | hashedDrops | 0.6871 | 0.6871 | 0.6871 |
| 0.9 | hashSolo | 0.6831 | 0.6831 | 0.6831 |
| 0.9 | deMULTIplex2 | 0.6673 | 0.6673 | 0.6673 |
| 0.9 | HTOdemux | 0.548 | 0.548 | 0.548 |
| 0.9 | demuxmix | 0.4981 | 0.4077 | 0.6588 |
| 0.9 | BFF | 0.4106 | 0.4106 | 0.4106 |
| 0.9 | GMMdemux | 0.3256 | 0.3227 | 0.3272 |

**Table S7.** Agreement of MULTIseq labels tagging cell line identity (used as truth) with results of genotype-based demultiplexing of cells from inter- and intra-species tetraploid composite cell lines. Because of genotype demultiplexing tools’ inability to identify droplets containing more than two genomes, cells inferred to be doublets from MULTIseq data were excluded. A1 and A2 are technical replicates of the same library. N/A values indicate that the program did not run in the allowed time frame (12 days).

| Library | Program | Precision | Recall | F |
| --- | --- | --- | --- | --- |
| A1 | CellBouncer | 0.960 | 0.950 | 0.955 |
| A1 | demuxalot | 0.945 | 0.945 | 0.945 |
| A1 | demuxlet | N/A | N/A | N/A |
| A1 | Vireo | 0.260 | 0.214 | 0.235 |
| A2 | CellBouncer | 0.857 | 0.841 | 0.849 |
| A2 | demuxalot | 0.827 | 0.827 | 0.827 |
| A2 | demuxlet | N/A | N/A | N/A |
| A2 | Vireo | 0.240 | 0.190 | 0.212 |
| B | CellBouncer | 0.592 | 0.589 | 0.591 |
| B | demuxalot | 0.577 | 0.577 | 0.577 |
| B | demuxlet | N/A | N/A | N/A |
| B | Vireo | 0.116 | 0.0874 | 0.100 |

**Table S8.** Gene Ontology enrichment results using FUNC^102^, for genes significantly differentially expressed (adjusted p-value < 0.01) in human/chimpanzee composite cells with both human and chimpanzee mitochondria, compared to human/chimpanzee composite cells with only either human or chimpanzee mitochondria. FWER: family-wise error rate. Terms with FWER < 0.05 are shown.

| FWER | GO ID | Term |
| --- | --- | --- |
| 0 | GO:0008219 | cell death |
| 0 | GO:0005737 | cytoplasm |
| 0.002 | GO:0012501 | programmed cell death |
| 0.002 | GO:0006915 | apoptotic process |
| 0.002 | GO:0030330 | DNA damage response, signal transduction by p53 class mediator |
| 0.002 | GO:0005622 | intracellular anatomical structure |
| 0.006 | GO:0070062 | extracellular exosome |
| 0.007 | GO:0033176 | proton-transporting V-type ATPase complex |
| 0.008 | GO:1903561 | extracellular vesicle |
| 0.008 | GO:0043230 | extracellular organelle |
| 0.008 | GO:0065010 | extracellular membrane-bounded organelle |
| 0.009 | GO:0072331 | signal transduction by p53 class mediator |
| 0.013 | GO:0010941 | regulation of cell death |
| 0.013 | GO:0043067 | regulation of programmed cell death |
| 0.014 | GO:0042770 | signal transduction in response to DNA damage |
| 0.02 | GO:0042981 | regulation of apoptotic process |
| 0.027 | GO:0033179 | proton-transporting V-type ATPase, V0 domain |
| 0.029 | GO:0043226 | organelle |
| 0.039 | GO:0031982 | vesicle |
| 0.044 | GO:0009894 | regulation of catabolic process |
| 0.047 | GO:0072332 | intrinsic apoptotic signaling pathway by p53 class mediator |
| 0.048 | GO:0031329 | regulation of cellular catabolic process |

**Table S9.** Gene Ontology enrichment results using FUNC^102^, for genes significantly differentially expressed (adjusted p-value < 0.01) in chimpanzee/bonobo composite cells with both chimpanzee and bonobo mitochondria, compared to chimpanzee/bonobo composite cells with only bonobo mitochondria. FWER: family-wise error rate. Terms with FWER < 0.05 are shown.

| FWER | GO ID | term |
| --- | --- | --- |
| 0 | GO:0031974 | membrane-enclosed lumen |
| 0 | GO:0043233 | organelle lumen |
| 0 | GO:0070013 | intracellular organelle lumen |
| 0 | GO:0003723 | RNA binding |
| 0 | GO:0031981 | nuclear lumen |
| 0 | GO:0005654 | nucleoplasm |
| 0 | GO:0005634 | nucleus |
| 0 | GO:0003676 | nucleic acid binding |
| 0 | GO:0043231 | intracellular membrane-bounded organelle |
| 0 | GO:0043229 | intracellular organelle |
| 0 | GO:0043227 | membrane-bounded organelle |
| 0 | GO:0034641 | cellular nitrogen compound metabolic process |
| 0 | GO:0032991 | protein-containing complex |
| 0 | GO:1990904 | ribonucleoprotein complex |
| 0 | GO:0005622 | intracellular anatomical structure |
| 0 | GO:0043226 | organelle |
| 0 | GO:0006139 | nucleobase-containing compound metabolic process |
| 0 | GO:0044237 | cellular metabolic process |
| 0 | GO:1901363 | heterocyclic compound binding |
| 0 | GO:0097159 | organic cyclic compound binding |
| 0 | GO:0046483 | heterocycle metabolic process |
| 0 | GO:0006725 | cellular aromatic compound metabolic process |
| 0 | GO:1901360 | organic cyclic compound metabolic process |
| 0 | GO:0006807 | nitrogen compound metabolic process |
| 0 | GO:0140513 | nuclear protein-containing complex |
| 0 | GO:0043170 | macromolecule metabolic process |
| 0 | GO:0031967 | organelle envelope |
| 0 | GO:0031975 | envelope |
| 0 | GO:0006396 | RNA processing |
| 0 | GO:0090304 | nucleic acid metabolic process |
| 0 | GO:0098798 | mitochondrial protein-containing complex |
| 0 | GO:0008152 | metabolic process |
| 0 | GO:0010467 | gene expression |
| 0 | GO:0016071 | mRNA metabolic process |
| 0 | GO:0003729 | mRNA binding |
| 0 | GO:0071704 | organic substance metabolic process |
| 0 | GO:0000375 | RNA splicing, via transesterification reactions |
| 0 | GO:0044238 | primary metabolic process |
| 0 | GO:0044271 | cellular nitrogen compound biosynthetic process |
| 0 | GO:0008380 | RNA splicing |
| 0 | GO:0000377 | RNA splicing, via transesterification reactions with bulged adenosine as nucleophile |
| 0 | GO:0000398 | mRNA splicing, via spliceosome |
| 0 | GO:0006397 | mRNA processing |
| 0 | GO:0044260 | cellular macromolecule metabolic process |
| 0 | GO:1901576 | organic substance biosynthetic process |
| 0 | GO:0009058 | biosynthetic process |
| 0 | GO:0044249 | cellular biosynthetic process |
| 0 | GO:0009059 | macromolecule biosynthetic process |
| 0 | GO:0034645 | cellular macromolecule biosynthetic process |
| 0 | GO:0051276 | chromosome organization |
| 0 | GO:0043232 | intracellular non-membrane-bounded organelle |
| 0 | GO:0016070 | RNA metabolic process |
| 0 | GO:0043228 | non-membrane-bounded organelle |
| 0 | GO:0005730 | nucleolus |
| 0 | GO:0015980 | energy derivation by oxidation of organic compounds |
| 0 | GO:0098800 | inner mitochondrial membrane protein complex |
| 0 | GO:0071840 | cellular component organization or biogenesis |
| 0 | GO:0045333 | cellular respiration |
| 0 | GO:0022613 | ribonucleoprotein complex biogenesis |
| 0 | GO:0005740 | mitochondrial envelope |
| 0 | GO:0005739 | mitochondrion |
| 0 | GO:0060255 | regulation of macromolecule metabolic process |
| 0 | GO:0006119 | oxidative phosphorylation |
| 0 | GO:1901566 | organonitrogen compound biosynthetic process |
| 0 | GO:0051171 | regulation of nitrogen compound metabolic process |
| 0 | GO:0034654 | nucleobase-containing compound biosynthetic process |
| 0 | GO:0006996 | organelle organization |
| 0 | GO:0018130 | heterocycle biosynthetic process |
| 0 | GO:0005737 | cytoplasm |
| 0 | GO:0051172 | negative regulation of nitrogen compound metabolic process |
| 0 | GO:0006259 | DNA metabolic process |
| 0 | GO:0019438 | aromatic compound biosynthetic process |
| 0 | GO:0031324 | negative regulation of cellular metabolic process |
| 0 | GO:0016043 | cellular component organization |
| 0 | GO:0080090 | regulation of primary metabolic process |
| 0 | GO:0010629 | negative regulation of gene expression |
| 0 | GO:0005681 | spliceosomal complex |
| 0 | GO:0022904 | respiratory electron transport chain |
| 0 | GO:0005515 | protein binding |
| 0 | GO:1903311 | regulation of mRNA metabolic process |
| 0 | GO:0019222 | regulation of metabolic process |
| 0 | GO:0005743 | mitochondrial inner membrane |
| 0 | GO:1901362 | organic cyclic compound biosynthetic process |
| 0 | GO:0010604 | positive regulation of macromolecule metabolic process |
| 0 | GO:0006412 | translation |
| 0 | GO:0031323 | regulation of cellular metabolic process |
| 0 | GO:0046034 | ATP metabolic process |
| 0 | GO:0009060 | aerobic respiration |
| 0 | GO:0009890 | negative regulation of biosynthetic process |
| 0 | GO:0005694 | chromosome |
| 0 | GO:0006091 | generation of precursor metabolites and energy |
| 0 | GO:0031327 | negative regulation of cellular biosynthetic process |
| 0 | GO:1901293 | nucleoside phosphate biosynthetic process |
| 0 | GO:1902494 | catalytic complex |
| 0 | GO:0006281 | DNA repair |
| 0 | GO:0043043 | peptide biosynthetic process |
| 0 | GO:2000113 | negative regulation of cellular macromolecule biosynthetic process |
| 0 | GO:0009141 | nucleoside triphosphate metabolic process |
| 0 | GO:0010556 | regulation of macromolecule biosynthetic process |
| 0 | GO:0003682 | chromatin binding |
| 0 | GO:0019219 | regulation of nucleobase-containing compound metabolic process |
| 0 | GO:0009206 | purine ribonucleoside triphosphate biosynthetic process |
| 0 | GO:0043933 | protein-containing complex subunit organization |
| 0 | GO:0010558 | negative regulation of macromolecule biosynthetic process |
| 0 | GO:0009145 | purine nucleoside triphosphate biosynthetic process |
| 0 | GO:2000112 | regulation of cellular macromolecule biosynthetic process |
| 0 | GO:0006518 | peptide metabolic process |
| 0 | GO:0043021 | ribonucleoprotein complex binding |
| 0 | GO:0031966 | mitochondrial membrane |
| 0 | GO:0005753 | mitochondrial proton-transporting ATP synthase complex |
| 0 | GO:0019866 | organelle inner membrane |
| 0 | GO:0045259 | proton-transporting ATP synthase complex |
| 0 | GO:0000228 | nuclear chromosome |
| 0 | GO:0030880 | RNA polymerase complex |
| 0 | GO:0030684 | preribosome |
| 0 | GO:0005635 | nuclear envelope |
| 0 | GO:0005829 | cytosol |
| 0 | GO:0032040 | small-subunit processome |
| 0 | GO:0098687 | chromosomal region |
| 0.001 | GO:0010468 | regulation of gene expression |
| 0.001 | GO:0051252 | regulation of RNA metabolic process |
| 0.001 | GO:0006360 | transcription by RNA polymerase I |
| 0.001 | GO:0009201 | ribonucleoside triphosphate biosynthetic process |
| 0.001 | GO:0009889 | regulation of biosynthetic process |
| 0.001 | GO:0043604 | amide biosynthetic process |
| 0.001 | GO:0071103 | DNA conformation change |
| 0.001 | GO:0006974 | cellular response to DNA damage stimulus |
| 0.001 | GO:0003730 | mRNA 3'-UTR binding |
| 0.001 | GO:0042393 | histone binding |
| 0.001 | GO:0010605 | negative regulation of macromolecule metabolic process |
| 0.001 | GO:0009893 | positive regulation of metabolic process |
| 0.001 | GO:0009165 | nucleotide biosynthetic process |
| 0.001 | GO:0006754 | ATP biosynthetic process |
| 0.001 | GO:0006325 | chromatin organization |
| 0.001 | GO:0051173 | positive regulation of nitrogen compound metabolic process |
| 0.001 | GO:0044085 | cellular component biogenesis |
| 0.001 | GO:0009892 | negative regulation of metabolic process |
| 0.001 | GO:0031326 | regulation of cellular biosynthetic process |
| 0.001 | GO:0045934 | negative regulation of nucleobase-containing compound metabolic process |
| 0.001 | GO:0007049 | cell cycle |
| 0.001 | GO:0006839 | mitochondrial transport |
| 0.001 | GO:0009142 | nucleoside triphosphate biosynthetic process |
| 0.001 | GO:0009205 | purine ribonucleoside triphosphate metabolic process |
| 0.001 | GO:0033554 | cellular response to stress |
| 0.001 | GO:0044267 | cellular protein metabolic process |
| 0.001 | GO:0051253 | negative regulation of RNA metabolic process |
| 0.001 | GO:0045935 | positive regulation of nucleobase-containing compound metabolic process |
| 0.001 | GO:0006122 | mitochondrial electron transport, ubiquinol to cytochrome c |
| 0.001 | GO:0009144 | purine nucleoside triphosphate metabolic process |
| 0.001 | GO:0009199 | ribonucleoside triphosphate metabolic process |
| 0.001 | GO:0017148 | negative regulation of translation |
| 0.001 | GO:0006356 | regulation of transcription by RNA polymerase I |
| 0.001 | GO:0006403 | RNA localization |
| 0.001 | GO:0065003 | protein-containing complex assembly |
| 0.001 | GO:0051028 | mRNA transport |
| 0.001 | GO:0080135 | regulation of cellular response to stress |
| 0.001 | GO:0007005 | mitochondrion organization |
| 0.001 | GO:0045943 | positive regulation of transcription by RNA polymerase I |
| 0.001 | GO:0015985 | energy coupled proton transport, down electrochemical gradient |
| 0.001 | GO:0015986 | ATP synthesis coupled proton transport |
| 0.001 | GO:0016591 | RNA polymerase II, holoenzyme |
| 0.001 | GO:0055029 | nuclear DNA-directed RNA polymerase complex |
| 0.002 | GO:0043484 | regulation of RNA splicing |
| 0.002 | GO:0031325 | positive regulation of cellular metabolic process |
| 0.003 | GO:0042773 | ATP synthesis coupled electron transport |
| 0.003 | GO:0042775 | mitochondrial ATP synthesis coupled electron transport |
| 0.003 | GO:0006260 | DNA replication |
| 0.003 | GO:0009152 | purine ribonucleotide biosynthetic process |
| 0.003 | GO:0009891 | positive regulation of biosynthetic process |
| 0.003 | GO:0022618 | ribonucleoprotein complex assembly |
| 0.003 | GO:0010557 | positive regulation of macromolecule biosynthetic process |
| 0.003 | GO:0005687 | U4 snRNP |
| 0.003 | GO:0033018 | sarcoplasmic reticulum lumen |
| 0.003 | GO:0005685 | U1 snRNP |
| 0.003 | GO:0000428 | DNA-directed RNA polymerase complex |
| 0.004 | GO:0050657 | nucleic acid transport |
| 0.004 | GO:0050658 | RNA transport |
| 0.004 | GO:1901564 | organonitrogen compound metabolic process |
| 0.004 | GO:0000278 | mitotic cell cycle |
| 0.004 | GO:0036261 | 7-methylguanosine cap hypermethylation |
| 0.004 | GO:0006457 | protein folding |
| 0.004 | GO:0051236 | establishment of RNA localization |
| 0.004 | GO:0016604 | nuclear body |
| 0.004 | GO:0071013 | catalytic step 2 spliceosome |
| 0.005 | GO:0071826 | ribonucleoprotein complex subunit organization |
| 0.005 | GO:0022900 | electron transport chain |
| 0.005 | GO:0043603 | cellular amide metabolic process |
| 0.005 | GO:0051254 | positive regulation of RNA metabolic process |
| 0.005 | GO:0034622 | cellular protein-containing complex assembly |
| 0.005 | GO:0006164 | purine nucleotide biosynthetic process |
| 0.005 | GO:0034249 | negative regulation of cellular amide metabolic process |
| 0.005 | GO:0009260 | ribonucleotide biosynthetic process |
| 0.005 | GO:0055086 | nucleobase-containing small molecule metabolic process |
| 0.01 | GO:0006753 | nucleoside phosphate metabolic process |
| 0.011 | GO:0051168 | nuclear export |
| 0.011 | GO:0005759 | mitochondrial matrix |
| 0.014 | GO:0061980 | regulatory RNA binding |
| 0.014 | GO:0034709 | methylosome |
| 0.014 | GO:0044615 | nuclear pore nuclear basket |
| 0.014 | GO:0045263 | proton-transporting ATP synthase complex, coupling factor F(o) |
| 0.015 | GO:0010608 | posttranscriptional regulation of gene expression |
| 0.015 | GO:0051726 | regulation of cell cycle |
| 0.015 | GO:0019646 | aerobic electron transport chain |
| 0.017 | GO:0006417 | regulation of translation |
| 0.017 | GO:0046390 | ribose phosphate biosynthetic process |
| 0.017 | GO:1903047 | mitotic cell cycle process |
| 0.017 | GO:0045296 | cadherin binding |
| 0.018 | GO:0072522 | purine-containing compound biosynthetic process |
| 0.02 | GO:0003677 | DNA binding |
| 0.022 | GO:0048024 | regulation of mRNA splicing, via spliceosome |
| 0.022 | GO:0042254 | ribosome biogenesis |
| 0.022 | GO:0031328 | positive regulation of cellular biosynthetic process |
| 0.024 | GO:1990234 | transferase complex |
| 0.025 | GO:0044391 | ribosomal subunit |
| 0.027 | GO:0008134 | transcription factor binding |
| 0.028 | GO:0050684 | regulation of mRNA processing |
| 0.028 | GO:0005488 | binding |
| 0.029 | GO:0009117 | nucleotide metabolic process |
| 0.03 | GO:0005689 | U12-type spliceosomal complex |
| 0.031 | GO:0034655 | nucleobase-containing compound catabolic process |
| 0.031 | GO:0006401 | RNA catabolic process |
| 0.031 | GO:0034248 | regulation of cellular amide metabolic process |
| 0.034 | GO:0045893 | positive regulation of transcription, DNA-templated |
| 0.034 | GO:1903508 | positive regulation of nucleic acid-templated transcription |
| 0.034 | GO:0032269 | negative regulation of cellular protein metabolic process |
| 0.034 | GO:1902680 | positive regulation of RNA biosynthetic process |
| 0.036 | GO:2000767 | positive regulation of cytoplasmic translation |
| 0.039 | GO:0032268 | regulation of cellular protein metabolic process |
| 0.039 | GO:0005683 | U7 snRNP |
| 0.039 | GO:0070937 | CRD-mediated mRNA stability complex |
| 0.039 | GO:0061695 | transferase complex, transferring phosphorus-containing groups |
| 0.04 | GO:0000793 | condensed chromosome |
| 0.041 | GO:0036464 | cytoplasmic ribonucleoprotein granule |
| 0.041 | GO:0071007 | U2-type catalytic step 2 spliceosome |
| 0.041 | GO:0005643 | nuclear pore |
| 0.044 | GO:0034719 | SMN-Sm protein complex |
| 0.045 | GO:1990204 | oxidoreductase complex |
| 0.045 | GO:0010494 | cytoplasmic stress granule |
| 0.045 | GO:0016469 | proton-transporting two-sector ATPase complex |
| 0.047 | GO:0003712 | transcription coregulator activity |
| 0.047 | GO:0005684 | U2-type spliceosomal complex |
| 0.049 | GO:0005840 | ribosome |

**Table S10.** Gene Ontology enrichment results using FUNC^102^, for genes significantly differentially expressed (adjusted p-value < 0.01) in human/chimp composite cells with both species’ mitochondria relative to those with one species’ mitochondria, and in chimpanzee/bonobo composite cells with both chimpanzee and bonobo mitochondria relative to chimpanzee/bonobo composite cells with only bonobo mitochondria. FWER: family-wise error rate. Terms with FWER < 0.05 are shown.

| FWER | GO ID | term |
| --- | --- | --- |
| 0.013 | GO:0071456 | cellular response to hypoxia |
| 0.017 | GO:0036294 | cellular response to decreased oxygen levels |
| 0.019 | GO:0006977 | DNA damage response, signal transduction by p53 class mediator resulting in cell cycle arrest |
| 0.019 | GO:0071453 | cellular response to oxygen levels |
| 0.036 | GO:0002039 | p53 binding |
| 0.04 | GO:0031571 | mitotic G1 DNA damage checkpoint signaling |
| 0.04 | GO:0044819 | mitotic G1/S transition checkpoint signaling |

**Table S11.** Log2 fold change values from differential expression analysis using DESeq2^101^, comparing interspecies composite cells with both species’ mitochondria to those with only one. One cell class (chimpanzee/bonobo cells with chimpanzee mitochondria) was excluded because there were fewer than 10 cells identified using at least 100 informative reads. Shown are genes for which all three comparisons are significant. Gene ontology enrichments for this list of genes are shown in Table S10.

| gene | Chimp/Bonobo both vs bonobo | Human/Chimp both vs chimp | Human/Chimp both vs human |
| --- | --- | --- | --- |
| ANP32E | -0.8244367 | -0.5725004 | -0.5230313 |
| ENSG00000226965 | -3.428982 | 2.93594197 | 2.35599863 |
| ENSG00000251095 | -2.8927968 | 2.36780417 | 2.18120155 |
| ENSG00000287238 | 3.19613466 | 3.08328448 | 3.05689101 |
| FDXR | 1.71254138 | 1.32150504 | 1.31778909 |
| MDM2 | -2.7234912 | 3.01242552 | 2.72662556 |
| MORC4 | -6.1259156 | 2.82405744 | 3.69221881 |
| PLK3 | 3.39906391 | 2.75730583 | 3.45056284 |
| SNRPB | -1.0106242 | -0.4982487 | -0.5311065 |
| TIGAR | 1.27130244 | 1.51769162 | 1.15763434 |

## Supplementary figures

**Figure S1.**
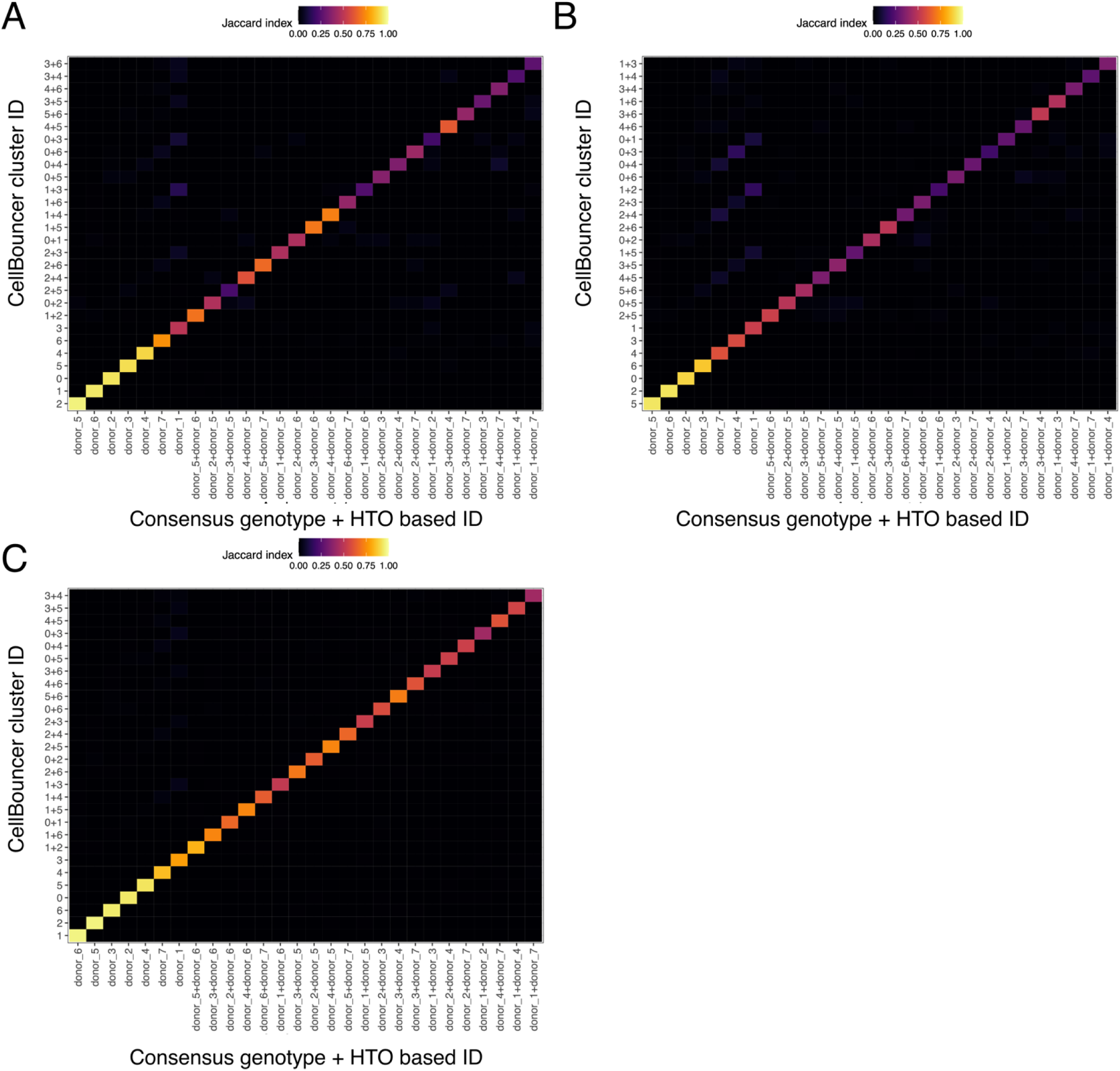
Comparison of cell labels from consensus genotype and CellPlex-based assignments (X-axis) vs inferred assignments without prior genotype data, using CellBouncer’s MT-to-VCF pipeline. Color denotes Jaccard similarity between the group of cells with the label on the X and the Y axis; doublet identities are named as component singlet identities separated by “+”. A: 2,000 cell downsampled data set. B: 5,000 cell downsampled data set. C: full data set (∼33,000 cells).

**Figure S2.**
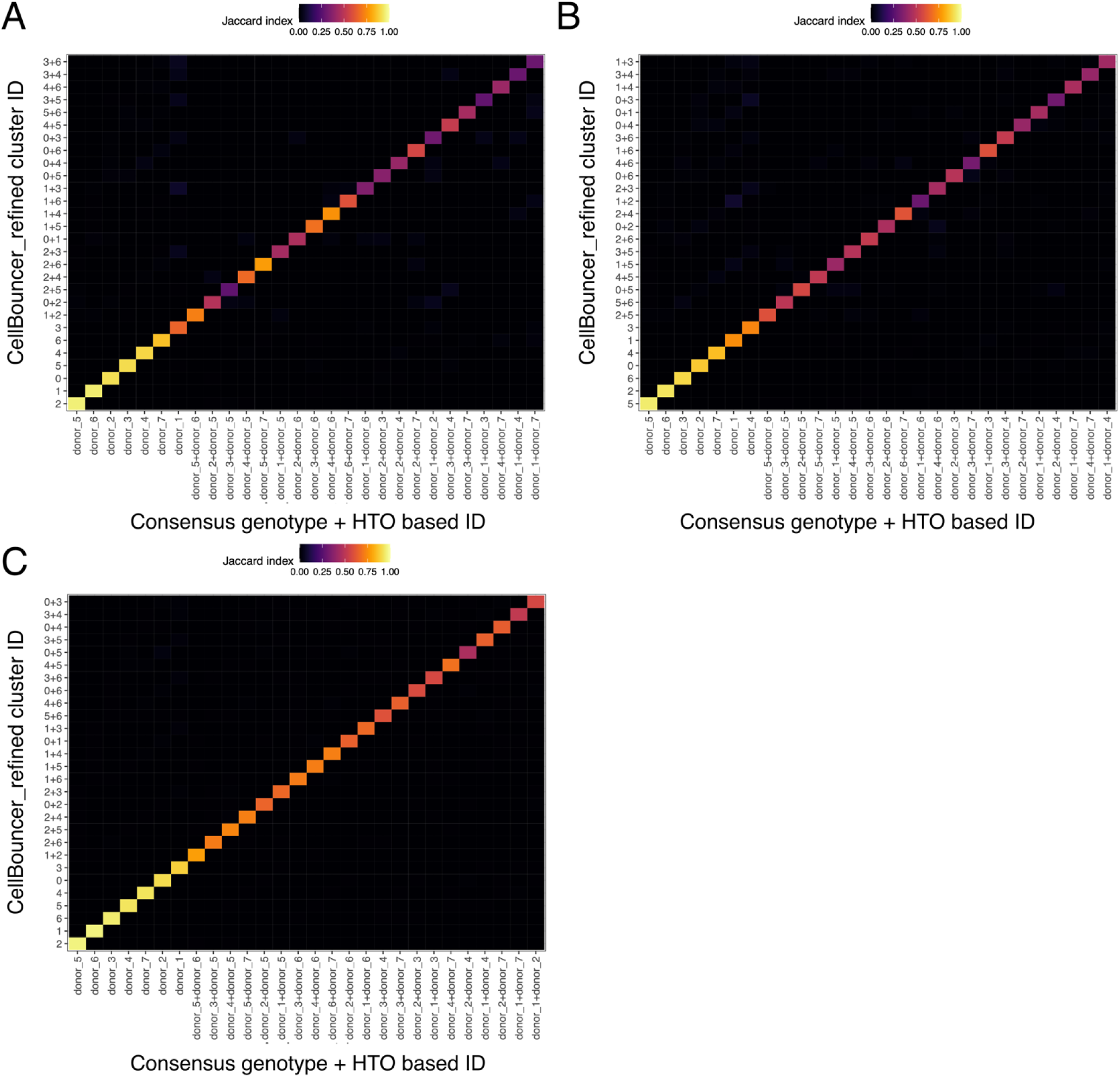
Comparison of cell labels from consensus genotype and CellPlex-based assignments (X-axis) vs inferred assignments without prior genotype data, using CellBouncer’s MT-to-VCF pipeline, followed by refine_vcf to re-infer genotypes given assignments, and another run of demux_vcf using the refined genotypes. Color denotes Jaccard similarity between the group of cells with the label on the X and the Y axis; doublet identities are named as component singlet identities separated by “+”. A: 2,000 cell downsampled data set. B: 5,000 cell downsampled data set. C: full data set (∼33,000 cells).

**Figure S3.**
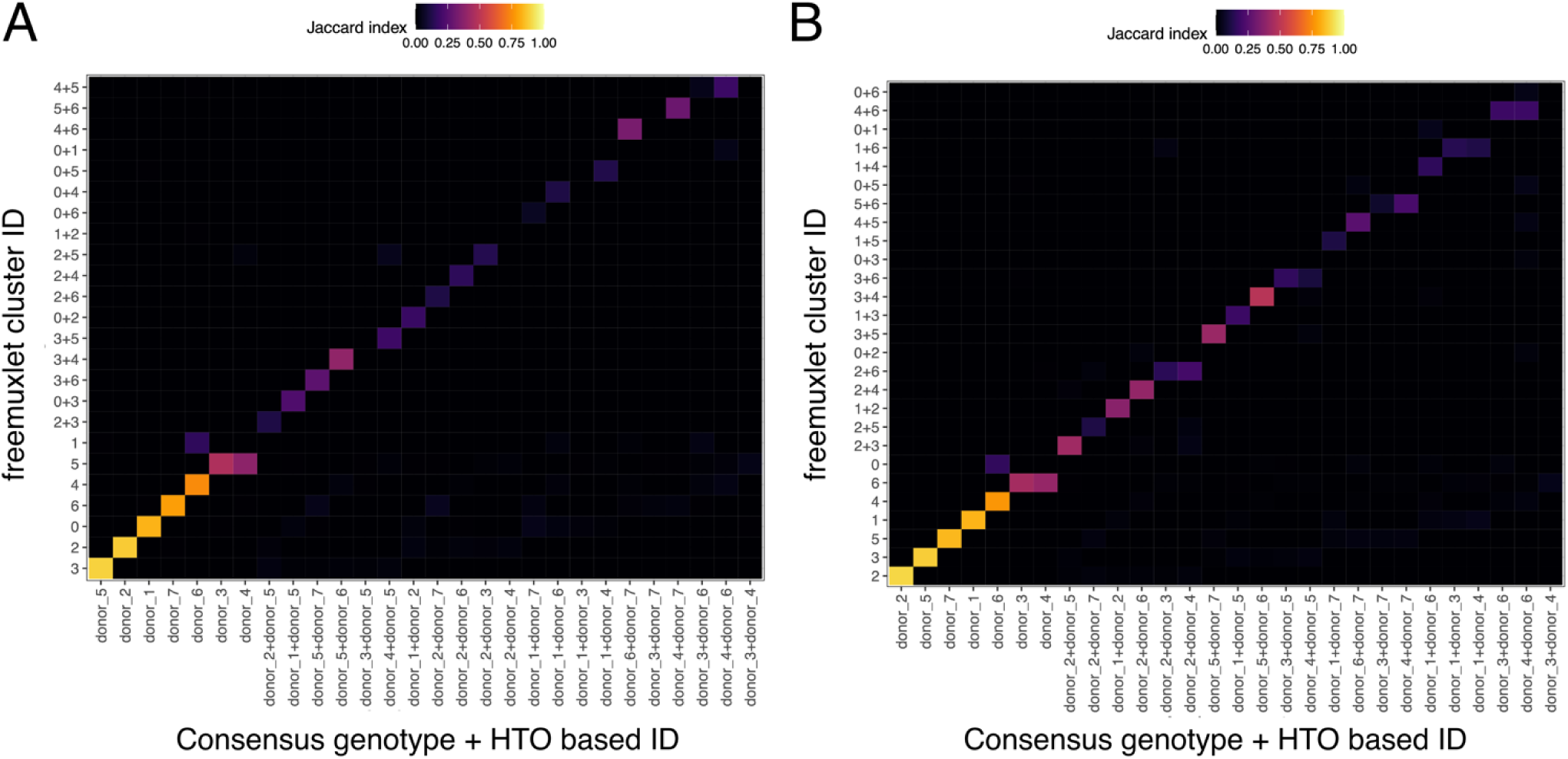
Comparison of cell labels from consensus genotype and CellPlex-based assignments (X-axis) vs inferred assignments without prior genotype data, using freemuxlet^19^. Color denotes Jaccard similarity between the group of cells with the label on the X and the Y axis; doublet identities are named as component singlet identities separated by “+”. A: 2,000 cell downsampled data set. B: 5,000 cell downsampled data set. Full data set is missing because the run did not complete.

**Figure S4.**
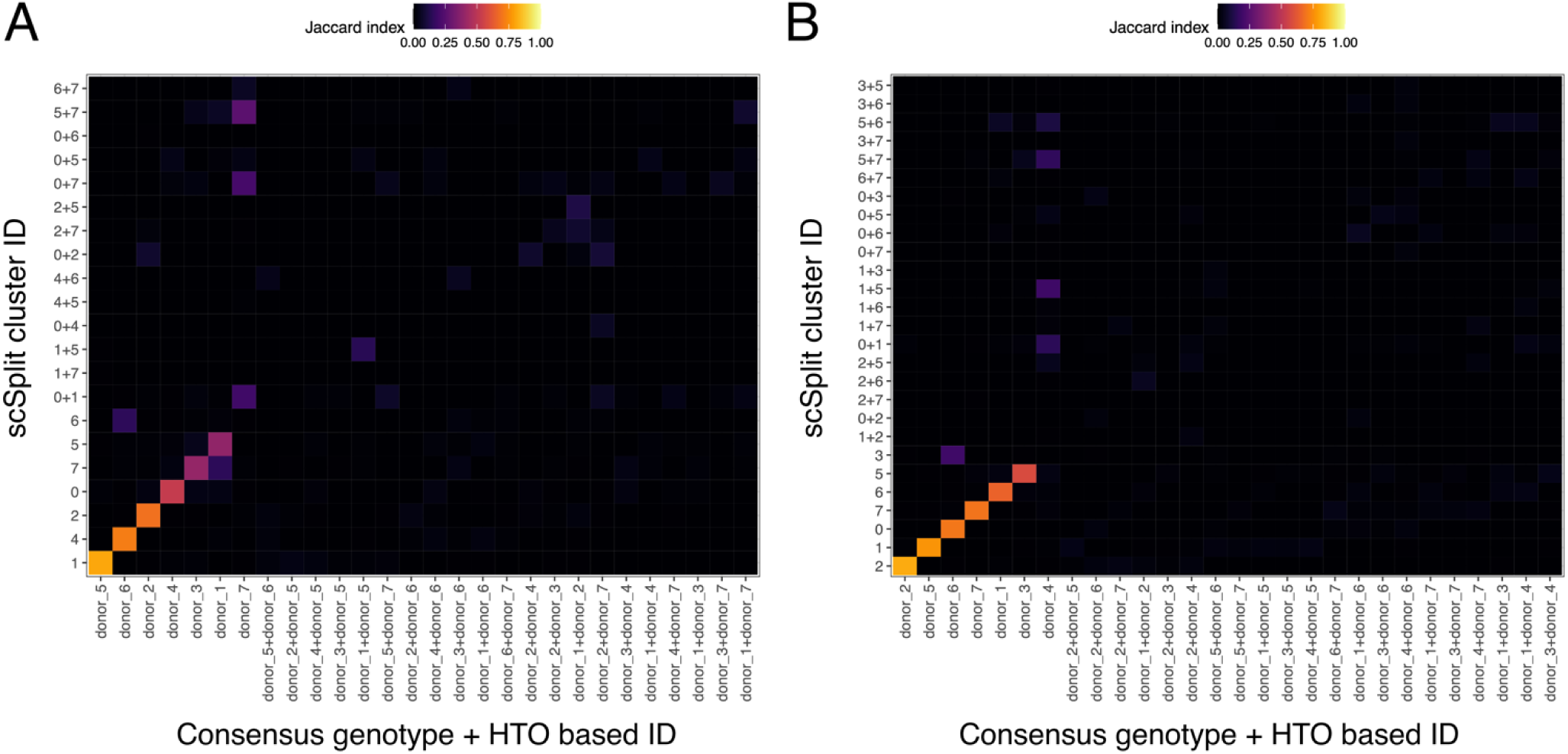
Comparison of cell labels from consensus genotype and CellPlex-based assignments (X-axis) vs inferred assignments without prior genotype data, using scSplit^18^. Color denotes Jaccard similarity between the group of cells with the label on the X and the Y axis; doublet identities are named as component singlet identities separated by “+”. A: 2,000 cell downsampled data set. B: 5,000 cell downsampled data set. Full data set is missing because the run did not complete.

**Figure S5.**
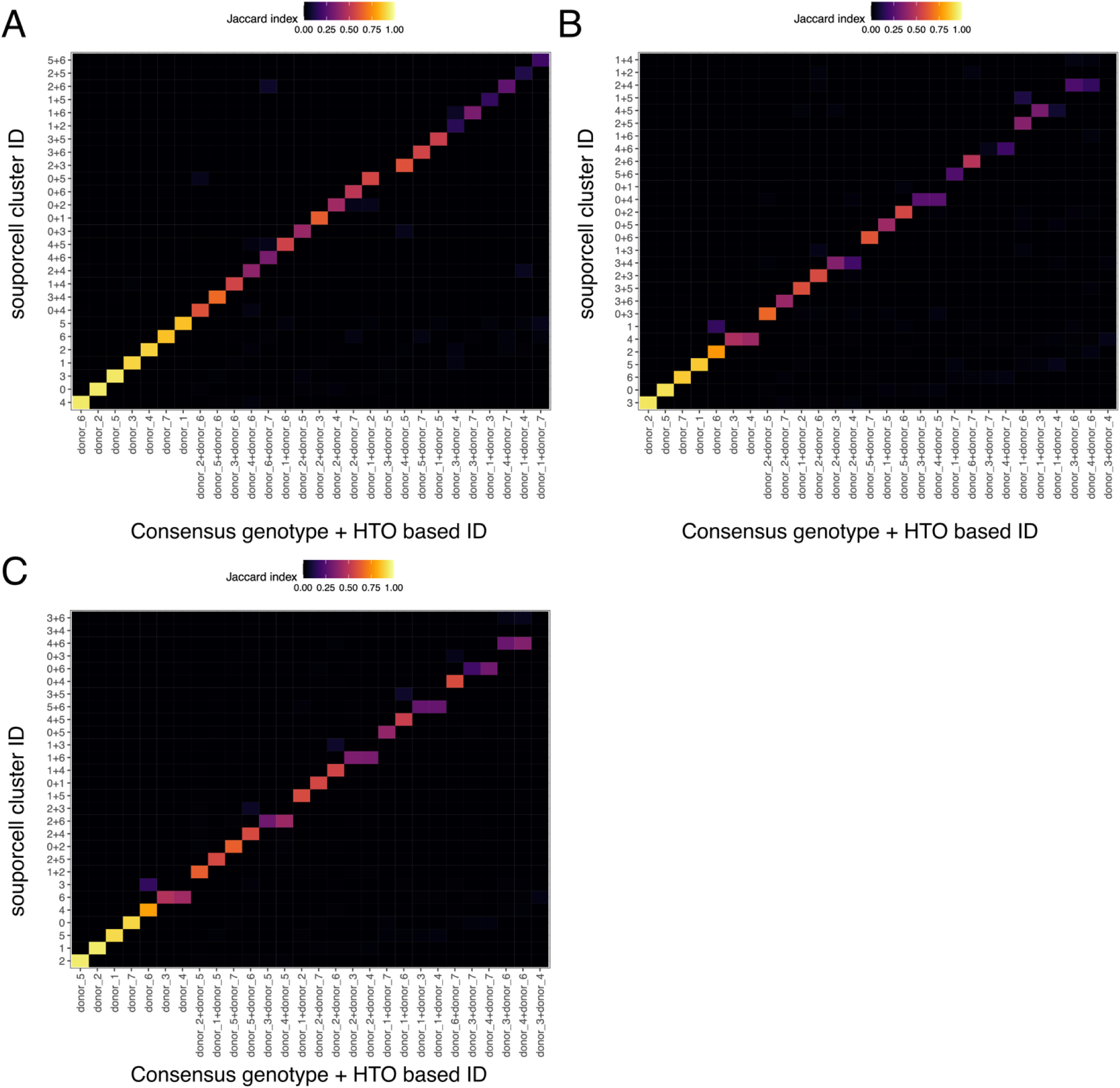
Comparison of cell labels from consensus genotype and CellPlex-based assignments (X-axis) vs inferred assignments without prior genotype data, using souporcell^17^. Color denotes Jaccard similarity between the group of cells with the label on the X and the Y axis; doublet identities are named as component singlet identities separated by “+”. A: 2,000 cell downsampled data set. B: 5,000 cell downsampled data set. C: full data set (∼33,000 cells).

**Figure S6.**
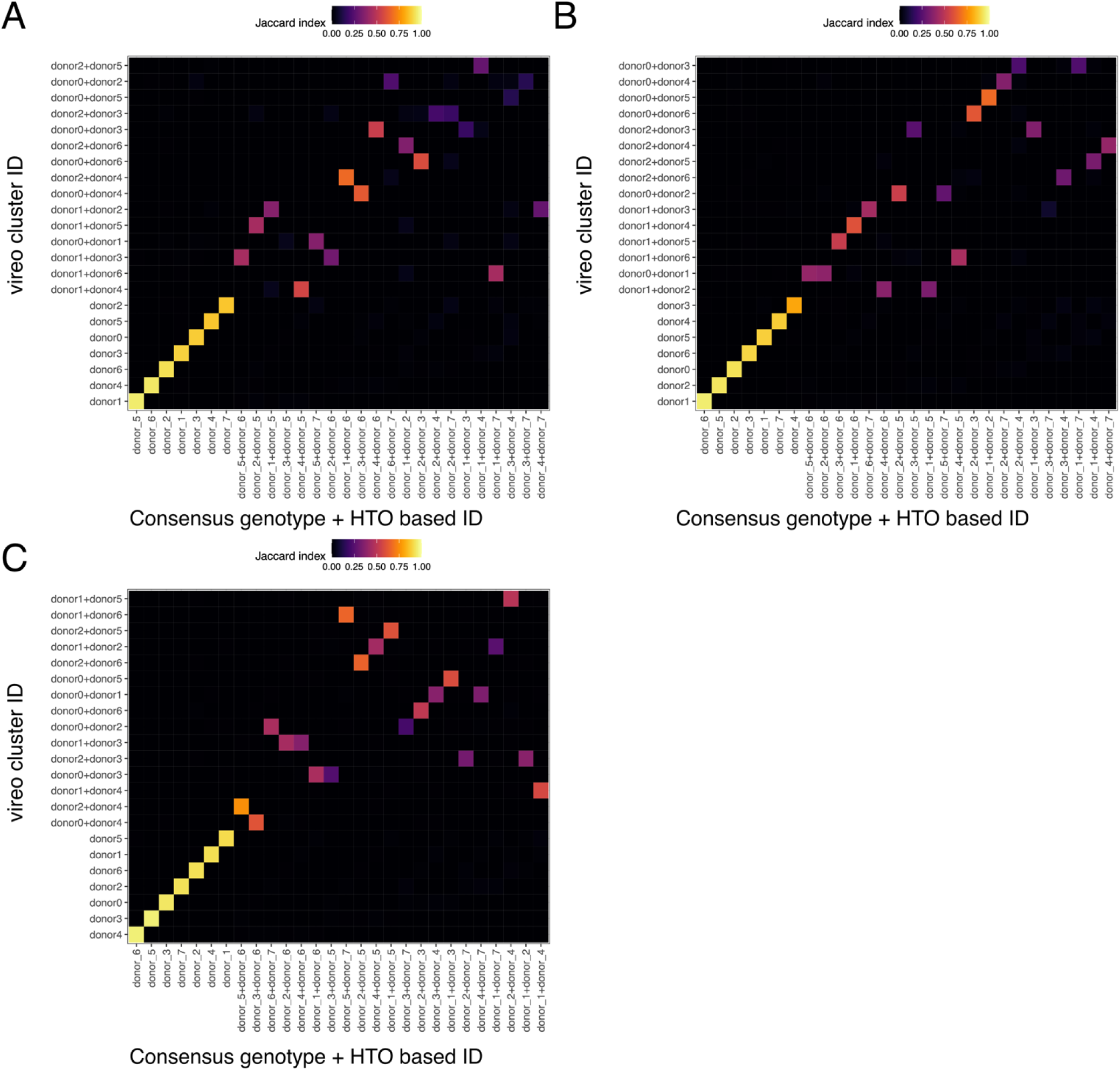
Comparison of cell labels from consensus genotype and CellPlex-based assignments (X-axis) vs inferred assignments without prior genotype data, using Vireo^15^. Color denotes Jaccard similarity between the group of cells with the label on the X and the Y axis; doublet identities are named as component singlet identities separated by “+”. A: 2,000 cell downsampled data set. B: 5,000 cell downsampled data set. C: full data set (∼33,000 cells).

**Figure S7.**
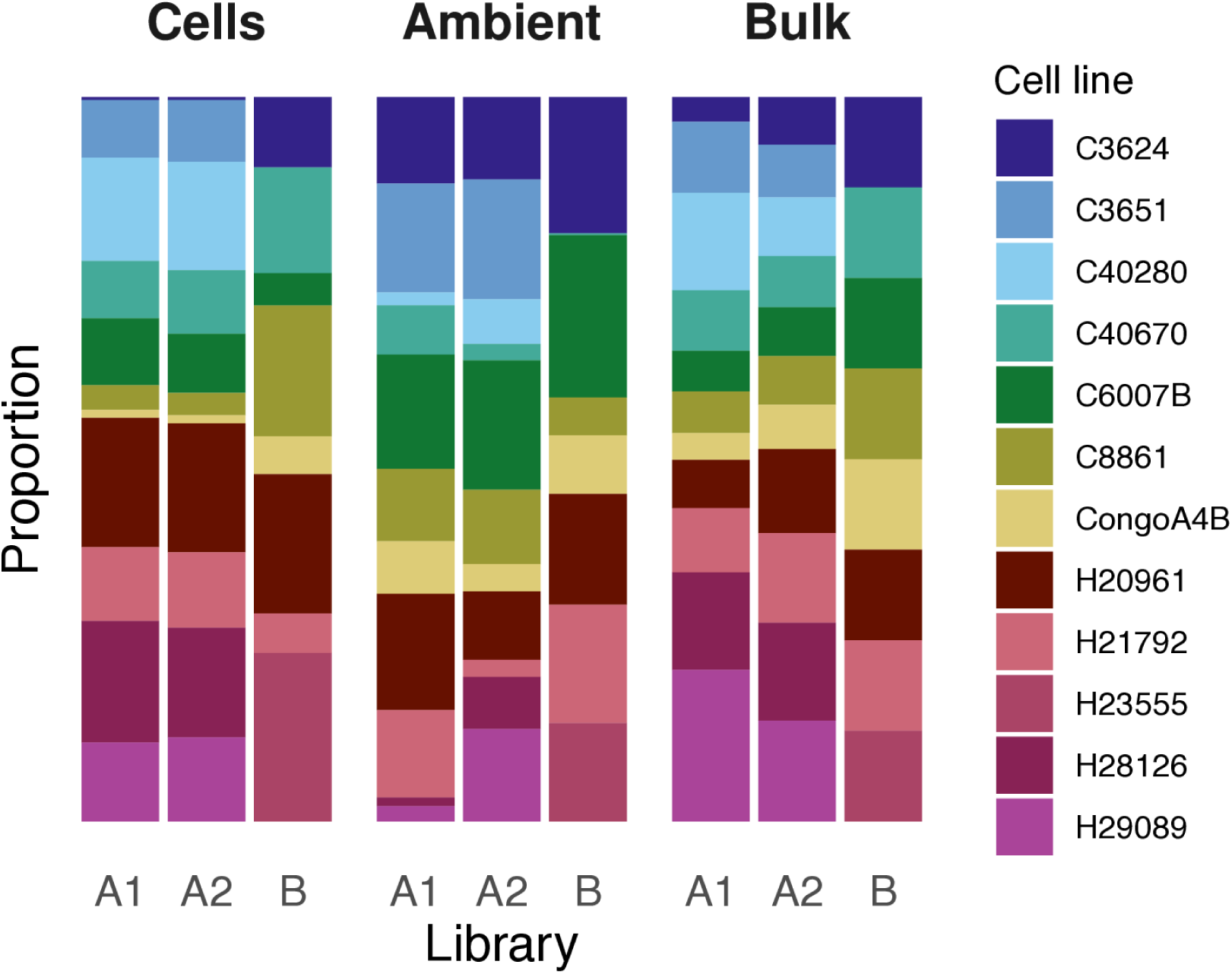
Proportions of total RNA from each contributor cell line from cell assignments using demux_vcf (left panel), proportion of ambient RNA inferred to originate from each contributor cell line (middle panel), and proportion of bulk RNA inferred to come from each contributor cell line (right panel), in three single cell RNA-seq libraries produced from tetraploid composite cell lines. A1 and A2 are technical replicates of the same library, and cell lines beginning “C” are chimpanzee lines (with the exception of CongoA4B, a bonobo cell line), and cell lines beginning “H” are human cell lines.

**Figure S8.**
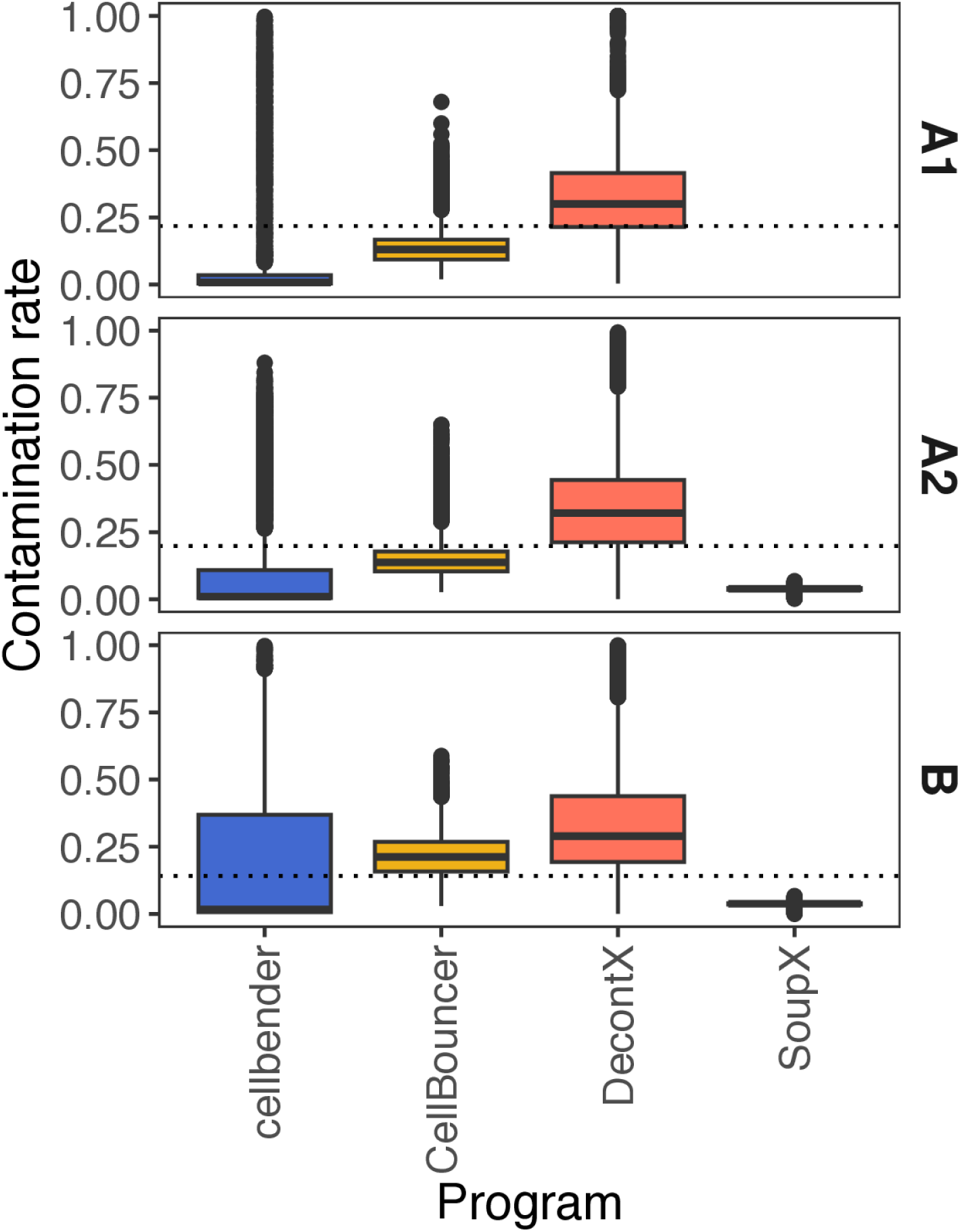
Comparison of inferred per-cell contamination rates using four programs (CellBouncer, cellbender, DecontX, and SoupX), on three single cell RNA-seq libraries from tetraploid composite cell lines. The dotted lines are estimates produced by comparing inferred bulk proportions to the proportions of cells assigned to each identity using demux_vcf (see Online Methods). Panels are libraries; A1 and A2 are replicates of the same library. SoupX was unable to run on library A1.

**Figure S9.**
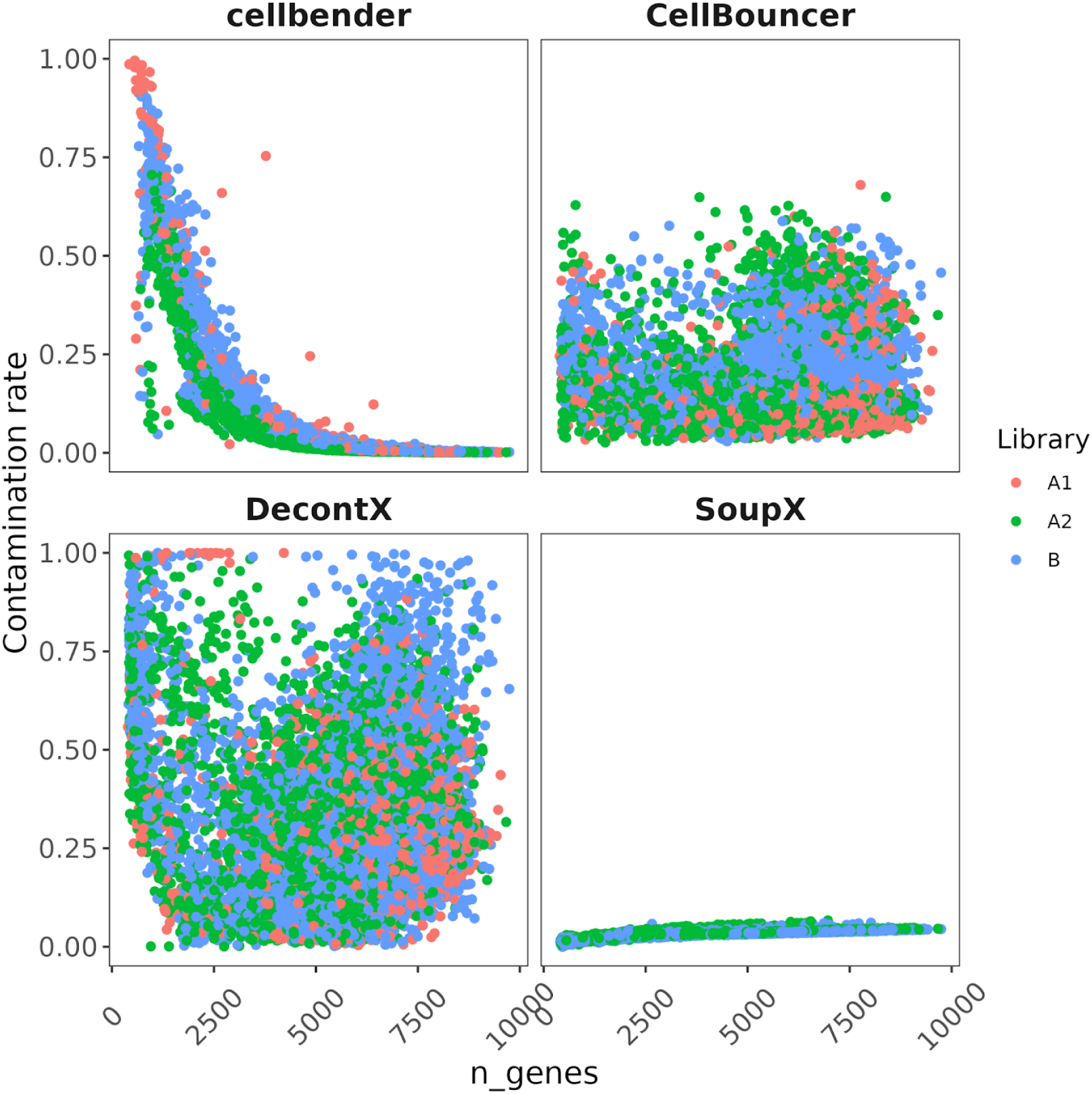
Comparison of per-cell contamination rate estimates inferred using four programs (cellbender, CellBouncer, DecontX, and SoupX) to the number of genes detected per cell in three single-cell RNA-seq libraries produced from tetraploid composite cell lines. A1 and A2 are technical replicates of the same library. SoupX was unable to run on library A1.

**Figure S10.**
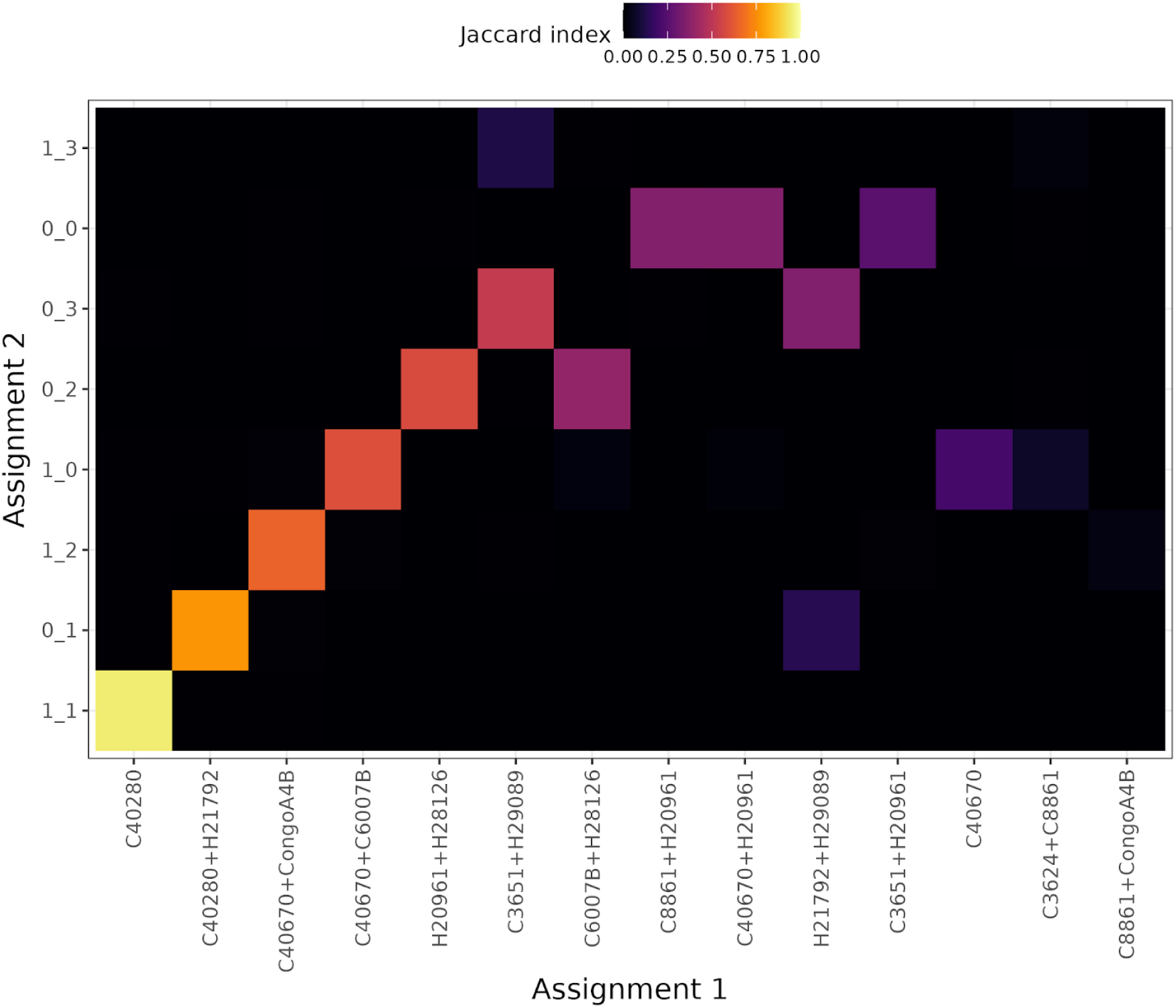
Genotype-based identities from demux_vcf (X-axis) versus inferred mitochondrial haplotypes using demux_mt with one round of subclustering (Y-axis) for tetraploid composite RNA-seq library A1. Subcluster identities are numeric IDs from each round of subclustering, separated by underscores. Colors denote Jaccard index of overlap between the set of cells belonging to the identity on the X and the identity on the Y axis.

**Figure S11.**
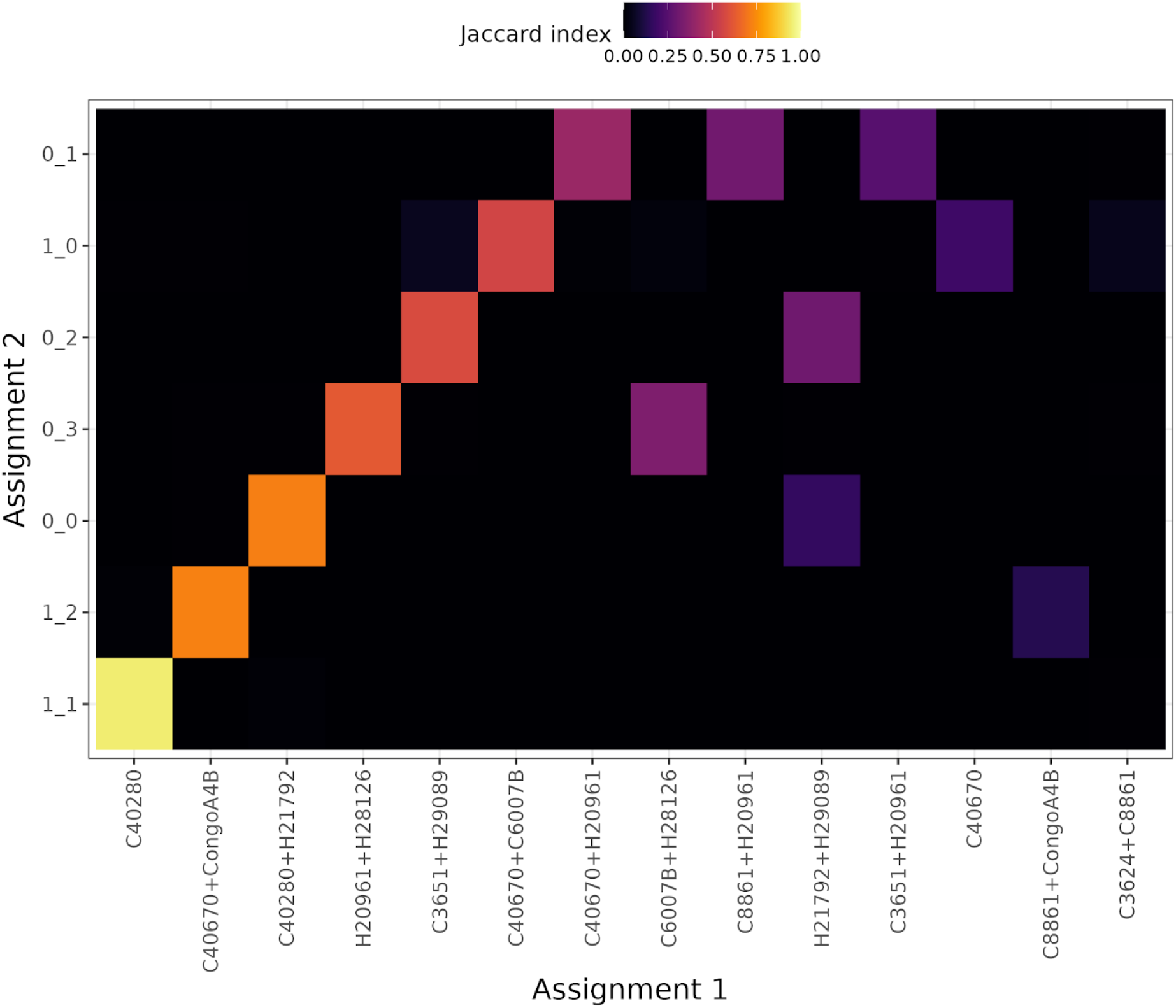
Genotype-based identities from demux_vcf (X-axis) versus inferred mitochondrial haplotypes using demux_mt with one round of subclustering (Y-axis) for tetraploid composite RNA-seq library A2. Subcluster identities are numeric IDs from each round of subclustering, separated by underscores. Colors denote Jaccard index of overlap between the set of cells belonging to the identity on the X and the identity on the Y axis.

**Figure S12.**
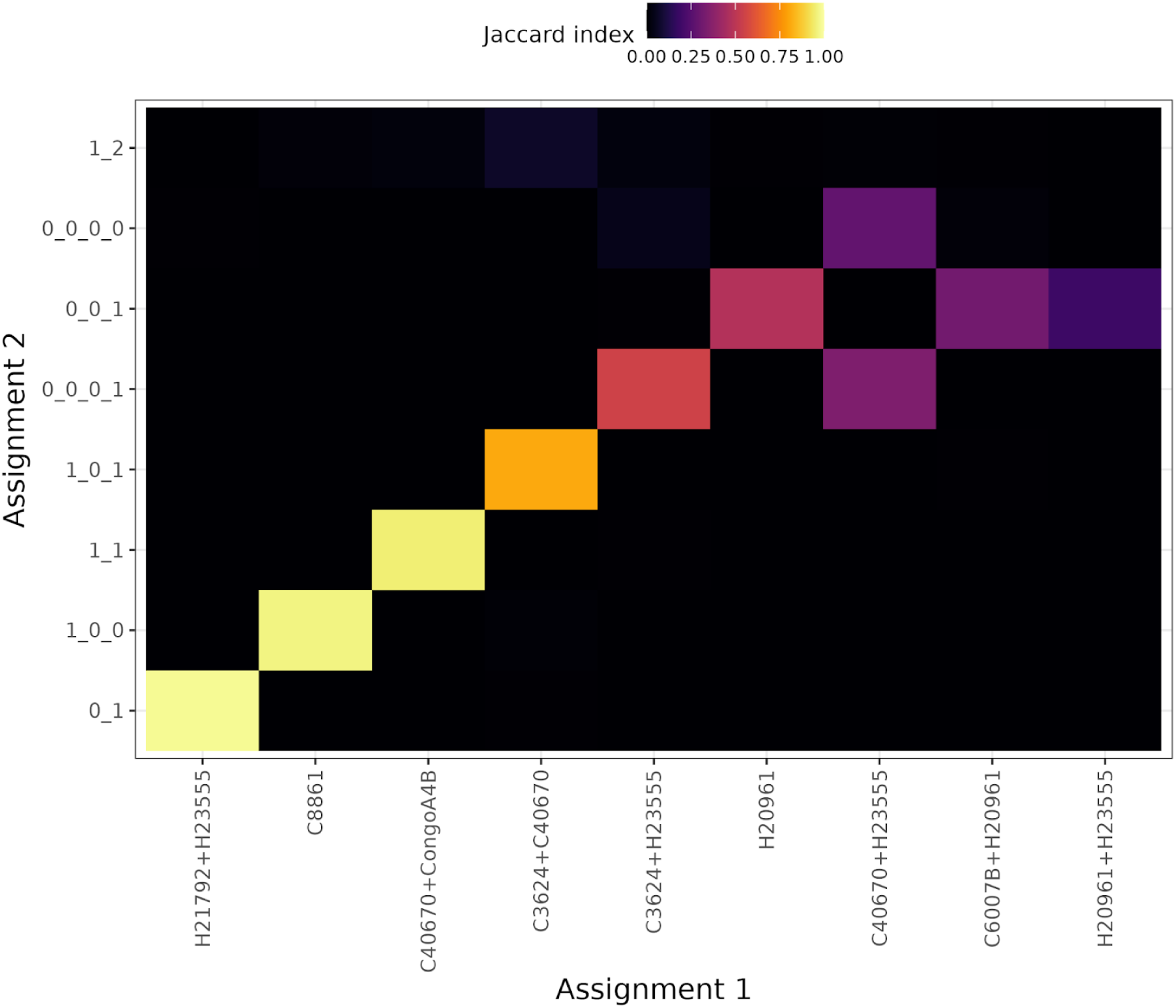
Genotype-based identities from demux_vcf (X-axis) versus inferred mitochondrial haplotypes using demux_mt with three rounds of subclustering (Y-axis) for tetraploid composite RNA-seq library B. Subcluster identities are numeric IDs from each round of subclustering, separated by underscores. Colors denote Jaccard index of overlap between the set of cells belonging to the identity on the X and the identity on the Y axis.

**Figure S13.**
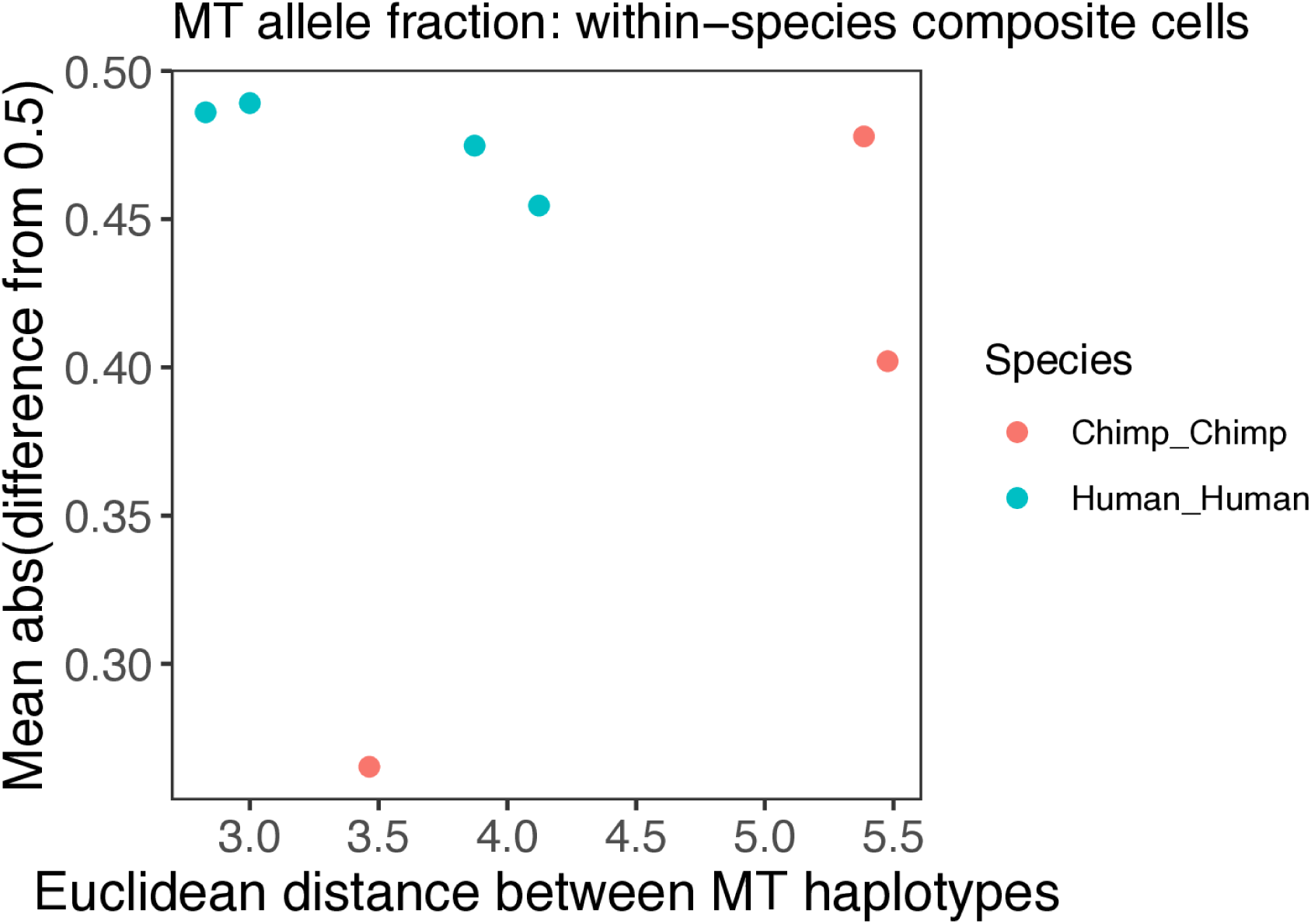
Relationship between the similarity of mitochondrial haplotypes of contributor lines and ratios of read counts matching each contributor line’s mitochondrial haplotype in within-species composite cells. X-axis: Euclidean distance between the two mitochondrial haplotypes in a composite cell line (computed using the dist function in R after encoding each mitochondrial haplotype as a bit string, with the ancestral allele = 0 and derived allele = 1 at each SNP). Y-axis: mean divergence in mitochondrial read fraction from expectation under equal proportions of each mitochondrion (0.5), in composite cells. Read fractions are count1/(count1+count2), where count1 is the number of reads matching contributor line 1’s mitochondrial haplotype and count2 is the number of reads matching contributor line 2’s mitochondrial haplotype, only considering SNPs that segregate between the two contributor mitochondrial haplotypes. This plot shows that species differences in the number of cells retaining mitochondria from both contributor lines are unlikely due to lower numbers of SNPs segregating between human than chimpanzee mitochondrial haplotypes, and therefore unlikely to be a technical artifact.

**Figure S14.**
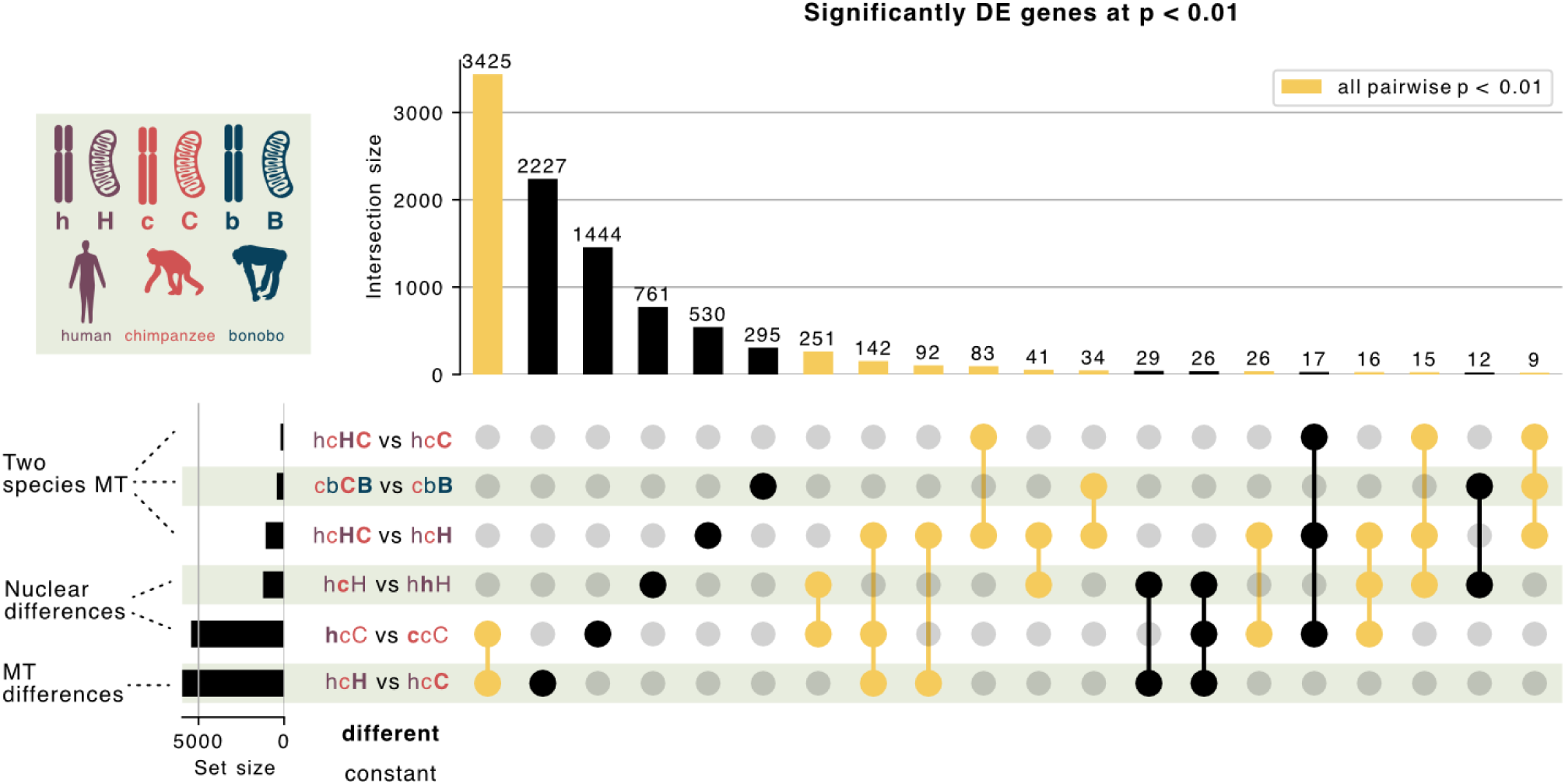
UpSet plot^109^ showing overlap of sets of differentially expressed genes between different classes of cells. For each comparison, cells with p < 0.01 of differential expression were selected, and the set of significantly DE genes for each cell class was tested for overlap with all others. Sets were highlighted in yellow when all pairwise comparisons between member gene sets were significant (p < 0.01) according to the hypergeometric CDF. Each cell class is named according to the legend; bold text highlights the difference in each comparison. This plot shows significant concordance between responses to having two species’ mitochondria and hints that the human genome may mount a larger transcriptional response to the presence of chimpanzee mitochondria than the chimpanzee genome’s response to the presence of human mitochondria.

**Figure S15.**
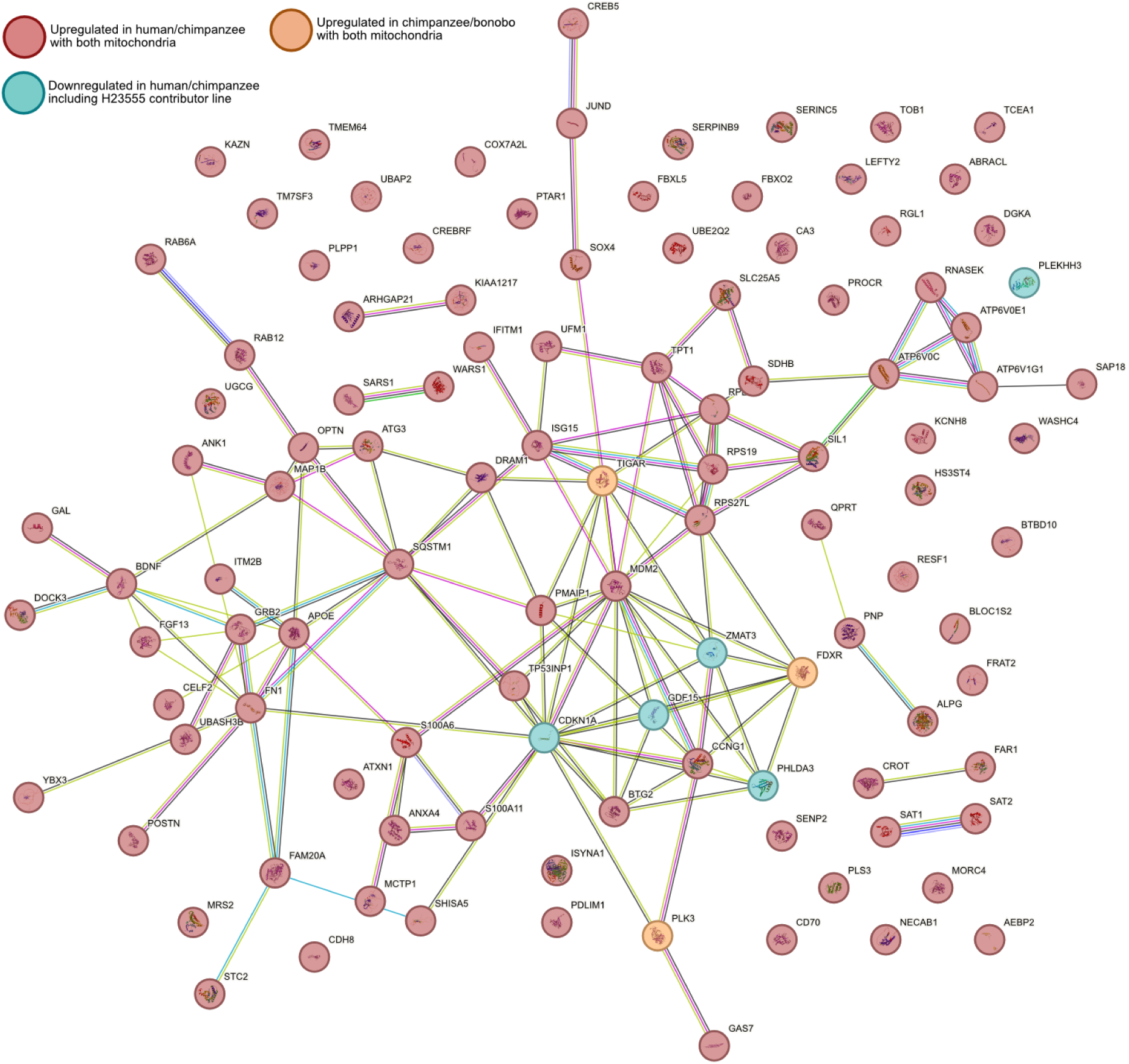
STRING110 network showing all genes significantly upregulated in response to having both human and chimpanzee mitochondria (p < 0.01, log_2_ fold change > 0 comparing human/chimpanzee cells with both species’ mitochondria to both human/chimpanzee cells with human mitochondria and human/chimpanzee cells with chimpanzee mitochondria). Protein-protein interactions (p = 4.11 x 10^-11^), and genes involved in p53 signaling (p = 0.022) are enriched in this set. Orange genes are also upregulated (p < 0.01) in chimpanzee/bonobo cells with both species’ mitochondria relative to those with only bonobo mitochondria (enriched for overlap with this set, p = 1.34 x 10^-8^). Teal genes were downregulated (p < 0.01) in human/chimpanzee cells from contributor line H23555, relative to other human/chimpanzee cells (enriched for overlap with this set, p ≈ 0). This plot reveals a key part of the regulatory network involved in p53 signaling that is activated when two species’ mitochondria are present, and deactivated by cells capable of surviving with two species’ mitochondria.

**Figure S16.**
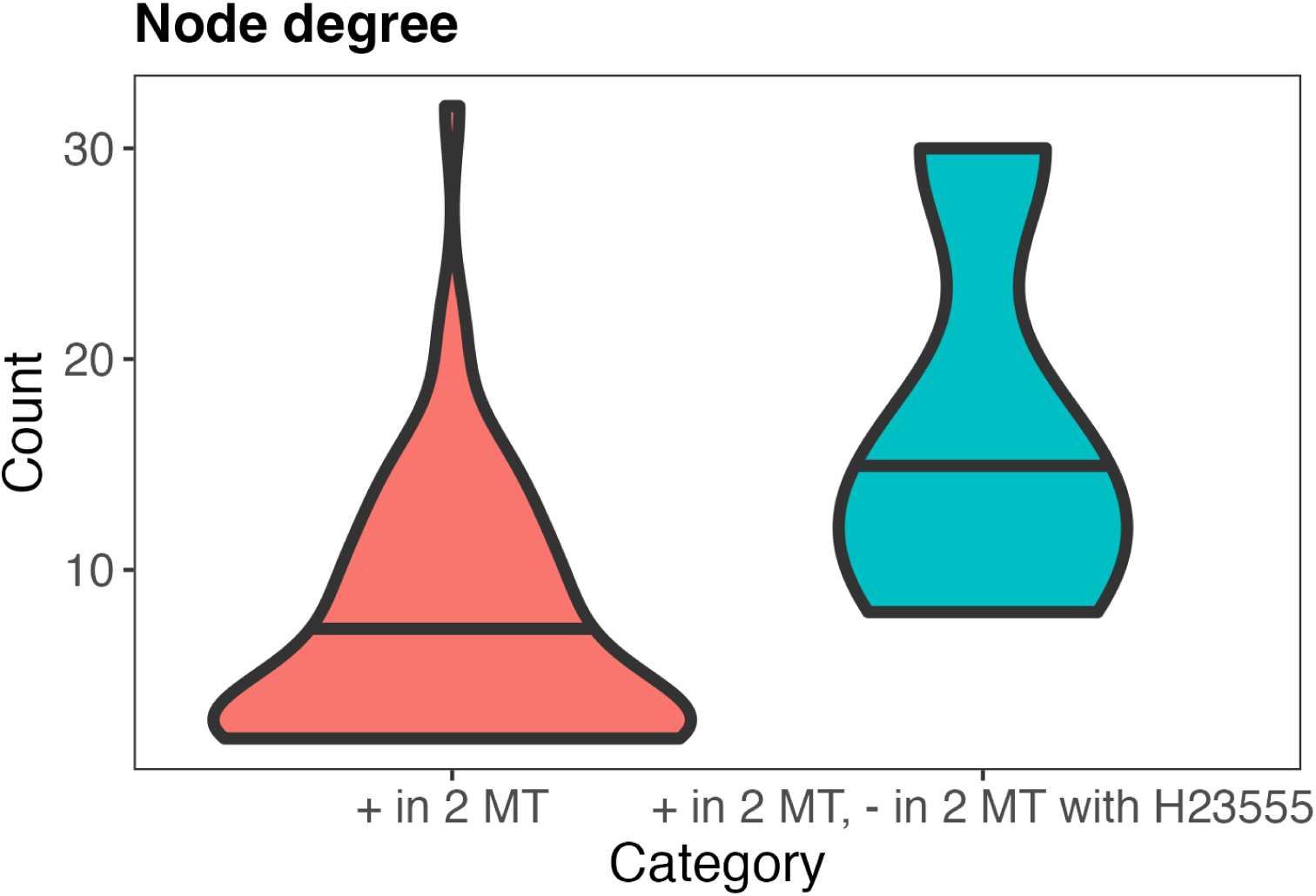
Number of edges (degree or connectivity) of nodes in Figure S15, colored by whether or not they were downregulated in cells derived from the contributor line H23555. Genes downregulated in human/chimpanzee cells derived from contributor line H23555, relative to those that did not derive from this line, not only downregulated a significant number of genes upregulated in human/chimpanzee cells with two species’ mitochondria, but the downregulated genes tended to be higher-connectivity “hub” genes (Kolmogorov-Smirnov p = 0.05392). This suggests that these genes may be key regulators in the transcriptional response to the presence of two species’ mitochondria.

**Figure S17.**
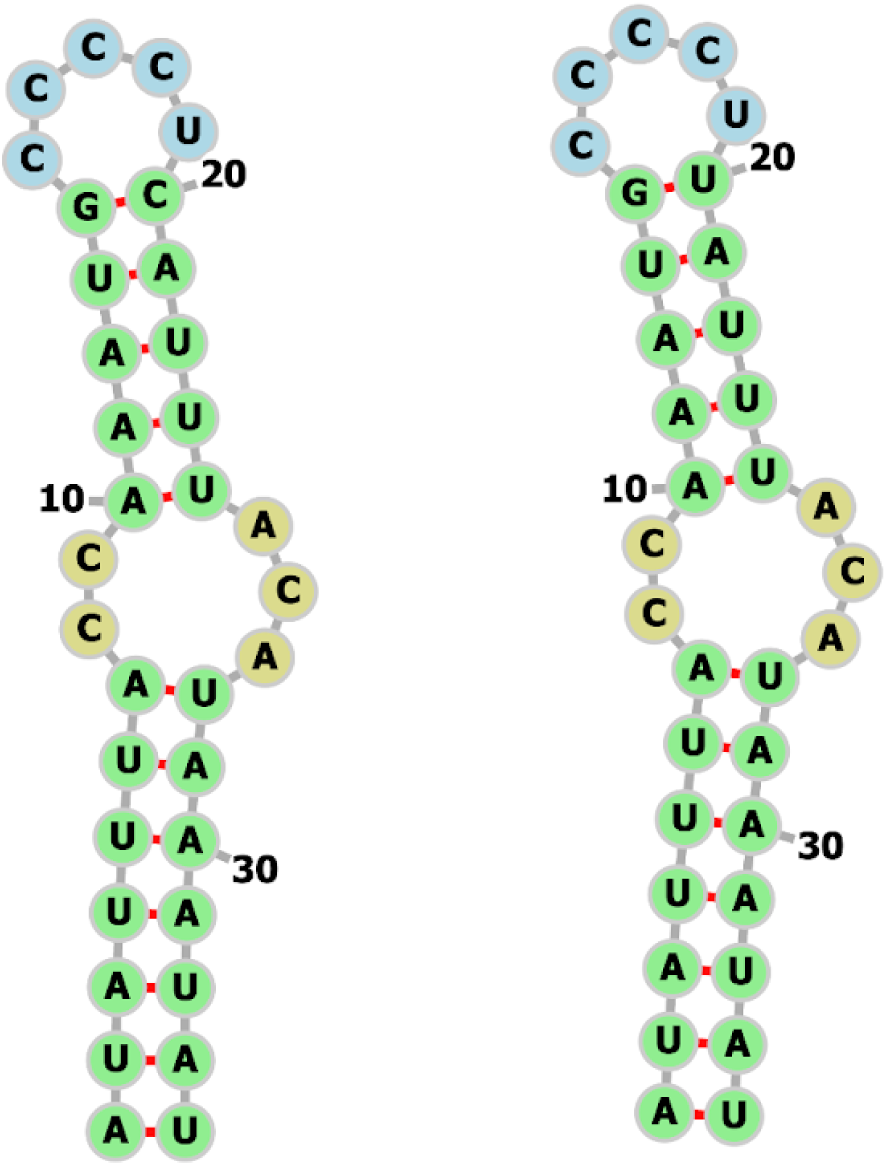
Predicted hairpin-loop secondary structure from RNAfold^111^ for the region around fixed the human-specific derived T-C mutation at chrM:10479 (hg38). This region (chrM:10459-10492 inclusive) was chosen by predicting structure for a broader region and selecting bases involved in the hairpin structure around the focal site. The mutation of interest is labeled as base position 20 in both structures. Left: human reference sequence; Right: human (hg38) reference sequence with the base back-mutated to the ancestral allele. Predicted base pairing probability between base 14 and 20 in the human (left) structure is 0.948, while the same probability with the ancestral allele is 0.611.

